# Deconvolution of Single Homeologous Polymorphism (SHP) drives phylogenetic analysis of allopolyploids

**DOI:** 10.1101/2025.07.17.665301

**Authors:** R. Sancho, P. Catalán, J.P. Vogel, B. Contreras-Moreira

## Abstract

The genomic and evolutionary study of allopolyploid organisms involves multiple copies of homeologous chromosomes, making their assembly, annotation, and phylogenetic analysis challenging. Bioinformatics tools and protocols have been developed to study polyploid genomes, but sometimes require the assembly of their genomes, or at least the genes, limiting their use. We have developed AlloSHP, a command-line tool for detecting and extracting single homeologous polymorphisms (SHPs) from the subgenomes of allopolyploid species. This tool integrates three main algorithms, WGA, VCF2ALIGNMENT and VCF2SYNTENY, and allows the detection SHPs for the study of diploid-polyploid complexes with available diploid progenitor genomes, without assembling and annotating the genomes of the allopolyploids under study. AlloSHP has been validated on three diploid-polyploid plant complexes, *Brachypodium*, *Brassica*, and *Triticum*-*Aegilops*, and a set of synthetic hybrid yeasts and their progenitors of the genus *Saccharomyces*. The results and congruent phylogenies obtained from the four datasets demonstrate the potential of AlloSHP for the evolutionary analysis of allopolyploids with a wide range of ploidy and genome sizes.

AlloSHP combines the strategies of simultaneous mapping against multiple reference genomes and syntenic alignment of these genomes to call SHPs, using as input data a single VCF file and the reference genomes of the known or closest extant diploid progenitor species. This novel approach provides a valuable tool for the evolutionary study of allopolyploid species, both at the interspecific and intraspecific levels, allowing the simultaneous analysis of a large number of accessions and avoiding the complex process of assembling polyploid genomes.

## Background

A polyploid is an organism with three or more complete sets of chromosomes. Two types of polyploids can be defined according to their origin; while autopolyploids result from non-reduced gametic crosses within or between populations of the same species, allopolyploids derive from crosses between different species, i.e. hybrids (Ramsey & Schemske, 1998). It is estimated that 30-70% of all angiosperms are polyploids (Levin & Donald A, 2002; Masterson, 1994; Stebbins, 1949; Wood et al., 2009). This range varies depending on the methodology and the taxonomic circumscription of the plants used to estimate it (Halabi et al., 2023). There is consensus that polyploidization is one of the key mechanisms of speciation and is ubiquitous among angiosperms (Ramsey & Schemske, 1998; Soltis et al., 2009). Indeed, it is accepted that all seed plants have undergone at least one round of whole genome duplication (WGD) in their evolutionary history and are considered to have a paleopolyploid ancestry (Y. Jiao et al., 2011; Renny-Byfield & Wendel, 2014). Although polyploidization events have played a key role in plant evolution, they have also occurred to a lesser extent in other organisms such as animals and fungi, resulting in polyploid species of fish, amphibians, reptiles, and, although extremely rare, also birds and mammals.

Polyploid species can be found among invertebrates such as crustaceans, insects, mollusks, annelids or nematodes, among others (Gregory & Mable B.K., 2005; Van de Peer & Meyer, 2005). In the case of fungi, some exist as stable haploid, diploid, or polyploid (heteroploid) cells or organisms, while others change ploidy under certain conditions (Albertin & Marullo, 2012; Todd et al., 2017). Rapid advances in both sequencing technologies and bioinformatics make it possible to generate and analyze vast amounts of sequencing data from a wide variety of organisms. This is also leading to a remarkable and continuous increase in the number of available genomes, allowing the analysis of species and populations at the pan-genomic level, i.e. using multiple reference genomes simultaneously [(Gordon et al. (2017), *Brachypodium distachyon*; Jayakodi et al. (2020), barley; W. B. Jiao & Schneeberger (2020), and Kang et al. (2023), *Arabidopsis*; Montenegro et al. (2017), wheat; Golicz et al. (2016), *Brassica oleracea*; among other species reviewed in Bayer et al. (2020), and Schreiber et al. (2024)]. Although sequencing technologies and bioinformatics tools for assembly are advancing at a dizzying pace, genome assembly and evolutionary studies of allopolyploid organisms remain challenging for possessing multiple sets of chromosomes from their homeologous genomes. (Kong et al., 2023; K. Li et al., 2023; Mason, 2015; Soltis et al., 2016; Y. Wang et al., 2023).

Several approaches have been developed for phylogenetic studies of polyploid species. However, many of them focus on coding regions (genes), which on the one hand requires prior assembly and annotation of their genes, and on the other hand leaves out the intergenic regions, which contain many informative loci that can help infer the evolutionary relationships of populations of diploid-polyploid complexes. Thus, Bombarely et al. (2014) used the consensus diploid transcriptome to identify homeologous SNPs in the genus *Glycine* and then built a progenitor reference set for each polyploid species joining the progenitors’ diploid transcriptome sets. These references were used to separate reads according to their preferential mapping to one or the other progenitor genome.

Oxelman et al. (2017) reviewed the explicit species network methods, such as the permutation approach (e.g., PhyloNet; Than et al., 2008; Than & Nakhleh, 2009) and simultaneous gene tree and species network inference (AlloppNet; Jones et al., 2013), used to infer the allopolyploid origins of multiple genera. Marcussen et al. (2015) constructed a dated allopolyploid network from individual gene trees of the genus *Viola* using three low-copy nuclear genes (GPI, NRPD2a, and SDH). Kamneva et al. (2017) studied the phylogeny of several diploid and polyploid species of the genus *Fragaria* using a large number of multilabeled gene trees, and Sancho et al. (2022) developed a protocol (PhyloSD) to infer the homeologous subgenomes in *Brachypodium* allopolyploids and reconstruct their evolution, even when their diploid ancestors are unknown (orphan subgenomes).

Other approaches have been developed to analyze interspecific hybrids and allopolyploids using SNPs or synteny and microsynteny-based approaches. However, these two strategies have not been combined to reconcile syntenic homeologous SNPs in a single alignment that allows phylogenetic inference and studies of population structure at the subgenomic level. Regarding SNP discovery, some approaches use the genomes of their extant closest available progenitor genomes to infer the SNPs from each of the homeologous subgenomes of the allopolyploid, defined as homeoSNPs. For example, Page et al. (2013) implemented the PolyCat pipeline to map and categorize the genomic data generated from allopolyploids and it was tested in cotton. Peralta et al. (2013) developed SNiPloid, a web tool focused on SNP analysis of RNA-Seq data obtained from allotetraploids. Mithani et al. (2013) and Khan et al. (2016) implemented HANDS and HANDS2, a method to characterize homeolog-specific polymorphisms (HSPs) in polyploid genomes, tested in the allopolyploids bread wheat and *Brassica* genus. Kulkarni et al. (2023) developed the Comprehensive Allopolyploid Genotyper (CAPG) method, which uses a likelihood to weight read alignments against both subgenomic references and then genotype individual allopolyploids from whole-genome resequencing data. Page & Udall (2015) and Phillips (2024) have published reviews of the various methods for mapping and categorizing reads, as well as the variant calling approaches required for the study of polyploid organisms.

Although synteny and microsynteny-based approaches have been widely used in phylogenetic inference, they have mainly studied diploid species and have focused on the synteny of coding regions, i.e., the collinearity of genes (Walden & Schranz, 2023, Brassica; Liu et al., 2023, rosids; Zhao et al., 2021, angiosperms), without exploiting the phylogenetic information of the rest of the genome.

Similarly, available approaches to reveal the homeologous subgenomes of allopolyploid species have also focused on coding regions. These require genomes to be previously assembled and annotated, and sometimes prior information on the diploid species of particular group. Furthermore, these protocols can sometimes be very complex for users without prior bioinformatics knowledge.

To simplify the bioinformatics protocols used in the study of allopolyploids, as well as the type of sequences and prior information required, we have developed an integrative whole-genome synteny-based phylogenetic inference approach to detect syntenic homeoSNPs, defined here as Single Homeologous Polymorphisms (SHP), for the study of diploid-polyploid complexes for which diploid progenitor genomes are available. The AlloSHP pipeline does not require assembling nor annotating the genomes of the allopolyploids under study. Our approach allows phylo(sub)genomic analysis at both the inter- and intra-specific levels, providing insight into which lineages may be involved in the origin and evolution of the different polyploid populations that are part of diploid-polyploid complexes.

## Implementation

The input files required to use this protocol are i) the reference genomes of the diploid progenitor species whose syntenic positions are going to be determined (FASTA format, Figure 1A) and ii) a VCF summarizing the mapping of reads of all the polyploid samples to be studied (Figure 1B). The VCF is obtained by merging BAM files, one per sample, and must contain the DP (total read depth) field. The step-by-step instructions and the external software used are detailed in the following sections and supplementary information.

**Figure 1.**
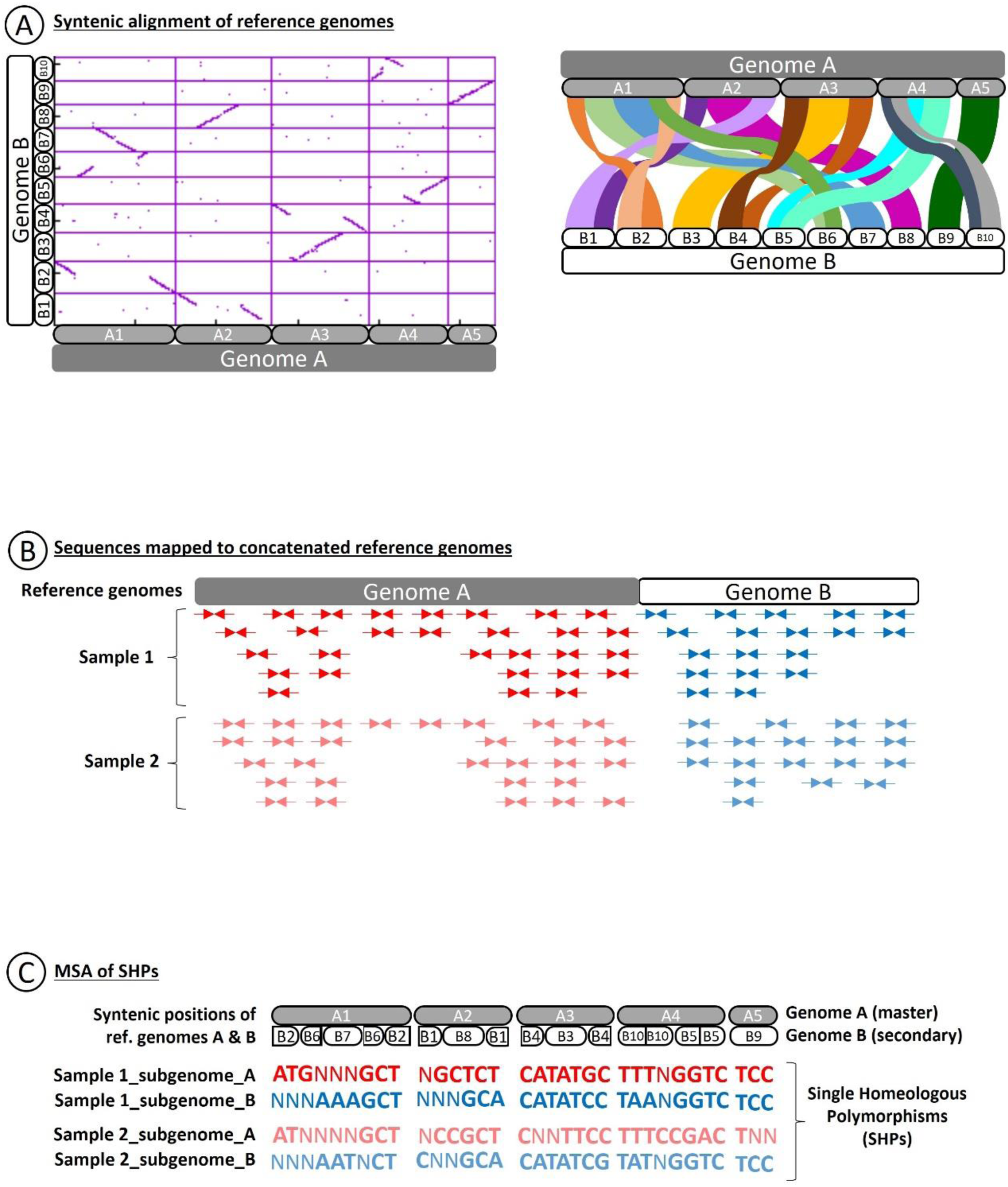
Simplified illustration of the main pipeline processes. **(A)** Alignment of syntenic regions between reference genomes A (chromosomes A1-A5) and B (chromosomes B1-B10) using CGaln (left) and displayed in a riparian plot (right). **(B)** Mapping of reads against the concatenated reference genomes A and B. Reads from allopolyploid samples (sample 1 and sample 2) are represented by arrows (forward and reverse). Red and blue colors indicate reads mapped against genome A or B, respectively. **(C)** Multiple sequence alignment of Single Homeologous Polymorphisms (SHPs). Each letter corresponds to an SHP mapped against genome A (red) or B (blue). Each sample (samples 1 and 2) has as many draft subgenomes as reference genomes used in the mapping.

The AlloSHP detection pipeline has three core algorithms, included as scripts in the repository https://github.com/eead-csic-compbio/AlloSHP: WGA (Whole Genome Alignment), VCF2ALIGNMENT, and VCF2SYNTENY. Each algorithm plays a key role in extracting the syntenic positions between the reference genomes, determining the most confident SNPs according to sequencing depth and missing sample thresholds, and finally combining both types of information to deconvolute and extract SHPs (Figure 1; Figure 2).

**Figure 2.**
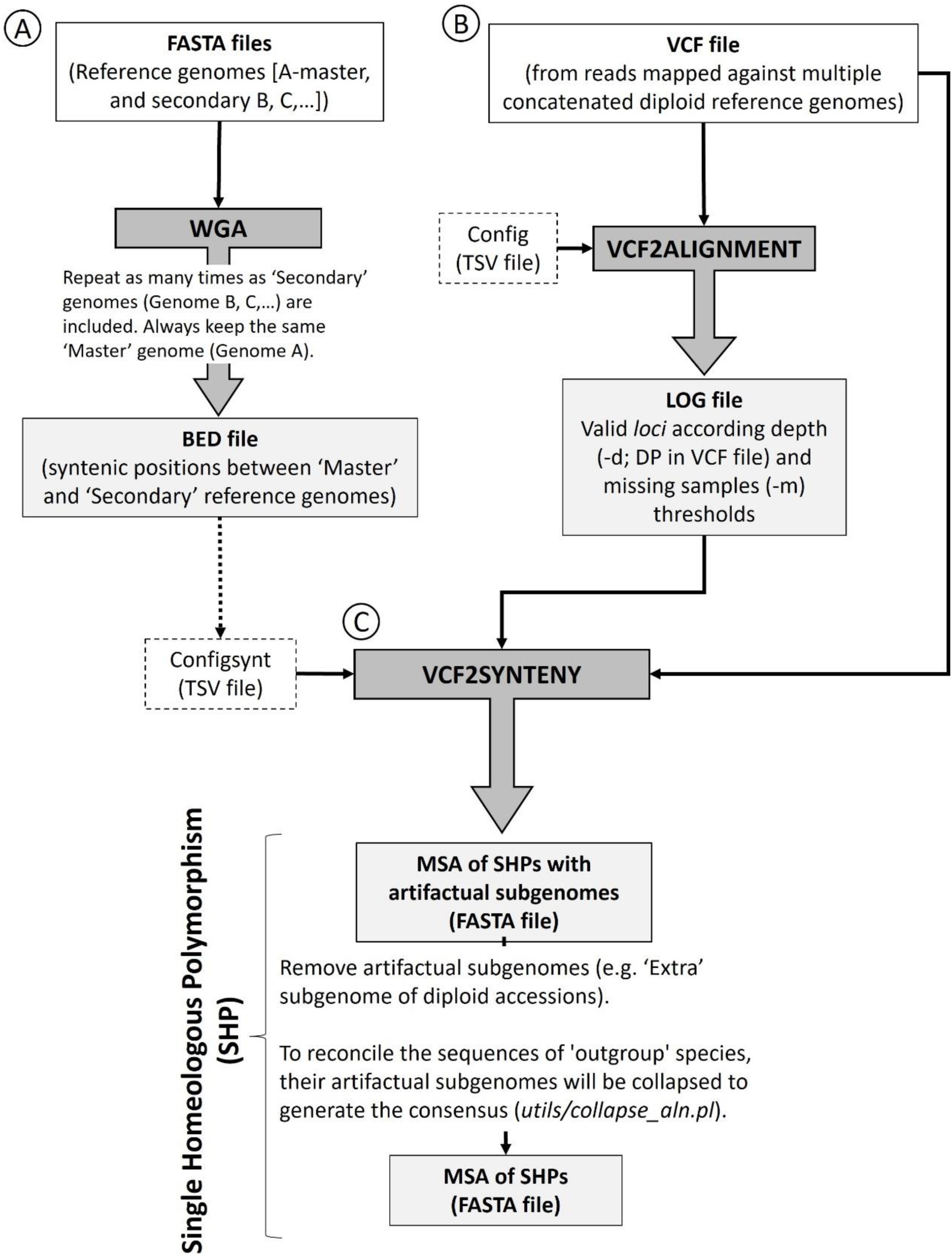
Flowchart of the main tasks and deliverables of the AlloSHP pipeline. The gray squares indicate the three main algorithms (WGA **(A)**, VCF2ALIGNMENT **(B)**, and VCF2SYNTENY**(C)**) that make up the pipeline. The white squares indicate the input files required for each algorithm and the resulting output files. The dashed boxes indicate the two configuration files needed for the VCF2ALIGNMENT and VCF2SYNTENY algorithms.

The ***Whole Genome Alignment (WGA)*** algorithm by default calls the aligner CGaln (Nakato & Gotoh, 2010) to perform the alignment and detect the syntenic segments between the reference genomes after soft-masking repeated sequences (Figure 1A; 2A). One progenitor reference genome is defined as "primary or master (genome A)" and the others as "secondary (genomes B, C, …)". There can be any number of “secondary” progenitor reference genomes according to the ploidy of the allopolyploids under study, but they must always be aligned against the same master reference genome [e.g. the study of an allohexaploid includes three reference genomes from its diploid progenitors. One genome is established as the "master (A)," while the other two are established as "secondary (B and C)"]. The syntenic positions are extracted and used downstream. Three main output files are generated: (i) a 0-based BED list of syntenic positions indicating chromosome, position, strand, nucleotide, CGaln syntenic block, and SNP presence between primary and secondary reference genomes; (ii) a Log file containing the parameters, thresholds, and additional information extracted from the CGaln output; and (iii) a PDF dot plot file showing the syntenic regions between reference genomes. This needs to be inspected to assess the quality of the alignment in terms of matched regions and noise.

The ***VCF2ALIGNMENT*** algorithm parses an input VCF (variant call format) file and produces a list of valid homozygous positions (valid loci), taking into account the minimum read depth (-d) and maximum missing samples (-m) thresholds. Heterozygous sites are handled as missing data to avoid detecting allelic SNPs as homeologous SNPs (Kulkarni et al., 2023). The result is a LOG file with a list of positions (valid loci) that have passed the thresholds. This list indicates the chromosome and position with respect to the master reference genome, as well as the number of missing samples and the called nucleotide (ATGC). In addition, at the end of the file, the total number of valid loci and polymorphic loci is shown, as well as for each target sample and per reference chromosome. Optionally, multiple sequence alignments (MSA) can be calculated, but on this protocol, this only really makes sense on the next step (Figure 1B; 2B).

The ***VCF2SYNTENY*** algorithm parses the i) VCF file obtained from reads mapped to multiple concatenated reference genomes, ii) the LOG file of valid loci computed by VCF2ALIGNMENT, and iii) the synteny-based equivalent coordinates (BED file) computed by WGA to align the polymorphic loci referenced by syntenic positions, separating them on each reference genome, and defining them as SHPs. The resulting MSA will have as many subgenomes as reference genomes were used (Figure 1C; 2C). Some of these might be artifacts resulting from residual unspecific reads mapped against one of the reference genomes, which can be detected by the low percentage of reads mapped and therefore valid positions recovered; these must be eliminated by the user. Outgroup species sequences can optionally be added into the SHP alignment by indicating in the configsynt file the BED file of the syntenic positions between the master genome and the outgroup species genome computed with WGA (see details in *Brachypodium* case below).

## Evaluation of the protocol: species, reference genomes and target samples

The protocol was tested and validated using the genome sequences of four diploid-polyploid complexes from three plant groups (*Brachypodium distachyon* complex, *Triticum-Aegilops* complex, and *Brassica* complex) and species from one yeast genus (*Saccharomyces* haploid, diploidized species and synthetic hybrids). The data sets include sequences from the allopolyploids under study and their closest extant diploid progenitor species. Accessions from the diploid species were also included as control samples. The genomes of the diploid species were used as references to conduct the mapping of the reads from the allopolyploid and diploid species of each complex to reconstruct their phylogeny at the subgenome level (Figure S1; Table S1).

The *Brachypodium distachyon* complex analysis included the diploid *B. distachyon* (Vogel et al., 2010) and *B. stacei* (JGI Genome Portal: https://genome.jgi.doe.gov/portal/; Catalan et al. (unpublished)) reference genomes and two diploid *B. distachyon* (ABR2 and Bd21), two diploid *B. stacei* (ABR114 and T.E4.3), and two allotetraploid *B. hybridum* (Bhyb26 and ABR113) ecotypes as samples (JGI Genome Portal; (Gordon et al., 2017, 2020; Scarlett et al., 2023)). *Oryza sativa* Japonica group cv. Nipponbare (NCBI: https://www.ncbi.nlm.nih.gov/; (Shang L et al., 2023)) was included as outgroup species genome (Figure S1A; Table S1).

The *Brassica* complex analysis included the diploid *Brassica oleracea* (Parkin et al., 2014) and diploid *Br. rapa* (Zhang et al., 2018) reference genomes and two diploid *Br. oleracea*, two diploid *Br. rapa* and five allotetraploid *Br. napus* cultivars as samples (Parkin et al. (2014)) (Figure S1B; Table S1).

The *Triticum-Aegilops* complex analysis included the diploid *Triticum urartu* (Ling et al., 2018), diploid *Aegilops tauschii* (L. Wang et al., 2021) and diploid *Ae. speltoides* (L. F. Li et al., 2022) reference genomes and the diploids *T. urartu*, *Ae. tauschii*, *Ae. speltoides*, the allotetraploid *T. turgidum*, and the allohexaploid *T. aestivum* accessions as samples (NCBI; Ling et al., 2018; Zimin et al., 2017) (Figure S1C; Table S1).

*Saccharomyces* yeast analysis included the haploid *Saccharomyces cerevisiae* (Engel et al., 2022), *S. kudriavzevii*, *S. mikatae*, *S. paradoxus (Procházka et al., 2012; Yue et al., 2017)*, and *S. uvarum* reference genomes, and a diploidized *S. cerevisiae* and six other *Saccharomyces* synthetic hybrids [yHRWh24 (*S. cerevisiae x S. kudriavzevii x S. mikatae x S. uvarum*), yHRWh4 (*S. mikatae x S. kudriavzevii*), yHRW134 (Diploidized *S. cerevisiae*), yHRWh13 (S*. mikatae x S. uvarum*), yHRWh10 (*S. cerevisiae x S. uvarum*), yHRWh18 (*S. kudriavzevii x S. paradoxus*), yHRWh51 (*S. uvarum x S. mikatae x S. kudriavzevii*) (see Peris et al., 2020; NCBI)] as samples (Figure S1D; Table S1).

## Evaluation of the protocol: Pre-processing and integrative pipeline application

The same pre-processing of the sequence reads was applied to the four data sets. For each of the samples to be analyzed, the quality check (QC) report was generated using FastQC 0.11.9 software (Andrews, 2019.; http://www.bioinformatics.babraham.ac.uk/projects/fastqc) before and after filtering the sequences using Trimmomatic 0.39 (Bolger et al., 2014). The reads that passed QC were mapped against the concatenated reference genome (including the diploid reference genomes of the progenitor species of the polyploid species under study) using minimap2.1-r1122 (H. Li, 2018, 2021). The mapped reads (SAM format) were converted to BAM format and sorted using samtools 1.16.1 (Danecek et al., 2021) software (--sort and --view functions). Samtools was also used to obtain the statistics of the mapped reads, both in general (--flagstats) and per chromosome of each reference genome (--index and --idxstats). Variant calling was performed using the bcftools 1.16 software (Danecek et al., 2021) (--mpleup with parameters -a DP and -call. The resulting VCF files for each sample were compressed using bgzip 1.19 (Bonfield et al., 2021), and indexed and merged using bcftools (--index and --merge) to create the multi-sample VCF file. This VCF file is used as the input file for the VCF2ALIGNMENT and VCF2SYNTENY algorithms.

Syntenic positions between reference genomes were then calculated using WGA, which requires the *utils/mapcoords.pl* script. By default, this algorithm uses the CGaln software (Nakato & Gotoh, 2010) to compute syntenic blocks and positions. The aligner GSAlign (Lin & Hsu, 2020) was also integrated and tested; however, the analysis performed on the *Brachypodium* reference genomes showed poorer results in terms of recovered syntenic regions and this test was not further extended.

Repetitive regions of the reference genomes had to be masked out, so WGA applied a repetitive region detection and masking procedure by default using Red (Girgis, 2015) and Red2Ensembl.py (Contreras-Moreira et al., 2021). The resulting 0-based BED file contained the list of syntenic positions and was one of the input files used by the VCF2SYNTENY algorithm. In addition, WGA also generated a PDF plot of resulting syntenic blocks by calling the gnuplot 6.0 program (Williams et al., 2023), which helped to visually control the quality of any produced alignments (Figure S2). The parameters used for the four data sets analyzed are listed in Table S2. Bedtools intersect v2.31.1 (Quinlan & Hall, 2010) was used to estimate the proportion of SHPs sitting in annotated genes and non-coding regions of the master genome, as annotated in the respective GFF file.

The VCF2ALIGNMENT algorithm was used to filter the positions in the VCF file that passed the filtering step with respect to the read depth (-d) and number of missing samples (-m) thresholds. In addition, indels were filtered out from the VCF to eliminate downstream inconsistencies. The imposed thresholds were d ≥ 5 (i.e. DP ≥ 5 reads in VCF file) for all data set analyzed, and m ≤ 3 for *Brachypodium*, m ≤ 5 for *Brassica*, m ≤ 3 for *Triticum-Aegilops*, and m ≤ 10 for *Saccharomyces* for missing data. The resulting LOG file with information about the positions that passed the filters was used as the input file for VCF2SYNTENY.

The VCF2SYNTENY algorithm reconciled the information of the syntenic positions obtained by WGA (BED file) and the valid positions obtained by VCF2ALIGNMENT (LOG file), effectively deconvoluting SHPs, which were assigned to the corresponding subgenomes. It is important to note that it is necessary to specify which reference genome will be established as the master genome, since this will be the one used for referencing the positions in the generated multiple sequence alignment (MSA). When using two reference genomes, either one can be selected as the master genome. When three or more genomes are used for evolutionary close species, the longest genome is generally selected as the master genome. However, the results of syntenic blocks obtained with CGaln should always be considered when making this decision.

## Phylogenetic analysis using syntenic homeoSNP datasets

Each of the four SHPs datasets obtained using our protocol was processed by removing artifactual subgenomes with low percentages of mappings and recovered syntenic SNPs. They were then filtered using the snp-sites v.2.5.1 tool (A. J. Page et al., 2016) to retain only the variable positions. Phylogenetics trees were inferred using IQtree v.2.2.2.6 software (Minh et al., 2020) with parameters -m MFP+ASC -AICc -alrt 1000 -B 1000 -T AUTO. Phylograms were rooted at the midpoint, except for the *Brachypodium* phylogram, which was rooted using the *O. sativa* outgroup, and visualized using the FigTree v.1.4.4 software (Rambaut, 2018; http://tree.bio.ed.ac.uk/software/figtree/). The same alignment used to infer the phylogeny was used to calculate the genome-wide average nucleotide identity (gwANI) using the pANIto software (A. J. Page et al., 2018; https://github.com/sanger-pathogens/panito).

## Results

### Proportion of reads mapped against diploid reference genomes

The proportions of reads mapped using minimap2 against each concatenated diploid progenitor reference genome were as expected. Thus, the reads of the diploid species were mostly mapped against their corresponding diploid species reference genome. The ecotypes of the diploid species *B. stacei* and *B. distachyon* mapped between 92% and 98% against their respective genomes (Table S3A). The cultivars of the diploid species *Br. oleracea* and *Br. rapa* mapped between 78% and 88% of reads (Table S3B). Between 87% and 97% of reads from diploids of the *Triticum-Aegilops* complex, *Ae. speltoides*, *Ae. tauschii* and *T. urartu*, mapped to their respective genomes (Table S3C). In yeast, diploidized *S. cerevisiae* mapped 96% of reads to its reference genome (Table S3D).

Regarding the polyploid samples analyzed, the number of mappings against each reference genome varied depending on the species and ploidy. Thus, the mapping ratio of the allotetraploid ecotypes of *B. hybridum* (DDSS) was approximately 50:50 (subgenome D: subgenome S), but with some differences between ecotypes. The ABR113 ecotype showed 50% of the reads mapped against the *B. distachyon* reference genome compared to 54% of the Bhyb26 ecotype (subgenome D) and the respective 49% and 46% against the *B. stacei* reference genome (subgenome S) (Table S3A). The accessions of the allotetraploid *Brassica napus* (ArArCoCo) also showed a 50:50 ratio of mappings to their reference genomes, with slightly more mappings against the *Br. rapa* (subgenome Ar) reference genome (51-52%) than against the *Br. oleracea* reference genome (subgenome Co) (48-49% of mappings) (Table S3B). In the case of the *Triticum-Aegilops* complex, the allotetraploid *T. turgidum* (AABB) mapped predominantly against its progenitors, *T. urartu* (46%; subgenome A) and *Aegilops speltoides* (40%; subgenome B), while 14% of the reads that mapped against *Ae. tauschii* were considered artefacts and eliminated from downstream analyses. The allohexaploid *T. aestivum* (AABBDD) mapped 34% against *T. urartu* (subgenome A), 29% against *Ae. speltoides* (subgenome B) and 37% against *Ae. tauschii* (subgenome D) (Table S3C).

The six *Saccharomyces* synthetic hybrids analyzed mapped predominantly against their parents, but with high variability in their proportions (Table S3D). The synthetic hybrid Sce x Sku x Smi x Suv mapped predominantly against two of its parents, *S. mikatae* (Smi; 40%) and *S. kudriavzevii* (Sku; 38%), and to a lesser extent against its two other parents, *S. cerevisiae* (Sce; 12%) and *S. uvarum* (Suv; 11%). A very small percentage of reads (Spa; 0.1%) mapped against the reference genome *S. paradoxus*, which is not a parent of this hybrid. The synthetic hybrid Smi x Sku mapped 51% and 48% of reads against its two reference genomes, *S. mikatae* (Smi) and *S. kudriavzevii* (Sku), respectively. Less than 1% of the reads mapped against the other three non-parental reference genomes of the synthetic hybrid. The Smi x Suv hybrid mapped 60% and 68% against its parental *S. mikatae* (Smi) and *S. uvarum* (Suv) reference genomes, respectively. Less than 2% of the remaining reads mapped to the genomes of non-progenitor species. The Sce x Suv hybrid mapped 59% of reads against its progenitor genome *S. cerevisiae* (Sce) and 38% against that of *S. uvarum* (Suv). The remaining mappings (3%) were likely unspecific. The synthetic hybrid Sku x Spa, resulting from the cross between *S. kudriavzevii* x *S. paradoxus*, showed a balanced proportion of 48% mapping to both parental genomes. Finally, the Suv x Smi x Sku synthetic hybrid from the cross between *S. uvarum*, *S. mikatae* and *S. kudriavzevii* mapped predominantly against its parental genomes *S. mikatae* (Smi; 38%) and *S. kudriavzevii* (Sku; 37%), and less to that of *S. uvarum* (Suv; 24%) (Table S3D).

### Syntenic positions recovered among diploid progenitor reference genomes by WGA

The computation of syntenic positions between diploid reference genomes, master versus secondary, using WGA and its main dependency CGaln, is a fundamental step for the detection of SHPs from subgenomes of the polyploids under study. The number of syntenic positions recovered in each data set varied depending on the homology between the different chromosomes of the species. Thus, the reference genomes used in the *Brachypodium* group presented the highest percentage of synteny among tested plants. The *Brachypodium stacei* reference genome (ABR114 ecotype) had syntenic positions covering 29% of the *B. distachyon* reference genome (Bd21 ecotype) determined as a master reference genome (Figure S2A; Table S4A). As expected, the syntenic regions between *B. distachyon* (master genome) and *Oryza sativa* (outgroup genome) were notably reduced to 2.5% of the *B. distachyon* genome. Most of these regions (98.6%) were located within coding regions (Table S4A). In the case of *Brassica*, this percentage was significantly reduced, with syntenic positions between the *Br. rapa* and *Br. oleracea* reference genomes covering only the 5% of the *Br. oleracea* master reference genome (Figure S2B; Table S4B). Similarly, synteny between the reference genomes of *Aegilops* and *T. urartu* was approximately 3% of the *T. urartu* master reference genome (Figure S2C; Table S4C). In the yeast example, the compared species showed synteny ranging from 32% (*S. uvarum*) to 55% (*S. paradoxus*) of the master reference genome *S. cerevisiae* (Figure S2D; Table S4D).

The percentage of syntenic positions detected in coding regions (genes) of the master genome varied significantly among the species analyzed. In *Brachypodium* (Table S4A), 73.5% of detected syntenic positions were located in genes, compared to only 48-43% in *Brassica* (Table S4B) and *Triticum*-*Aegilops* (Table S4C). In yeast, 86.8-96.2% of the syntenic positions were located in coding regions (Table S4D).

### Single Homeologous Polymorphisms (SHPs) recovered by AlloSHP pipeline

The percentages of SHPs recovered for each subgenome (Table S5A-D) generally correlate with those obtained in the mappings (Table S3A-D). However, in some polyploids, changes in the number of predominant SHPs were observed. For example, the same proportion of SHPs was recovered in the ecotypes ABR113 and Bhyb26 the allotetraploid *B. hybridum*, with slightly more SHPs obtained from the S-subgenome (Bsta; 50.5%) than from the D-subgenome (Bdis; 49.5%) (Table S5A), while these proportions were inverse in the mappings (Table S3A).

The accessions of the allotetraploid *Brassica napus* showed the same proportion of SHPs (Table S5B) as the mappings (Table S3B), with percentages of 51-52% of SNPs from the A subgenome (Brr*; Br. rapa*) and 48-49% of SHPs from the C subgenome (Bro; *Br. oleracea*) (Table S5B).

However, in the *Triticum-Aegilops* group, some variations between mapping (Table S3C) and SHPs recovered for each subgenome were detected (Table S5C). As expected, in the *Ae. tauschii* and *T. urartu* diploid samples, 96% and 99% of the recovered SHPs (Table S5C) came from mappings against their own reference genomes. However, in the case of the diploid *Ae. speltoides* sample analyzed, although 86% of the reads mapped against its own reference genome (Table S3C), the percentage of recovered SHPs was more unspecific, showing 65%, 22% and 13% SHPs recovered from the *Ae. speltoides*, *Ae. tauschii* and *T. urartu* reference genomes, respectively (Table S5C). In the allotetraploid *T. turgidum*, 53% and 34% of the recovered SHPs correspond to subgenome A (Tur; *T. urartu*) and subgenome B (Aes; *Ae. speltoides*), respectively (Table S5C). SHPs recovered from the allohexaploid *T. aestivum* are distributed among its three subgenomes A, B and D with percentages of about 38%, 20% and 42%, respectively (Table S5C). The greatest variation in the percentage of mappings (Table S3A-C) and SHPs detected for each subgenome (Table S5A-C) may be influenced by the percentage of masked genome sequences, i.e., the number and size of repetitive regions. In the three reference genomes of *Triticum*-*Aegilops* complex species, 80–86% of the genomes were masked. In contrast, in the reference genomes of *Brachypodium* and *Brassica*, the percentage was 31–35%.

The proportions of SHPs recovered in the synthetic yeast hybrids (Table S5D) differed from previously obtained mappings (Table S3D), especially in those recovered from the parent *S. uvarum*, with notable reductions. This is because the reference genome of *S. uvarum* showed only 32.2% synteny with the primary reference genome *S. cerevisiae* (Table S4D). Meanwhile, the genomes of *S. kudriavzevii*, *S. mikatae*, *and S. paradoxus* showed proportions of 42%, 44%, and 55%, respectively (Table S4D). Furthermore, the *S. uvarum* genome has a higher percentage of masked regions due to repetitive sequences (38% of masked genome) than the other *Saccharomyces* reference genomes (25-27% of masked genome). The hybrid *S. mikatae* x *S. kudriavzevii* showed a very similar proportion of SHPs (Smi: 52%; Sku: 48%; Table S5D) to mapped reads (Smi: 51%; Sku: 48%; Table S3D). However, the hybrid *S. mikatae* x *S. uvarum* and *S. cerevisiae* x *S. uvarum* showed a marked predominance of SHPs from the parent *S. mikatae* genome (78%), in the Smi x Suv hybrid, and from the *S. cerevisiae* genome (81%) in the Sce x Suv hybrid, compared to its other parent *S. uvarum* with percentages of SHPs of 22% and 19%, respectively (Table S5D). It should also be noted that the proportions of each parental genome in the synthetic hybrids are variable (see Peris et al., 2020). The synthetic hybrid resulting from the cross of three parents, *S. uvarum* x *S. mikata*e x *S. kudriavzevii* also showed a notable reduction in the number of SHPs of the parent *S. uvarum* (SHPs: 11%, Table S5D) versus 24% mapped reads (Table S3D) compared to the predominance of its other two parents *S. kudriavzevii* (SHPs: 43%; mappings: 37%) and *S. mikatae* (SHPs: 46%; mappings: 38%). The synthetic hybrid resulting from the crossing of four parents, *S. cerevisiae* x *S. kudriavzevii* x *S. mikatae* x *S. uvarum*, also showed this trend with reduced values compared to the *S. uvarum* parent (SHPs: 3%; mappings: 11%), while the *S. cerevisiae* parent also showed reduced (11%) but similar percentages of SHPs to the mappings (12%). The genomes of the parents *S. mikatae* (SHPs: 45%; mappings: 40%) and *S. kudriavzevii* (SHPs: 41%; mappings: 38%) were predominant in this synthetic hybrid (Tables S5D; S3D).

### Inference of subgenomic phylogenies from SHPs

The multiple alignments of SHPs used for phylogenetic analysis contain 5,958,612, 980,493, 8,556,827, and 1,540,088 total sites for the *Brachypodium*, *Brassica*, *Triticum-Aegilops*, and *Saccharomyces* groups, of which 4,378,851, 787,595, 6,377,583, and 1,441,544 are parsimony-informative, respectively (Table S6).

The inferred phylogeny in the *Brachypodium distachyon* complex shows 100/100 SH-aLRT/UltraFast bootstrap support across all branches (Figure 3A). A total of 642,818 SNPs from *Oryza sativa*, syntenic to the *B. distachyon* master genome, were included in the MSA to root the *Brachypodium* tree. Both *B. hybridum*-Bhyb26 subgenomes (S and D) are resolved as more ancestral than those of the recent *B. hybridum*-ABR113 subgenomes. Furthermore, these two subgenomic lineages, Bhyb26-S and Bhyb26-D, diverged earlier than their respective progenitor species lineages (*B. stacei* TE4.3 and ABR114; *B. distachyon* Bd21 and ABR2), suggesting that the ancestral B*. stacei* and *B. distachyon* parents of *B. hybridum* Bhyb26 went extinct or have not been sampled yet (Gordon et al., 2020; Scarlett et al., 2023). This demonstrates the potential of our method applied to intra/inter-species studies to infer the different putative diploid progenitors (subgenomes), distinguishing within the same species the most ancestral and recent ones. (Figure 3A). The gwANI matrix (Table S7A) showed higher identities between *B. hybridum* ABR113 ecotype subgenomes D and S and the genomes of ecotypes of the diploid progenitor species *B. distachyon* and *B. stacei* (D-subgenome vs *B. distachyon* ecotypes: 94–95%; S-subgenome vs *B. stacei* ecotypes: 96–97%) than those obtained for the *B. hybridum* Bhyb26 ecotype subgenomes (D-subgenome vs *B. distachyon* ecotypes: 87–88%; S-subgenome vs *B. stacei* ecotypes: 85%) (Table S7A). This was reflected in the higher divergences of the Bhyb26 subgenomes from its diploid progenitor species compared to that of ABR113 as shown in the phylogenetic tree (Figure 3A) and in agreement with previous finding (Gordon et al., 2020; Mu et al., 2023; Scarlett et al., 2023).

**Figure 3.**
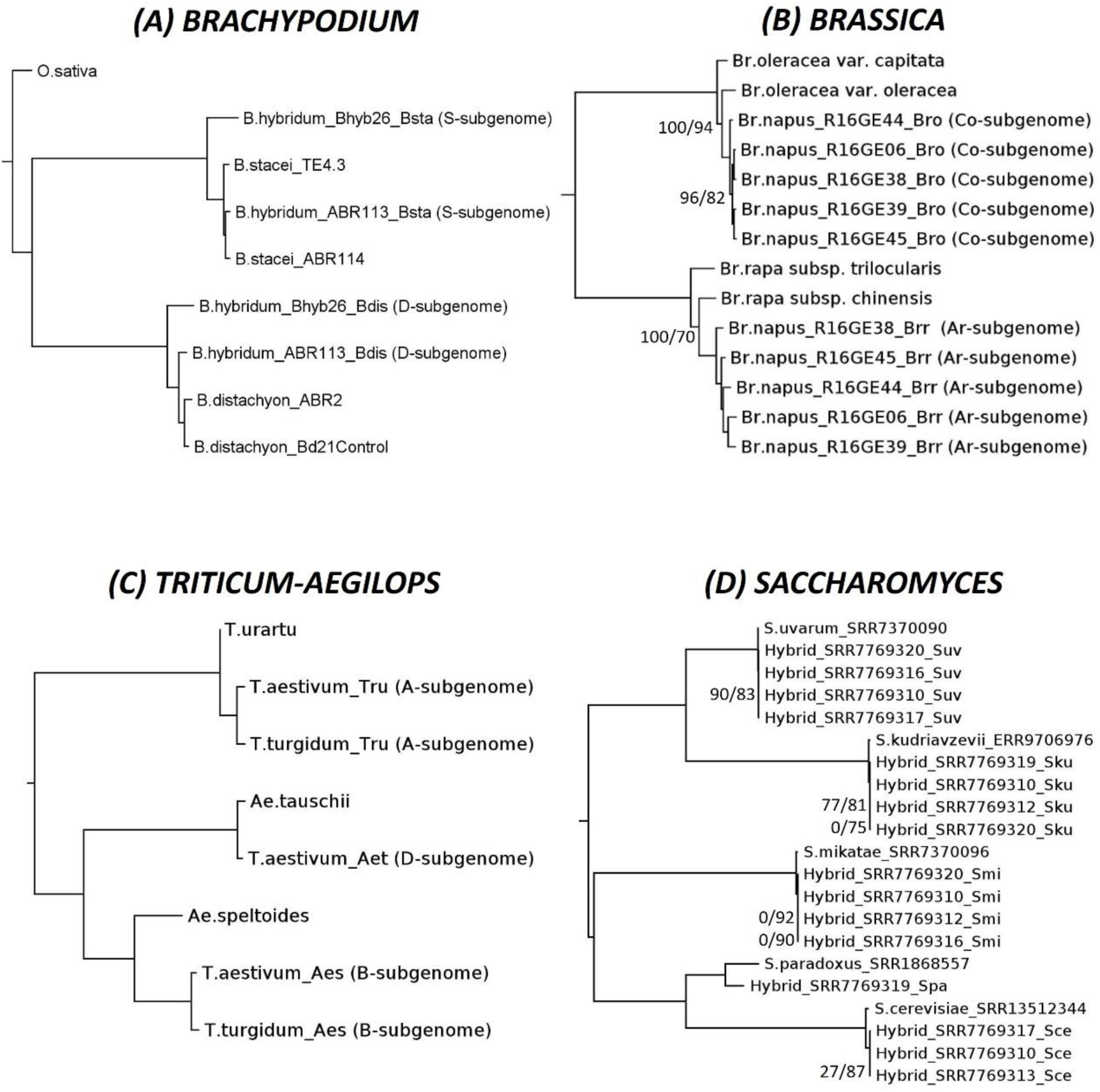
Phylograms inferred using the SHPs alignment from the four diploid-polyploid complex datasets [**(A)** *Brachypodium distachyon* complex, **(B)** *Brassica* complex, **(C)** *Triticum-Aegilops* complex, and **(D)** *Saccharomyces* haploids and synthetic hybrids] used for the pipeline validation. Numbers indicate branches with SH-aLRT/UltraFast Bootstrap supports (BS) <80/95; the remaining branches have 100/100 values.

The phylogeny of the *Brassica* complex showed high support across all branches. Only three branches had high support but below the recommended 80 and 95 SH-aLRT and UFboot thresholds (Figure 3B). Two groups were distinguished based on the two diploid species, *Br. oleracea* and *Br. rapa*, progenitors of the allotetraploid *Br. napus*, and the respective subgenomes of the allotetraploid accessions clustered according to their diploid progenitor. Among *Br. rapa* subspecies, subsp. *tricularis* was more ancestral than subsp. *chinensis*. Regarding the Co-subgenome (*Br. oleracea* subgenome), the *Br. napus* R16G44 sample showed a more ancestral divergence than the other samples, which were resolved into two (R16GE06/ R16GE38 and R16GE39/ R16GE45) more recently evolved sister groups. In contrast, the Ar-subgenome clade (*Br. rapa*), showed the successive divergences of the R16G38, R16GE06 and R16GE39 lineages. As in the *Brachypodium distachyon* complex, different phylogenetic resolutions for one and the other subgenomic lineages were also detected in the *Brassica* allotetraploid species studied (Figure 3B). The gwANI values of the Co (Bro) subgenomes of *Br. napus* showed a higher similarity among them (96.6-98%) than among the Ar (Brr) subgenomes (90.5-94.8%) (Table S7B). These results, at the intraspecific level in *Br. napus* and focusing only in the syntenic regions shared between progenitors, agree with those obtained by Khan et al. (2016) using HANDS2 software to compare the subgenomes of *Brassica carinata* (BBCC), *Br. juncea* (AABB), and *Br. napus* (AACC), indicating that the three C genomes of *Brassica* are more similar to each other than the three A genomes.

The inferred phylogeny in the *Triticum-Aegilops* complex showed 100/100 SH-aLRT/UltraFast Bootstrap support across all branches (Figure 3C). The two main clades corresponded to *T. urartu* and the A-subgenomes of *T. aestivum* and T. *turgidum*, and the *Aegilops* clade, including *Ae. tauschii* and the D-subgenome of *T. aestivum*, and *Ae. speltoides* with the B-subgenomes of *T. turgidum* and *T. aestivum* (Figure 3C) matching previous studies (Glémin et al., 2019; L. F. Li et al., 2022; Marcussen et al., 2014). The A subgenomes (Tru) of the allopolyploids *T. turgidum* and *T. aestivum* showed a high gwANI of 96.1%, as well as the B subgenomes (Aes) of the same species with 95.2% (Table S7C). The identity between the subgenomes of allopolyploids and their closest diploid ancestors varied considerably. The D subgenome of T. aestivum was 96.6% identical to *Ae. tauschii*. However, this decreased to 91% between the A subgenomes and the *T. urartu* progenitor, and to 68-70% between the B subgenomes and the *Ae. speltoides* progenitor (Table S7C).

The phylogeny of the yeast *Saccharomyces* showed two clades. One formed by the *species S. uvarum* and *S. kudriavzevii* and the subgenomes of hybrids shared with one of these parents, and the other clade formed by the species *S. mikatae* and the sister species *S. paradoxus* and *S. cerevisiae* (Figure 3D). The subgenomes of the hybrids were grouped with the corresponding parental genomes. The support for the nodes of the main clades and groups was 100/100. This support dropped significantly within the subgenomic clades from the same parent. These clades often collapsed into polytomies due to the similarity of the recovered syntenic SNP sets, but were divergent from their parental genome (Figure 3D). The gwANI for subgenomes within a single parent among the different synthetic hybrids studied was 100% (Table S7D). Given the origin of these hybrids (Peris et al., 2020), their high level of identity is not surprising. However, the use of this set of synthetic samples resulting from multiparent crosses was used in the present study as a proof of concept to analyze the behavior of our protocol with such many reference genomes and the corresponding synthetic hybrids.

## Discussion

### Strengths and limitations of the protocol

This protocol does not require the targeted polyploids to be assembled nor annotated, as the sequenced reads are mapped directly against the reference genomes of the diploid progenitor species. This allows the selection of SHPs from both coding and intergenic regions (excluding repetitive regions). This approach allows us to reconcile a large number of polyploid subgenomes in a single alignment (Figures 1 and 2). In the analyses performed on allotetraploids (*Brachypodium hybridum*, *Triticum turgidum*, *Brassica napus*), allohexaploid (*Triticum aestivum*), and synthetic yeast hybrids resulting from crosses of up to four different parents (synthetic *Saccharomyces* hybrids) (Figure S1; Table S1), we did not observe any limitation with respect to the ploidy of the allopolyploid under study, as long as the genomes of its extant diploid progenitor species are available and show syntenic regions between them.

Due to the mapping and syntenic alignment strategy, our protocol is only effective for inferring the subgenomes of allopolyploid organisms, not autopolyploids. Therefore, it is necessary to know which diploid species are the progenitor species or the closest extant relatives of the polyploid species under study, and the reference genomes of these diploids must be available. However, it is also possible to analyze polyploids whose progenitor species are uncertain. To do this, all diploid species that could be potential progenitors should be included as putative progenitors, and the resulting percentage of mappings could provide clues to their most likely involvements as progenitor species. Caution should be taken, however, for those allopolyploid species with unknown, extinct, or closely related putative progenitor species.

Another limitation is related to the variants considered downstream, since to avoid conflicts between the positions extracted from the VCF file and the syntenic positions of the reference diploid progenitor genomes, indels are discarded and only SNPs are used for downstream analyses.

Some sources of bias may arise from the sequencing quality of the polyploid reads, as well as the assembly quality of the diploid reference genomes used (Phillips, 2024). To this end, the user must set thresholds and filters for FastQC, mapping quality, syntenic blocks, additional VCF filters resulting from the mappings and variant calling tools used.

Although we have not found any limitation of our pipeline with respect to the size of the samples to be analyzed, it must be considered that some of the algorithms can generate bulky output files (tens of GB in the case of the *Triticum-Aegilops* complex) and might have high RAM requirements, with peaks that can exceed 100 GB in the case *of Triticum-Aegilops*. The size of the output files, the required RAM and the processing time are directly related to the size of the VCF input file and the size and number of reference genomes used (Table S8A, B).

### Technical considerations of syntenic alignment, mapping and variant calling steps

Some of the technical and methodological considerations that users should consider are related to obtaining the VCF file, a step prior to using our pipeline. Although this study does not propose optimal filtering parameters and thresholds for the mapping and variant calling process in the polyploid genome analysis, there are previous studies and reviews such as those conducted by Clevenger et al. (2015), Cooke et al. (2022) and Phillips (2024), that suggest some values for these parameters as applied to polyploidy species studies. In any case, the parameters should be fine-tuned based on the polyploid organism and sequencing data (Ning et al., 2024). Likewise, the parameters applied in the CGaln software used to obtain the syntenic regions must also be optimized in each case, although in this work those indicated gave the best results for the allopolyploid organisms studied, which suppose a good representation due to their wide range of genome sizes and ploidies.

### Threshold of artifactual subgenomes

Since our pipeline infers as many subgenomes as reference genomes are used, another methodological aspect to consider is setting the minimum threshold of SHPs required to establish a subgenome as plausible and not an artifact produced by non-specific mapping of reads against some of the reference genomes. These cases should be reduced by increasing parameters such as mapping quality (mapQ) or read depth (DP), while maintaining a balance so as not to miss an excessive number of SHPs. In our study, this threshold varies between the organisms studied. In *Brachypodium*, plausible subgenomes were those with a percentage of SNPs (in diploid species) or SHPs (in allopolyploids) greater than 1.2% (Table S5A). In *Brassica*, this percentage increased to 14.8%, as *Brassica oleracea* var. *capitata* had a non-specific mapping to the reference genome of 16.3% of the reads, representing 14.8% of artifactual SHPs (Table S5B). In *Triticum*, the threshold was also raised to 21.6%, as this percentage of artifactual SHPs was recovered from the *Ae. speltoides* sample of reads non-specifically mapped to the *Ae. tauschii* genome (Table S5C). In the case of *Saccharomyces*, the percentage of artifactual SHPs was less than 0.4% of the total number of SHPs detected (Table S5D). These cutoffs are influenced by the quality of the assembly, the evolutionary proximity and the genomic identity between the progenitor reference genomes used. In any case, if the diploid progenitor species of the allopolyploids under study are known previously, this decision should not be problematic.

Two tests were performed on the *Brachypodium* data set to analyze the specificity of the mappings against the concatenated reference genomes. The first test consisted of using three reference genomes, including the two progenitor diploid species (*B. distachyon* and *B. stacei*; see Table S9) and a third non-progenitor diploid species (*B. sylvaticum*; *Brachypodium sylvaticum* v1.1 DOE-JGI) of the allotetraploids *B. hybridum*. The results indicated that the two *B. hybridum* ecotypes (Bhyb26 and ABR113) showed reduced non-specific mappings against the non-progenitor reference genome *B. sylvaticum*, accounting for only 4.4% and 2.3% of the total reads, respectively (Table S9).

The second test was designed to verify how the specificity of the mappings varies with the evolutionary proximity of the samples used as reference genomes. To do this, we mapped the reads of three *Brachypodium distachyon* ecotypes corresponding to each of the three clades/phylogenetic groups of this species (ABR2 from the Spanish group [S+ from the S+T+ clade], Bd21 from the Turkish group [T+ from the S+T+ clade], and BdTR8i from the EDF (Extremely Delayed Flowering) clade) against concatenated *B. distachyon* ecotypes genomes from the *B. distachyon* pangenome (Gordon et al., 2017). The EDF and S+T+ clades diverged less than one million years ago, and the S+ group is paraphyletic and, together with the monophyletic T+ group, forms the Spanish-Turkish clade, which are evolutionarily close (Gordon et al., 2017, 2020; Sancho et al., 2018). When we mapped the reads against two reference genomes (Table S10A), Tek2 from the EDF clade and ABR3 from the S+ group (S+T+ clade), 81% of the reads from the ABR2 sample (S+ group) mapped against the ABR3 genome (S+ group). The Bd21 reads (T+ group) mapped mostly (64%) against the closest genome, in this case the ABR3 genome (S+ group). The BdTR8i sample from the EDF clade mapped mostly (62%) against the Tek2 reference genome of this clade (Table S10A). When the same accessions, ABR2, Bd21 and BdTR8i, were mapped against the concatenated genomes of three *B. distachyon* ecotypes, one from each clade/group (Tek2 [EDF], ABR3 [S+] and BdTR12c [T+]) instead of only two, the percentages of mapped reads were more specific (Table S10B). ABR2 reads (S+) mapped 13%, 58% and 29% against the EDF, S+ and T+ references, respectively. BdTR8i (EDF) mapped 55%, 27.5% and 17.5%, respectively (Table S9B). Unlike the previous test, where the majority of the reads mapped to the reference genome of its own clade/group (Table S10A), in sample Bd21 (T+) 41% of reads mapped against the genome of the S+ group, followed by 35% and 24% of the T+ group and the EDF clade, respectively (Table S10B). In this case, we used an extreme example, where the reference genomes were ecotypes of the same species, and therefore they were extremely close, showing similar genomes (Gordon et al., 2017, 2020). Therefore, for adequate application of our pipeline we need to consider the evolutionary proximity of the reference genomes, especially at the intraspecific level. Another aspect to consider is the quality of the assembly of the reference genomes. In this test, preliminary versions of the genomes assembled from the *B. distachyon* pangenome were used (Gordon et al., 2017).

### Accuracy and perspectives of the subgenomic phylogenetic reconstruction of *Brachypodium*, *Triticum-Aegilops*, and *Brassica* groups, and *Saccharomyces* haploid-synthetic hybrids

The phylogeny of *the B. distachyon* complex inferred from SHPs (Figure 3A) showed the highest support for all the nodes and was consistent with that of Gordon et al. (2020). Both *B. stacei* and *B. distachyon* clustered with the respective S and D subgenomes of the allotetraploid *B. hybridum*, and the most divergent positions of both subgenomes of the older *B. hybridum* Bhyb26 ecotype were recovered with respect to their parents and the subgenomes of the more recent *B. hybridum* ABR113 ecotype. The *Brassica* phylogenetic tree showed divergences among and within the Ar and Co subgenomes of the five *Br. napus* accessions analysed (Figure 3B). Multiple phylogenetic and population studies have been carried out on the diploid species *Br. oleracea* and *Br. rapa*, as well as on the allotetraploid *Br. napus* (An et al., 2019; Bird et al., 2017; Gazave et al., 2016; Lv et al., 2020; Song et al., 2020). However, these studies require further expansion to confirm the multi-origin hypothesis of *Br. napus*. The present study used a small database to validate the pipeline, and using a large number of accessions is beyond this objective. Without further information on these samples and a much more extensive sample of *Br. rapa*, *Br. oleracea* and *Br. napus* accessions, no further conjectures on the inferred phylogeny can be made. Khan et al. (2016).

The genomes and phylogenies of the allopolyploids *T. aestivum* (6x) and *T. urartu* (4x) have been extensively studied, with *Ae. speltoides*, *Ae. tauschii* and *T. urartu* being proposed, albeit with some controversy, as their closest extant diploid progenitors (Appels et al., 2018; El Baidouri et al., 2017; Glémin et al., 2019; L. F. Li et al., 2022; Ling et al., 2018; Marcussen et al., 2014; Petersen et al., 2006). The phylogeny of the *Triticum-Aegilops* complex recovered in the present study showed the expected groupings among the allopolyploid *Triticum* subgenomes (A, B and D) and their closest extant diploid ancestral relative (Marcussen et al., 2014; Petersen et al., 2006). Comparing the number of SHPs detected in the *T. aestivum* and *T. turgidum* subgenomes, the number of SHPs in the A subgenomes of both species was similar (Table S5C). The *T. aestivum* D subgenome also recovered a high number of SHPs, even higher than the A subgenome. However, the number of SHPs in the B subgenome, although numerous, was reduced in both allopolyploid species by an order of magnitude compared to those recovered in the other subgenomes. Regarding base assignment to the B subgenome, Mithani et al. (2013) detected that their HANDS protocol reduced the accuracy of base assignment for the B subgenome and attributed this to the fact that *Ae. speltoides*, used as the diploid progenitor species of the B genome, is evolutionarily closer to the *T. aestivum* B subgenome than the progenitor species of the A and D subgenomes. The *Saccharomyces* phylogeny shows polytomies in all subgenomes (Figure 3D), with 100% gwANI for subgenomes within a single parent among the different synthetic hybrids studied (Table S10D). Given the origin of these hybrids (Peris et al., 2020), their high level of identity is not surprising. However, the use of this set of synthetic samples resulting from multiparent crosses was used in the present study as a proof of concept to analyze the precision of our protocol with such many reference genomes and the corresponding synthetic hybrids.

The proportions of SHPs in the synthetic hybrids (Table S5D) from the different parents were generally consistent with the genomic contributions shown in Figures 2b and 4 in Peris et al. (2020), with the notable exception of SHPs from the *S. uvarum* progenitor. These were reduced compared to the actual proportions in each parental genome. This bias is due to the lower synteny between the *S. uvarum* genome, and the master or primary *S. cerevisiae* genome established in our protocol (Table S4D).

The phylogenies obtained with AlloSHP in the four sets of diploid-allopolyploid species analyzed are consistent with previous studies that used different methodologies and nucleotide sequences (e.g. DNA and RNA loci). Therefore, AlloSHP can facilitate scaling up this type of analysis by increasing the number of allopolyploid and diploid progenitor populations and accessions, with the aim of studying their phylogenetic relationships at the subgenome level in greater detail.

## Conclusions

A simple command-line pipeline has been developed to detect and extract SHPs from the homeologous subgenomes of allopolyploid species by mapping the reads against the concatenated reference genomes of their extant progenitor diploid species and reconciling SHPs into a subgenomic multiple sequence alignment using the syntenic positions of the reference genomes. This protocol allows to generate from a single VCF file the SHP alignment necessary to perform subgenome-scale phylogenetic studies of allopolyploid organisms, requiring only the genomes of their closest existing diploid progenitors and the genomic or transcriptomic sequences of the allopolyploids under study. This novel approach provides a valuable tool for the evolutionary study of allopolyploid species, both at the inter- and intra-specific levels, allowing the simultaneous analysis of a large number of accessions and avoiding the complex process of assembling polyploid genomes.

## Availability and requirements

*Project name: AlloSHP*

*Project home page:* https://github.com/eead-csic-compbio/AlloSHP

*Operating system(s): Linux and MacOS*

*Programming language: Perl (95.3%), R (1.3%) and Makefile (3.4%)*

*Other requirements: Standard Linux utilities (gzip, grep, sort, perl, make, python3, g++, libdb-dev), Perl libraries (Getopt::Std, File::Temp, File::Basename, FindBin, DB_File, FileHandle) and third-party dependencies (Cgaln, GSAlign, Red, Red2Ensembl.py and gnuplot)*.

*License:* Apache License 2.0

## List of abbreviations

Aes: *Aegilops speltoides*
Aet: *Aegilops tauschii*
BAM: Binary Alignment Map format
BED: Browser Extensible Data format
Bdis: *Brachypodium distachyon*
Bsta: *Brachypodium stacei*
Bro: *Brassica oleracea*
Brr: *Brassica rapa*
DP: Total read depth
EDF: Extremely Delayed Flowering clade
gwANI: genome-wide Average Nucleotide Identity
homeoSNPs: homeologous Single Nucleotide Polymorphisms
HSPs: Homeolog-Specific Polymorphisms
MSA: Multiple Sequence Alignment
Osat: Oryza sativa
QC: Quality Check
S+: Spanish group
SAM: Sequence Alignment Map format
Sce: *Saccharomyces cerevisiae*
SHPs: Single Homeologous Polymorphisms
Sku: *Saccharomyces kudriavzevii*
Smi: *Saccharomyces mikatae*
Spa: *Saccharomyces paradoxus*
Suv: *Saccharomyces uvarum*
T+: Turkish group
Tru: *Triticum urartu*
VCF: Variant Call Format
WGD: Whole Genome Duplication

## Declarations

### Availability of data and materials

The datasets generated and/or analysed during the current study are available in public databases (Phytozome13, Genome Portal and NCBI), Supplementary materials and AlloSHP repository, https://github.com/eead-csic-compbio/AlloSHP.

### Authors’ contributions

R.S., B.C.-M., P.C planned and designed the study and supervised the work. B.C.-M., R.S. wrote the source code and documentation. J.V. provided data. R.S. drafted the manuscript. All authors contributed to the discussion of the results, the writing and approval of the final manuscript.

## Acknowledgements

The authors thank Miguel Campos (Escuela Politécnica Superior de Huesca, Universidad de Zaragoza) and Francesc Montardit-Tarda (Estación Experimental de Aula Dei, Consejo Superior de Investigaciones Científicas) for their valuable comments to improve the code and implementation of the AlloSHP scripts.

## References

Albertin, W., & Marullo, P. (2012). Polyploidy in fungi: Evolution after whole-genome duplication. In Proceedings of the Royal Society B: Biological Sciences (Vol. 279, Issue 1738, pp. 2497–2509). Royal Society of London. 10.1098/rspb.2012.0434

An, H., Qi, X., Gaynor, M. L., Hao, Y., Gebken, S. C., Mabry, M. E., McAlvay, A. C., Teakle, G. R., Conant, G. C., Barker, M. S., Fu, T., Yi, B., & Pires, J. C. (2019). Transcriptome and organellar sequencing highlights the complex origin and diversification of allotetraploid Brassica napus. Nature Communications, 10(1). 10.1038/s41467-019-10757-1

Andrews, S. (2019). FastQC. A quality control tool for high throughput sequence data. (0.11.9).

Appels, R., Eversole, K., Feuillet, C., Keller, B., Rogers, J., Stein, N., Pozniak, C. J., Choulet, F., Distelfeld, A., Poland, J., Ronen, G., Barad, O., Baruch, K., Keeble-Gagnère, G., Mascher, M., Ben-Zvi, G., Josselin, A. A., Himmelbach, A., Balfourier, F., … Wang, L. (2018). Shifting the limits in wheat research and breeding using a fully annotated reference genome. Science, 361(eaar7191). 10.1126/science.aar7191

Bayer, P. E., Golicz, A. A., Scheben, A., Batley, J., & Edwards, D. (2020). Plant pan-genomes are the new reference. Nature Plants, 6(8), 914–920. 10.1038/s41477-020-0733-0

Bird, K. A., An, H., Gazave, E., Gore, M. A., Pires, J. C., Robertson, L. D., & Labate, J. A. (2017). Population structure and phylogenetic relationships in a diverse panel of Brassica rapa L. Frontiers in Plant Science, 8(321). 10.3389/fpls.2017.00321

Bolger, A. M., Lohse, M., & Usadel, B. (2014). Trimmomatic: A flexible trimmer for Illumina sequence data. Bioinformatics, 30(15), 2114–2120. 10.1093/bioinformatics/btu170

Bombarely, A., Coate, J. E., & Doyle, J. J. (2014). Mining transcriptomic data to study the origins and evolution of a plant allopolyploid complex. PeerJ, 2(e391). 10.7717/peerj.391

Bonfield, J. K., Marshall, J., Danecek, P., Li, H., Ohan, V., Whitwham, A., Keane, T., & Davies Robert M. (2021). HTSlib: C library for reading/writing high-Throughput sequencing data. GigaScience, 10(2), 1–6. 10.1093/gigascience/giab007

Clevenger, J., Chavarro, C., Pearl, S. A., Ozias-Akins, P., & Jackson, S. A. (2015). Single nucleotide polymorphism identification in polyploids: A review, example, and recommendations. Molecular Plant, 8(6), 831–846. 10.1016/j.molp.2015.02.002

Contreras-Moreira, B., Filippi, C. V., Naamati, G., García Girón, C., Allen, J. E., & Flicek, P. (2021). K-mer counting and curated libraries drive efficient annotation of repeats in plant genomes. Plant Genome, 14(e20143). 10.1002/tpg2.20143

Cooke, D. P., Wedge, D. C., & Lunter, G. (2022). Benchmarking small-variant genotyping in polyploids. Genome Research, 32(2), 403–408. 10.1101/GR.275579.121

Danecek, P., Bonfield, J. K., Liddle, J., Marshall, J., Ohan, V., Pollard, M. O., Whitwham, A., Keane, T., McCarthy, S. A., Davies, R. M., & Li, H. (2021). Twelve years of SAMtools and BCFtools. GigaScience, 10(2). 10.1093/gigascience/giab008

El Baidouri, M., Murat, F., Veyssiere, M., Molinier, M., Flores, R., Burlot, L., Alaux, M., Quesneville, H., Pont, C., & Salse, J. (2017). Reconciling the evolutionary origin of bread wheat (Triticum aestivum). New Phytologist, 213(3), 1477–1486. 10.1111/nph.14113

Engel, S. R., Wong, E. D., Nash, R. S., Aleksander, S., Alexander, M., Douglass, E., Karra, K., Miyasato, S. R., Simison, M., Skrzypek, M. S., Weng, S., & Cherry, J. M. (2022). New data and collaborations at the Saccharomyces Genome Database: updated reference genome, alleles, and the Alliance of Genome Resources. Genetics, 220(220). 10.1093/genetics/iyab224

Gazave, E., Tassone, E. E., Ilut, D. C., Wingerson, M., Datema, E., Witsenboer, H. M. A., Davis, J. B., Grant, D., Dyer, J. M., Jenks, M. A., Brown, J., & Gore, M. A. (2016). Population genomic analysis reveals differential evolutionary histories and patterns of diversity across subgenomes and subpopulations of Brassica napus L. Frontiers in Plant Science, 7(525). 10.3389/fpls.2016.00525

Girgis, H. Z. (2015). Red: An intelligent, rapid, accurate tool for detecting repeats de-novo on the genomic scale. BMC Bioinformatics, 16(227). 10.1186/s12859-015-0654-5

Glémin, S., Scornavacca, C., Dainat, J., Burgarella, C., Viader, V., Ardisson, M., Sarah, G., Santoni, S., David, J., & Ranwez, V. (2019). Pervasive hybridizations in the history of wheat relatives. Science Advances, 5(eaav9188). 10.1126/sciadv.aav9188

Golicz, A. A., Bayer, P. E., Barker, G. C., Edger, P. P., Kim, H. R., Martinez, P. A., Chan, C. K. K., Severn-Ellis, A., McCombie, W. R., Parkin, I. A. P., Paterson, A. H., Pires, J. C., Sharpe, A. G., Tang, H., Teakle, G. R., Town, C. D., Batley, J., & Edwards, D. (2016). The pangenome of an agronomically important crop plant Brassica oleracea. Nature Communications, 7(13390). 10.1038/ncomms13390

Gordon, S. P., Contreras-Moreira, B., Levy, J. J., Djamei, A., Czedik-Eysenberg, A., Tartaglio, V. S., Session, A., Martin, J., Cartwright, A., Katz, A., Singan, V. R., Goltsman, E., Barry, K., Dinh-Thi, V. H., Chalhoub, B., Diaz-Perez, A., Sancho, R., Lusinska, J., Wolny, E., … Vogel, J. P. (2020). Gradual polyploid genome evolution revealed by pan-genomic analysis of Brachypodium hybridum and its diploid progenitors. Nature Communications, 11(1). 10.1038/s41467-020-17302-5

Gordon, S. P., Contreras-Moreira, B., Woods, D. P., Des Marais, D. L., Burgess, D., Shu, S., Stritt, C., Roulin, A. C., Schackwitz, W., Tyler, L., Martin, J., Lipzen, A., Dochy, N., Phillips, J., Barry, K., Geuten, K., Budak, H., Juenger, T. E., Amasino, R., … Vogel, J. P. (2017). Extensive gene content variation in the Brachypodium distachyon pan-genome correlates with population structure. Nature Communications, 8(2184). 10.1038/s41467-017-02292-8

Gregory, T. R., & Mable B.K. (2005). Polyploidy in Animals. In T. R. Gregory (Ed.), The Evolution of the Genome (pp. 427–517). Elsevier Inc. 10.1016/B978-0-12-301463-4.X5000-1

Halabi, K., Shafir, A., & Mayrose, I. (2023). PloiDB: the plant ploidy database. In New Phytologist (Vol. 240, Issue 3, pp. 918–927). John Wiley and Sons Inc. 10.1111/nph.19057

Jayakodi, M., Padmarasu, S., Haberer, G., Bonthala, V. S., Gundlach, H., Monat, C., Lux, T., Kamal, N., Lang, D., Himmelbach, A., Ens, J., Zhang, X. Q., Angessa, T. T., Zhou, G., Tan, C., Hill, C., Wang, P., Schreiber, M., Boston, L. B., … Stein, N. (2020). The barley pan-genome reveals the hidden legacy of mutation breeding. Nature, 588(7837), 284–289. 10.1038/s41586-020-2947-8

Jiao, W. B., & Schneeberger, K. (2020). Chromosome-level assemblies of multiple Arabidopsis genomes reveal hotspots of rearrangements with altered evolutionary dynamics. Nature Communications, 11(989). 10.1038/s41467-020-14779-y

Jiao, Y., Wickett, N. J., Ayyampalayam, S., Chanderbali, A. S., Landherr, L., Ralph, P. E., Tomsho, L. P., Hu, Y., Liang, H., Soltis, P. S., Soltis, D. E., Clifton, S. W., Schlarbaum, S. E., Schuster, S. C., Ma, H., Leebens-Mack, J., & depamphilis, C. W. (2011). Ancestral polyploidy in seed plants and angiosperms. Nature, 473(7345), 97–100. 10.1038/nature09916

Jones, G., Sagitov, S., & Oxelman, B. (2013). Statistical inference of allopolyploid species networks in the presence of incomplete lineage sorting. Systematic Biology, 62(3), 467–478. 10.1093/sysbio/syt012

Kamneva, O. K., Syring, J., Liston, A., & Rosenberg, N. A. (2017). Evaluating allopolyploid origins in strawberries (Fragaria) using haplotypes generated from target capture sequencing. BMC Evolutionary Biology, 17(180). 10.1186/s12862-017-1019-7

Kang, M., Wu, H., Liu, H., Liu, W., Zhu, M., Han, Y., Liu, W., Chen, C., Song, Y., Tan, L., Yin, K., Zhao, Y., Yan, Z., Lou, S., Zan, Y., & Liu, J. (2023). The pan-genome and local adaptation of Arabidopsis thaliana. Nature Communications, 14(6259). 10.1038/s41467-023-42029-4

Khan, A., Belfield, E. J., Harberd, N. P., & Mithani, A. (2016). HANDS2: Accurate assignment of homoeallelic base-identity in allopolyploids despite missing data. Scientific Reports, 6(29234). 10.1038/srep29234

Kong, W., Wang, Y., Zhang, S., Yu, J., & Zhang, X. (2023). Recent Advances in Assembly of Complex Plant Genomes. In *Genomics*, Proteomics and Bioinformatics (Vol. 21, Issue 3, pp. 427–439). Beijing Genomics Institute. 10.1016/j.gpb.2023.04.004

Kulkarni, R., Zhang, Y., Cannon, S. B., & Dorman, K. S. (2023). CAPG: comprehensive allopolyploid genotyper. Bioinformatics, 39(1). 10.1093/bioinformatics/btac729

Levin, & Donald A. (2002). The Role of Chromosomal Change in Plant Evolution. In D. A. Levin (Ed.), The Role of Chromosomal Change in Plant Evolution. Oxford Series in Ecology and Evolution. Oxford University Press.

Li, H. (2018). Minimap2: Pairwise alignment for nucleotide sequences. Bioinformatics, 34(18), 3094– 3100. 10.1093/bioinformatics/bty191

Li, H. (2021). New strategies to improve minimap2 alignment accuracy. Bioinformatics, 37(23), 4572– 4574. 10.1093/bioinformatics/btab705

Li, K., Xu, P., Wang, J., Yi, X., & Jiao, Y. (2023). Identification of errors in draft genome assemblies at single-nucleotide resolution for quality assessment and improvement. Nature Communications, 14(6556). 10.1038/s41467-023-42336-w

Li, L. F., Zhang, Z. Bin, Wang, Z. H., Li, N., Sha, Y., Wang, X. F., Ding, N., Li, Y., Zhao, J., Wu, Y., Gong, L., Mafessoni, F., Levy, A. A., & Liu, B. (2022). Genome sequences of five Sitopsis species of Aegilops and the origin of polyploid wheat B subgenome. Molecular Plant, 15(3), 488–503. 10.1016/j.molp.2021.12.019

Lin, H.-N., & Hsu, W.-L. (2020). GSAlign: An efficient sequence alignment tool for intra-species genomes. BMC Genomics, 21(182). 10.1186/s12864-020-6569-1

Ling, H. Q., Ma, B., Shi, X., Liu, H., Dong, L., Sun, H., Cao, Y., Gao, Q., Zheng, S., Li, Y., Yu, Y., Du, H., Qi, M., Li, Y., Lu, H., Yu, H., Cui, Y., Wang, N., Chen, C., … Liang, C. (2018). Genome sequence of the progenitor of wheat A subgenome Triticum urartu. Nature, 557(7705), 424–428. 10.1038/s41586-018-0108-0

Liu, L., Chen, M., Folk, R. A., Wang, M., Zhao, T., Shang, F., Soltis, D. E., & Li, P. (2023). Phylogenomic and syntenic data demonstrate complex evolutionary processes in early radiation of the rosids. Molecular Ecology Resources, 23, 1673–1688. 10.1111/1755-0998.13833

Lv, H., Wang, Y., Han, F., Ji, J., Fang, Z., Zhuang, M., Li, Z., Zhang, Y., & Yang, L. (2020). A high-quality reference genome for cabbage obtained with SMRT reveals novel genomic features and evolutionary characteristics. Scientific Reports, 10(12394). 10.1038/s41598-020-69389-x

Marcussen, T., Heier, L., Brysting, A. K., Oxelman, B., & Jakobsen, K. S. (2015). From gene trees to a dated allopolyploid network: Insights from the angiosperm genus viola (violaceae). Systematic Biology, 64(1), 84–101. 10.1093/sysbio/syu071

Marcussen, T., Sandve, S. R., Heier, L., Spannagl, M., Pfeifer, M., The International Wheat Genome Sequencing Consortium, Jakobsen, K. S., Wulff, B. B. H., Steuernagel, B., Mayer, K. F. X., & Olsen, O.-A. (2014). Ancient hybridizations among the ancestral genomes of bread wheat. Science, 345(6194). 10.1126/science.1251788

Mason, A. S. (2015). Challenges of Genotyping Polyploid Species. In J. Batley (Ed.), Plant Genotyping. Methods in Molecular Biology. Humana Press. 10.1007/978-1-4939-1966-6_12

Masterson, J. (1994). Stomatal Size in Fossil Plants: Evidence for Polyploidy in Majority of Angiosperms. Science, 264. http://science.sciencemag.org/

Minh, B. Q., Schmidt, H. A., Chernomor, O., Schrempf, D., Woodhams, M. D., von Haeseler, A., & Lanfear, R. (2020). IQ-TREE 2: New Models and Efficient Methods for Phylogenetic Inference in the Genomic Era. Molecular Biology and Evolution, 37(5), 1530–1534. 10.1093/molbev/msaa015

Mithani, A., Belfield, E., Brown, C., Jiang, C., Leach, L., & Harberd, N. (2013). HANDS: a tool for genome-wide discovery of subgenome-specific base-identity in polyploids. BMC Genomics, 14(653). 10.1186/1471-2164-14-653

Montenegro, J. D., Golicz, A. A., Bayer, P. E., Hurgobin, B., Lee, H. T., Chan, C. K. K., Visendi, P., Lai, K., Doležel, J., Batley, J., & Edwards, D. (2017). The pangenome of hexaploid bread wheat. Plant Journal, 90(5), 1007–1013. 10.1111/tpj.13515

Mu, W., Li, K., Yang, Y., Breiman, A., Yang, J., Wu, Y., Zhu, M., Wang, S., Catalan, P., Nevo, E., & Liu, J. (2023). Subgenomic Stability of Progenitor Genomes During Repeated Allotetraploid Origins of the Same Grass Brachypodium hybridum. Molecular Biology and Evolution, 40(12). 10.1093/molbev/msad259

Nakato, R., & Gotoh, O. (2010). Cgaln: fast and space-efficient whole-genome alignment. BMC Bioinformatics, 11(224). http://www.biomedcentral.com/1471-2105/11/224

Ning, W., Meudt, H. M., & Tate, J. A. (2024). A roadmap of phylogenomic methods for studying polyploid plant genera. Applications in Plant Sciences, 12(e11580). 10.1002/aps3.11580

Oxelman, B., Brysting, A. K., Jones, G. R., Marcussen, T., Oberprieler, C., & Pfeil, B. E. (2017). Phylogenetics of Allopolyploids. Annu. Rev. Ecol. Evol. Syst, 48, 543–557. 10.1146/annurev-ecolsys-110316-022729

Page, A. J., Sjunnebo, S., & Seemann, T. (2018). pANIto. Calculate genome wide average nucleotide identity (gwANI) for a multiFASTA alignment. (0.0.1).

Page, A. J., Taylor, B., Delaney, A. J., Soares, J., Seemann, T., Keane, J. A., & Harris, S. R. (2016). SNP-sites: rapid efficient extraction of SNPs from multi-FASTA alignments. Microbial Genomics, 2(4), e000056. 10.1099/mgen.0.000056

Page, J. T., Gingle, A. R., & Udall, J. A. (2013). PolyCat: A resource for genome categorization of sequencing reads from allopolyploid organisms. G3: Genes, Genomes, Genetics, 3(3), 517–525. 10.1534/g3.112.005298

Page, J. T., & Udall, J. A. (2015). Methods for mapping and categorization of DNA sequence reads from allopolyploid organisms. BMC Genetics, 16(Suppl 2). 10.1186/1471-2156-16-S2-S4

Parkin, I. A. P., Koh, C., Tang, H., Robinson, S. J., Kagale, S., Clarke, W. E., Town, C. D., Nixon, J., Krishnakumar, V., Bidwell, S. L., Denoeud, F., Belcram, H., Links, M. G., Just, J., Clarke, C., Bender, T., Huebert, T., Mason, A. S., Chris Pires, J., … Sharpe, A. G. (2014). Transcriptome and methylome profiling reveals relics of genome dominance in the mesopolyploid Brassica oleracea. Genome Biology, 15(6). 10.1186/gb-2014-15-6-r77

Peralta, M., Combes, M. C., Cenci, A., Lashermes, P., & Dereeper, A. (2013). SNiPloid: A utility to exploit high-throughput SNP data derived from RNA-Seq in allopolyploid species. International Journal of Plant Genomics, 2013(890123). 10.1155/2013/890123

Peris, D., Alexander, W. G., Fisher, K. J., Moriarty, R. V., Basuino, M. G., Ubbelohde, E. J., Wrobel, R. L., & Hittinger, C. T. (2020). Synthetic hybrids of six yeast species. Nature Communications, 11(2085). 10.1038/s41467-020-15559-4

Petersen, G., Seberg, O., Yde, M., & Berthelsen, K. (2006). Phylogenetic relationships of Triticum and Aegilops and evidence for the origin of the A, B, and D genomes of common wheat (Triticum aestivum). Molecular Phylogenetics and Evolution, 39(1), 70–82. 10.1016/j.ympev.2006.01.023

Phillips, A. R. (2024). Variant calling in polyploids for population and quantitative genetics. Applications in Plant Sciences, 12(4). 10.1002/aps3.11607

Procházka, E., Franko, F., Poláková, S., & Sulo, P. (2012). A complete sequence of Saccharomyces paradoxus mitochondrial genome that restores the respiration in S. cerevisiae. FEMS Yeast Research, 12(7), 819–830. 10.1111/j.1567-1364.2012.00833.x

Quinlan, A. R., & Hall, I. M. (2010). BEDTools: A flexible suite of utilities for comparing genomic features. Bioinformatics, 26(6), 841–842. 10.1093/bioinformatics/btq033

Ramsey, J., & Schemske, D. W. (1998). PATHWAYS, MECHANISMS, AND RATES OF POLYPLOID FORMATION IN FLOWERING PLANTS. Annu. Rev. Ecol. Syst, 29, 467–501. http://www.annualreviews.org;

Renny-Byfield, S., & Wendel, J. F. (2014). Doubling down on genomes: Polyploidy and crop plants. American Journal of Botany, 101(10), 1711–1725. 10.3732/ajb.1400119

Sancho, R., Cantalapiedra, C. P., López-Alvarez, D., Gordon, S. P., Vogel, J. P., Catalán, P., & Contreras-Moreira, B. (2018). Comparative plastome genomics and phylogenomics of Brachypodium: flowering time signatures, introgression and recombination in recently diverged ecotypes. New Phytologist, 218(4), 1631–1644. 10.1111/nph.14926

Sancho, R., Inda, L. A., Díaz-Pérez, A., Des Marais, D. L., Gordon, S., Vogel, J. P., Lusinska, J., Hasterok, R., Contreras-Moreira, B., & Catalán, P. (2022). Tracking the ancestry of known and ‘ghost’ homeologous subgenomes in model grass Brachypodium polyploids. Plant Journal, 109(6), 1535–1558. 10.1111/tpj.15650

Scarlett, V. T., Lovell, J. T., Shao, M., Phillips, J., Shu, S., Lusinska, J., Goodstein, D. M., Jenkins, J., Grimwood, J., Barry, K., Chalhoub, B., Schmutz, J., Hasterok, R., Catalán, P., & Vogel, J. P. (2023). Multiple origins, one evolutionary trajectory: gradual evolution characterizes distinct lineages of allotetraploid Brachypodium. Genetics, 223(2). 10.1093/genetics/iyac146

Schreiber, M., Jayakodi, M., Stein, N., & Mascher, M. (2024). Plant pangenomes for crop improvement, biodiversity and evolution. Nature Reviews Genetics, 25(8), 563–577. 10.1038/s41576-024-00691-4

Shang L, He W, Wang T, Yang Y, Xu Q, Zhao X, Yang L, Zhang H, Li X, Lv Y, Chen W, Cao S, Wang X, Zhang B, Liu X, Yu X, He H, Wei H, Leng Y, … Qian Q. (2023). A complete assembly of the rice Nipponbare reference genome. Molecular Plant, 16. 10.1016/j.molp.2023.08.00

Soltis, D. E., Albert, V. A., Leebens-Mack, J., Bell, C. D., Paterson, A. H., Zheng, C., Sankoff, D., DePamphilis, C. W., Wall, P. K., & Soltis, P. S. (2009). Polyploidy and angiosperm diversification. American Journal of Botany, 96(1), 336–348. 10.3732/ajb.0800079

Soltis, D. E., Visger, C. J., Blaine Marchant, D., & Soltis, P. S. (2016). Polyploidy: Pitfalls and paths to a paradigm. American Journal of Botany, 103(7), 1146–1166. 10.3732/ajb.1500501

Song, J.-M., Guan, Z., Hu, J., Guo, C., Yang, Z., Wang, S., Liu, D., Wang, B., Lu, S., Zhou, R., Xie, W.-Z., Cheng, Y., Zhang, Y., Liu, K., Yang, Q.-Y., Chen, L.-L., & Guo, L. (2020). Eight high-quality genomes reveal pan-genome architecture and ecotype differentiation of Brassica napus. Nature Plants, 6(1), 34–45. 10.1038/s41477-019-0577-7

Stebbins, G. L. (1949). THE EVOLUTIONARY SIGNIFICANCE OF NATURAL AND ARTIFICIAL POLYPLOIDS IN THE FAMILY GRAMINEAE. Hereditas, 35(1 S), 461–485. 10.1111/j.1601-5223.1949.tb03355.x

Than, C., & Nakhleh, L. (2009). Species tree inference by minimizing deep coalescences. PLoS Computational Biology, 5(9). 10.1371/journal.pcbi.1000501

Than, C., Ruths, D., & Nakhleh, L. (2008). PhyloNet: A software package for analyzing and reconstructing reticulate evolutionary relationships. BMC Bioinformatics, 9(322). 10.1186/1471-2105-9-322

Todd, R. T., Forche, A., & Selmecki, A. (2017). Ploidy Variation in Fungi: Polyploidy, Aneuploidy, and Genome Evolution. Microbiology Spectrum. 10.1128/microbiolspec.FUNK

Van de Peer, Y., & Meyer, A. (2005). Large-Scale Gene and Ancient Genome Duplications. In T. R. Gregory (Ed.), The Evolution of the Genome (pp. 329–368). Elsevier Inc. 10.1016/B978-0-12-301463-4.X5000-1

Vogel, J. P., Garvin, D. F., Mockler, T. C., Schmutz, J., Rokhsar, D., Bevan, M. W., Barry, K., Lucas, S., Harmon-Smith, M., Lail, K., Tice, H., Grimwood, J., McKenzie, N., Huo, N., Gu, Y. Q., Lazo, G. R., Anderson, O. D., You, F. M., Luo, M. C., … Baxter, I. (2010). Genome sequencing and analysis of the model grass Brachypodium distachyon. Nature, 463(7282), 763–768. 10.1038/nature08747

Walden, N., & Schranz, M. E. (2023). Synteny Identifies Reliable Orthologs for Phylogenomics and Comparative Genomics of the Brassicaceae. Genome Biology and Evolution, 15(3). 10.1093/gbe/evad034

Wang, L., Zhu, T., Rodriguez, J. C., Deal, K. R., Dubcovsky, J., McGuire, P. E., Lux, T., Spannagl, M., Mayer, K. F. X., Baldrich, P., Meyers, B. C., Huo, N., Gu, Y. Q., Zhou, H., Devos, K. M., Bennetzen, J. L., Unver, T., Budak, H., Gulick, P. J., … Dvorak, J. (2021). Aegilops tauschii genome assembly Aet v5.0 features greater sequence contiguity and improved annotation. G3: Genes, Genomes, Genetics, 11(12). 10.1093/g3journal/jkab325

Wang, Y., Yu, J., Jiang, M., Lei, W., Zhang, X., & Tang, H. (2023). Sequencing and Assembly of Polyploid Genomes. In Y. Van de Peer (Ed.), Polyploidy. Methods in Molecular Biology (Vol. 2545). Humana. 10.1007/978-1-0716-2561-3_23

Williams, T., Kelley, C., Bersch, C., Bröker, H.-B., Campbell, J., Cunningham, R., Denholm, D., Elber, G., Fearick, R., Grammes, C., Hart, L., Hecking, L., Juhász, P., Koenig, T., Kotz, D., Kubaitis, E., Lang, R., Lecomte, T., Lehmann, A., … Zellner, J. (2023). gnuplot 6.0 An Interactive Plotting Program. http://sourceforge.net/projects/gnuplot

Wood, T. E., Takebayashi, N., Barker, M. S., Mayrose, I., Greenspoon, P. B., & Rieseberg, L. H. (2009). The frequency of polyploid speciation in vascular plants. Proceedings of the National Academy of Sciences (PNAS), 106(33).

Yue, J. X., Li, J., Aigrain, L., Hallin, J., Persson, K., Oliver, K., Bergström, A., Coupland, P., Warringer, J., Lagomarsino, M. C., Fischer, G., Durbin, R., & Liti, G. (2017). Contrasting evolutionary genome dynamics between domesticated and wild yeasts. Nature Genetics, 49(6), 913–924. 10.1038/ng.3847

Zhang, L., Cai, X., Wu, J., Liu, M., Grob, S., Cheng, F., Liang, J., Cai, C., Liu, Z., Liu, B., Wang, F., Li, S., Liu, F., Li, X., Cheng, L., Yang, W., Li, M. he, Grossniklaus, U., Zheng, H., & Wang, X. (2018). Improved Brassica rapa reference genome by single-molecule sequencing and chromosome conformation capture technologies. Horticulture Research, 5(50). 10.1038/s41438-018-0071-9

Zhao, T., Zwaenepoel, A., Xue, J. Y., Kao, S. M., Li, Z., Schranz, M. E., & Van de Peer, Y. (2021). Whole-genome microsynteny-based phylogeny of angiosperms. Nature Communications, 12(1). 10.1038/s41467-021-23665-0

Zimin, A. V., Puiu, D., Hall, R., Kingan, S., Clavijo, B. J., & Salzberg, S. L. (2017). The first near-complete assembly of the hexaploid bread wheat genome, Triticum aestivum. GigaScience, 6(11). 10.1093/gigascience/gix097

