## Supplementary Material for "Deconvolution of Single Homeologous Polymorphism (SHP) drives phylogenetic analysis of allopolyploids"

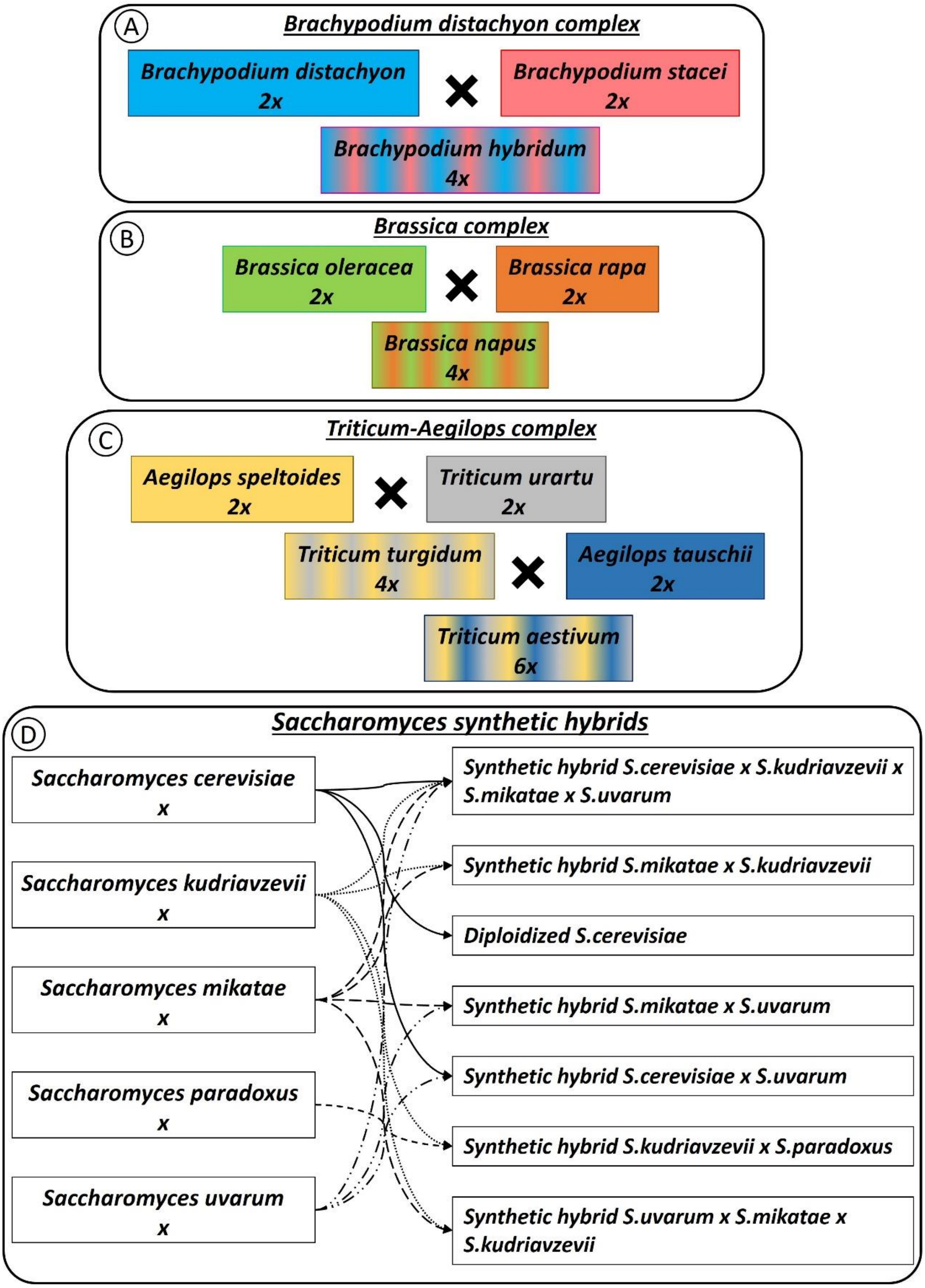


**Figure S1.** Diagram of the diploid and allopolyploid species comprising the **(A)** *Brachypodium distachyon*, **(B)** *Brassica*, **(C)** *Triticum-Aegilops*, and **(D)** *Saccharomyces* complexes used in pipeline validation.

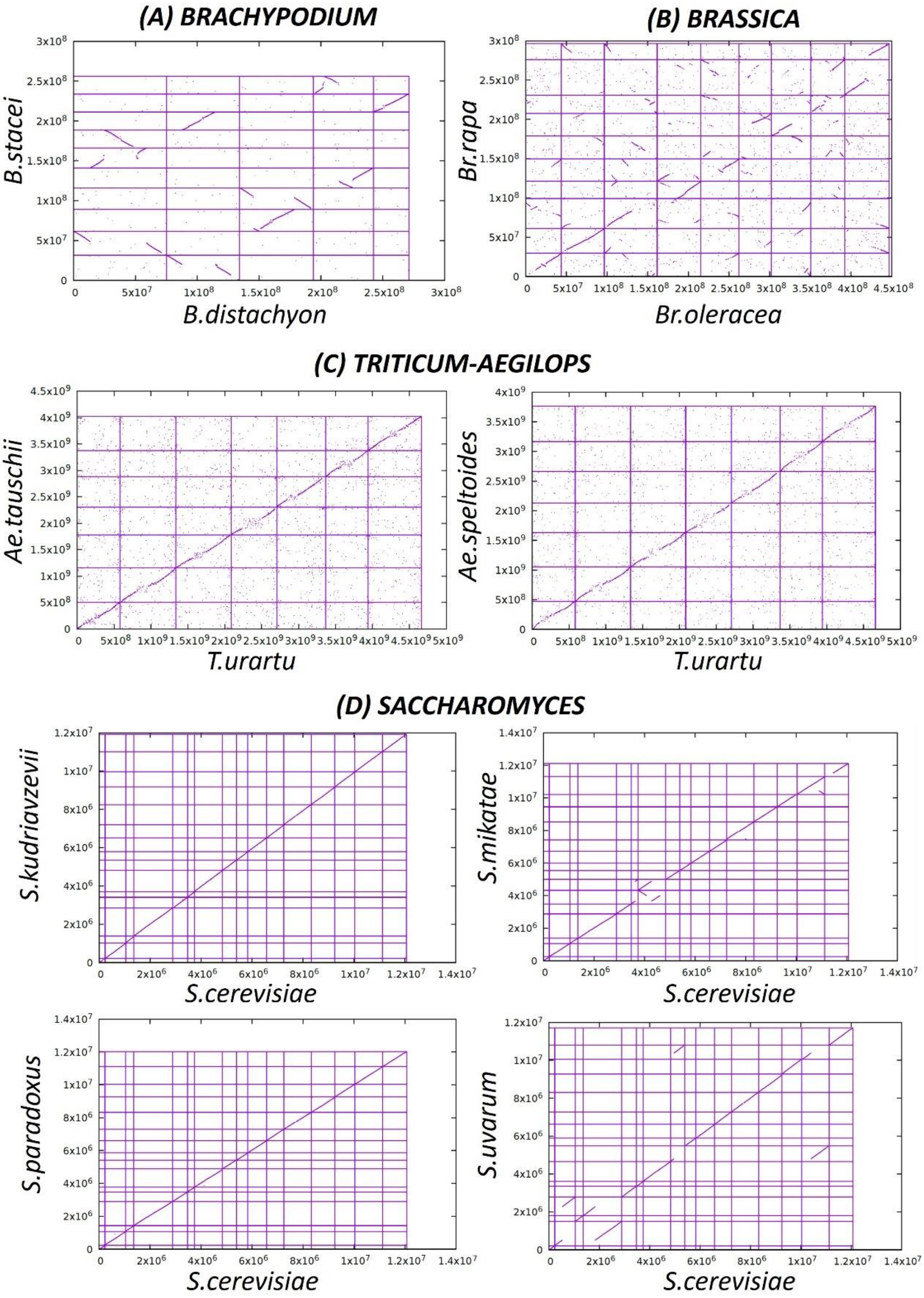


**Figure S2.** Dot plots of syntenic regions between diploid/haploid reference genomes for each diploid-polyploid complex data set were calculated using CGaln software. **(A)** *Brachypodium distachyon*, **(B)** *Brassica*, **(C)** *Triticum-Aegilops*, and **(D)** *Saccharomyces*.

**Table S1.** The list of diploid-polyploid complex species included in the pipeline validation. Diploid-polyploid complex (*Brachypodium*, *Brassica*, *Triticum-Aegilops* and *Saccharomyces*), Genome/Sample ID (species, variety, ecotype, line), Ploidy, no. chr (chromosome number), GS (genome size), Accession/file name (accession number or FASTA file name), Sample type (master or secondary reference genome, outgroup genome and diploid or allopolyploid sample) and Source (database and/or publication). Asterisk indicates to see Peris et al. (2020) for more information about ploidy and parental chromosomal contribution (for each strain, the number of chromosomes and the ploidy were estimated from the sppIDer plots).

| Diploid-polyploid complex | Genome/Sample ID | Ploidy | No. chr | GS | Accession/file name | Sample type | Source |
|---|---|---|---|---|---|---|---|
| BRACHYPODIUM | *Brachypodium distachyon* Bd21 (Reference) | 2x | 5 | 271.2 Mb | Bdistachyon_556_v3.0.fa.gz | Master Reference Genome | Phytozome13<br>IBI, 2010 |
| | *Brachypodium stacei* ABR114 (Reference) | 2x | 10 | 234.1 Mb | Brachypodium_stacei_var_ABR114.mainGenome.fasta | Secondary Reference Genome | Genome Portal<br>Catalán et al., unpublish |
| | *Oryza sativa* Japonica group cv. Nipponbare (Outgroup) | 2x | 12 | 385.7 Mb | GCF_034140825.1 | Outgroup Genome | NCBI<br>Shang et al., 2023 |
| | *Brachypodium distachyon* Bd21Control | 2x | 5 | - | 2042.3.1728.fastq.gz | diploid sample | Genome Portal |
| | *Brachypodium distachyon* ABR2 | 2x | 5 | - | 2263.4.1841.fastq.gz | diploid sample | Genome Portal |
| | *Brachypodium stacei* ABR114 | 2x | 10 | - | 2109.4.1764.fastq.gz | diploid sample | Genome Portal |
| | *Brachypodium stacei* TE4.3 | 2x | 10 | - | 7958.1.88312.GAGTGG.fastq.gz | diploid sample | Genome Portal |
| | *Brachypodium hybridum* ABR113 | 4x | 10+5 | - | 6245.6.41265.TTAGGC.anqr.fastq.gz<br>7951.2.88095.GTGGCC.anqdsp.fastq.gz<br>7951.3.88093.ACAGTG.anqdsp.fastq.gz | allopolyploid sample | Genome Portal |
| | *Brachypodium hybridum* Bhyb26 | 4x | 10+5 | - | 9505.1.135632.fastq.gz | allopolyploid sample | Genome Portal |
| BRASSICA | *Brassica oleracea* var. oleracea cv. TO1000 (Reference) | 2x | 9 | 488.6 Mb | GCF_000695525.1; BOL | Master Reference Genome | NCBI<br>Parkin et al., 2014 |
| | *Brassica rapa* cv. Chiifu-401-42 (Reference) | 2x | 10 | 352.8 Mb | GCF_000309985.2; CAAS_Brap_v3.01 | Secondary Reference Genome | NCBI<br>Zhang et al., 2018 |

| | | | | | | | |
|---|---|---|---|---|---|---|---|
| | *Brassica oleracea* var. capitata cv. OX-heart_923 | 2x | 9 | - | SRR13355052 | diploid sample | NCBI |
| | *Brassica oleracea* var. oleracea cv. TO1000 | 2x | 9 | - | SRR1212991<br>SRR1212997 | diploid sample | NCBI<br>Parkin et al., 2014 |
| | *Brassica rapa* subsp. chinensis cv. PC-fu | 2x | 10 | - | SRR16307040 | diploid sample | NCBI |
| | *Brassica rapa* subsp. trilocularis R-o-18 | 2x | 10 | - | SRR12738970<br>SRR12738972 | diploid sample | NCBI |
| | *Brassica napus* R16GE06 | 4x | 10+9 | - | SRR15524577 | allopolyploid sample | NCBI |
| | *Brassica napus* R16GE38 | 4x | 10+9 | - | SRR15524586 | allopolyploid sample | NCBI |
| | *Brassica napus* R16GE39 | 4x | 10+9 | - | SRR15524585 | allopolyploid sample | NCBI |
| | *Brassica napus* R16GE44 | 4x | 10+9 | - | SRR15524584 | allopolyploid sample | NCBI |
| | *Brassica napus* R16GE45 | 4x | 10+9 | - | SRR15524583 | allopolyploid sample | NCBI |
| TRITICUM-AEGILOPS | *Triticum urartu* cv. G1812 (Reference) | 2x | 7 | 4.8 Gb | GCF_003073215.2; Tu2.1 | Master Reference Genome | NCBI<br>Ling et al., 2018 |
| | *Aegilops tauschii* cv. AL8/78 (Reference) | 2x | 7 | 4.2 Gb | GCF_002575655.2; Aet_v5.0 | Secondary Reference Genome | NCBI<br>Wang et al., 2021 |
| | *Aegilops speltoides* isolate TS01 (Reference) | 2x | 7 | 4.1 Gb | GCA_021437245.1; ASM2143724v1 | Secondary Reference Genome | NCBI<br>Li et al., 2022 |
| | *Aegilops speltoides* isolate Y2032 | 2x | 7 | - | SRR17843190 | diploid sample | NCBI |
| | *Aegilops tauschii* isolate L4 | 2x | 7 | - | SRR14561445 | diploid sample | NCBI |
| | *Triticum urartu* cv. G1812 | 2x | 7 | - | SRR4010671<br>SRR4010672 | diploid sample | NCBI<br>Ling et al., 2018 |
| | *Triticum turgidum* cv. TD454 | 4x | 7+7 | - | SRR15681642 | allopolyploid sample | NCBI |
| | *Triticum aestivum* cv. Chinese Spring | 6x | 7+7+7 | - | SRR5817288 | allopolyploid sample | NCBI<br>Zimin et al., 2017 |
| SACCHAROMYCES | *Saccharomyces cerevisiae* R64 strain S288C | haploid | 16 | 12.1 Mb | GCF_000146045 | Master Reference Genome | NCBI<br>Engel et al., 2021 |
| | *Saccharomyces kudriavzevii* IFO1802 | haploid | 16 | 11.9 Mb | GCF_947243775 | Secondary Reference Genome | NCBI |

| | | | | | | |
|---|---|---|---|---|---|---|
| *Saccharomyces mikatae* IFO1815 | haploid | 16 | 12.1 Mb | GCF_947241705 | Secondary Reference Genome | NCBI |
| Saccharomyces *paradoxus* ASM207905v1 strain CBS432 | haploid | 16 | 12.0 Mb | GCF_002079055 | Secondary Reference Genome | NCBI<br>Yue et al., 2017<br>Procházka et al., 2012 |
| *Saccharomyces* uvarum ZP964 | haploid | 16 | 11.7 Mb | GCA_947243795 | Secondary Reference Genome | NCBI |
| *Saccharomyces cerevisiae* BY4743 | haploid | 16 | - | SRR13512344 | haploid sample | NCBI |
| *Saccharomyces kudriavzevii* CLIB 1664 | haploid | 16 | - | ERR9706976 | haploid sample | NCBI |
| *Saccharomyces mikatae* yHAB328 | haploid | 16 | - | SRR7370096 | haploid sample | NCBI |
| *Saccharomyces paradoxus* natural isolate - strain 95-7-1D | haploid | 16 | - | SRR1868557 | haploid sample | NCBI |
| *Saccharomyces uvarum* yHAB60 | haploid | 16 | - | SRR7370090 | haploid sample | NCBI |
| Synthetic hybrid *S. cerevisiae* yHRW134 x *S. kudriavzevii* ZP591 x S. mikatae IFO1815 x *S. uvarum* CBS7001 | 4.53* | * | - | SRR7769310 | synthetic hybrid sample yHRWh24 | NCBI<br>Peris et al., 2020 |
| Synthetic hybrid *S. mikatae* IFO1815 x *S. kudriavzevii* ZP591 | 3.97* | * | - | SRR7769312 | synthetic hybrid sample yHRWh4 | NCBI<br>Peris et al., 2020 |
| Diploidized *S. cerevisiae* GLBRCY101 | 2.03* | * | - | SRR7769313 | synthetic hybrid sample yHRW134 | NCBI<br>Peris et al., 2020 |
| Synthetic hybrid *S. mikatae* IFO1815 x *S. uvarum* CBS7001 | 4.00* | * | - | SRR7769316 | synthetic hybrid sample yHRWh13 | NCBI<br>Peris et al., 2020 |
| Synthetic hybrid *S.* cerevisiae yHRW134 x *S. uvarum* CBS7001 | 3.97* | * | - | SRR7769317 | synthetic hybrid sample yHRWh10 | NCBI<br>Peris et al., 2020 |
| Synthetic hybrid *S. kudriavzevii* ZP591 x *S. paradoxus* CBS432 | 4.00* | * | - | SRR7769319 | synthetic hybrid sample yHRWh18 | NCBI<br>Peris et al., 2020 |
| Synthetic hybrid *S. uvarum* CBS7001 x *S. mikatae* IFO1815 x *S. kudriavzevii* ZP591 | 5.41* | * | - | SRR7769320 | synthetic hybrid sample yHRWh51 | NCBI<br>Peris et al., 2020 |

**Table S2.** Parameters and thresholds applied to CGaln (Nakato and Gotoh, 2010) and *mapcoords* (in-house script), which are dependencies and subroutines of the WGA algorithm.

| Diploid reference genomes | | CGALN parameters | | | | | | | mapcoords parameters | |
|---|---|---|---|---|---|---|---|---|---|---|
| **Species A (Master genome)** | **Species B (Secondary genome)** | **-K (k-mer size)** | **-BS (block size)** | **-k (seed weight/width; -k1: 11/18; -k2: 12/19; -k3: 13/20 (11/18))** | **X (X drop-off at block-level)** | **-r (both strand)** | **-fc (filter colony to extract consistent set )** | **-cons (filter inconsistent HSPs at the HSP-chaining)** | **MAX MULTI BLOCK POSITIONS (max ratio of mapped positions in other syntenic blocks)** | **MAX MULTI POSITIONS (max ratio of coordinates with multiple positions in same syntenic block)** |
| *B. distachyon* | *B. stacei* | 11 | default | default | 12000 | yes | yes | yes | 0.25 | 0.05 |
| *B. distachyon* | *O. sativa* | 11 | default | default | 12000 | yes | yes | yes | 0.25 | 0.05 |
| *Br. oleracea* | *Br. rapa* | 11 | default | default | 12000 | yes | yes | yes | 0.25 | 0.05 |
| *T. urartu* | *Ae. speltoides* | 12 | 20000 | k2 | 10000 | yes | yes | yes | 0.25 | 0.05 |
| *T. urartu* | *Ae. tauchii* | 12 | 20000 | k2 | 10000 | yes | yes | yes | 0.25 | 0.05 |
| *S. cerevisiae* | *S. kudriavzevii* | 11 | default | default | default | yes | no | no | 0.25 | 0.20 |
| *S. cerevisiae* | *S. mikatae* | 11 | default | default | default | yes | no | no | 0.25 | 0.20 |
| *S. cerevisiae* | *S. paradoxus* | 11 | default | default | default | yes | no | no | 0.25 | 0.20 |
| *S. cerevisiae* | *S. uvarum* | 11 | default | default | default | yes | no | no | 0.25 | 0.20 |

**Table S3.** Statistics of the mappings of diploid and polyploid samples against the concatenated diploid reference genomes. Accession (species, ecotype, line, accession number and/or ID fasta file), Chr length (reference chromosome length in base pairs), mapped reads-segments (number of reads mapped against reference genome), % per chr (percentage of mappings per reference chromosome), % per (sub)genome (total percentage of mapping per reference genome). These statistics were calculated for each sample included in *Brachypodium* **(A)**, *Brassica* **(B)**, *Triticum-Aegilops* **(C)** and *Saccharomyces* **(D)** data sets using the samtools idxstats tool. Bold numbers indicate the predominant mappings.

**(A)**

| Accession | Reference genome and chromosome | | Chr length (bp) | mapped reads-segments | % per chr | % per (sub)genome |
|---|---|---|---|---|---|---|
| *B. distachyon* Bd21Control (Bd21Control_2042.3.1728) | *B. distachyon* | Bd1 | 75071545 | 51482338 | 29.57 | **97.81** |
| | | Bd2 | 59130575 | 35182123 | 20.20 | |
| | | Bd3 | 59640145 | 36326349 | 20.86 | |
| | | Bd4 | 48594894 | 29263299 | 16.81 | |
| | | Bd5 | 28630136 | 18070166 | 10.38 | |
| | *B. stacei* | Chr01 | 31564145 | 210478 | 0.12 | 2.19 |
| | | Chr02 | 29889700 | 147295 | 0.08 | |
| | | Chr03 | 27658146 | 1040099 | 0.60 | |
| | | Chr04 | 27050098 | 158875 | 0.09 | |
| | | Chr05 | 25022241 | 315630 | 0.18 | |
| | | Chr06 | 24930558 | 937004 | 0.54 | |
| | | Chr07 | 22499862 | 155369 | 0.09 | |
| | | Chr08 | 22836500 | 212417 | 0.12 | |
| | | Chr09 | 22451604 | 67471 | 0.04 | |
| | | Chr10 | 22136647 | 560672 | 0.32 | |
| *B. distachyon* ABR2 (ABR2_2263.4.1841) | *B. distachyon* | Bd1 | 75071545 | 56924347 | 29.85 | **97.91** |
| | | Bd2 | 59130575 | 38762627 | 20.33 | |
| | | Bd3 | 59640145 | 39899411 | 20.92 | |
| | | Bd4 | 48594894 | 31508343 | 16.52 | |
| | | Bd5 | 28630136 | 19630801 | 10.29 | |
| | *B. stacei* | Chr01 | 31564145 | 198202 | 0.10 | 2.09 |
| | | Chr02 | 29889700 | 108011 | 0.06 | |
| | | Chr03 | 27658146 | 1074054 | 0.56 | |
| | | Chr04 | 27050098 | 186492 | 0.10 | |
| | | Chr05 | 25022241 | 271389 | 0.14 | |
| | | Chr06 | 24930558 | 1100616 | 0.58 | |
| | | Chr07 | 22499862 | 130609 | 0.07 | |
| | | Chr08 | 22836500 | 179059 | 0.09 | |
| | | Chr09 | 22451604 | 113217 | 0.06 | |
| | | Chr10 | 22136647 | 623042 | 0.33 | |
| *B. stacei* ABR114 (ABR114_2109.4.1764) | *B. distachyon* | Bd1 | 75071545 | 8532314 | 2.17 | 4.47 |
| | | Bd2 | 59130575 | 1682265 | 0.43 | |

| | | | | | | |
|---|---|---|---|---|---|---|
| | | Bd3 | 59640145 | 3175533 | 0.81 | |
| | | Bd4 | 48594894 | 1909514 | 0.48 | |
| | | Bd5 | 28630136 | 2307360 | 0.59 | |
| | *B. stacei* | Chr01 | 31564145 | 46263714 | 11.74 | **95.53** |
| | | Chr02 | 29889700 | 42291393 | 10.74 | |
| | | Chr03 | 27658146 | 44608922 | 11.32 | |
| | | Chr04 | 27050098 | 38094562 | 9.67 | |
| | | Chr05 | 25022241 | 37584928 | 9.54 | |
| | | Chr06 | 24930558 | 36096711 | 9.16 | |
| | | Chr07 | 22499862 | 32015358 | 8.13 | |
| | | Chr08 | 22836500 | 33613474 | 8.53 | |
| | | Chr09 | 22451604 | 31710723 | 8.05 | |
| | | Chr10 | 22136647 | 34032211 | 8.64 | |
| *B. stacei* TE4.3 (TE4-3_7958.1.88312.GAGTGG) | *B. distachyon* | Bd1 | 75071545 | 1667824 | 4.67 | 7.76 |
| | | Bd2 | 59130575 | 282406 | 0.79 | |
| | | Bd3 | 59640145 | 310835 | 0.87 | |
| | | Bd4 | 48594894 | 218057 | 0.61 | |
| | | Bd5 | 28630136 | 291835 | 0.82 | |
| | *B. stacei* | Chr01 | 31564145 | 3987645 | 11.16 | **92.24** |
| | | Chr02 | 29889700 | 3653873 | 10.23 | |
| | | Chr03 | 27658146 | 4269030 | 11.95 | |
| | | Chr04 | 27050098 | 3254220 | 9.11 | |
| | | Chr05 | 25022241 | 3155208 | 8.83 | |
| | | Chr06 | 24930558 | 3219265 | 9.01 | |
| | | Chr07 | 22499862 | 2713225 | 7.60 | |
| | | Chr08 | 22836500 | 2936694 | 8.22 | |
| | | Chr09 | 22451604 | 2568150 | 7.19 | |
| | | Chr10 | 22136647 | 3190489 | 8.93 | |
| *B. hybridum* ABR113 (6245.6_plus_7951.2_plus_7951.3) | *B. distachyon* | Bd1 | 75071545 | 16132607 | 15.98 | **50.27** |
| | | Bd2 | 59130575 | 10339808 | 10.24 | |
| | | Bd3 | 59640145 | 10791176 | 10.69 | |
| | | Bd4 | 48594894 | 8186710 | 8.11 | |
| | | Bd5 | 28630136 | 5294625 | 5.24 | |
| | *B. stacei* | Chr01 | 31564145 | 5935439 | 5.88 | **49.73** |
| | | Chr02 | 29889700 | 5486397 | 5.43 | |
| | | Chr03 | 27658146 | 6976472 | 6.91 | |
| | | Chr04 | 27050098 | 4722519 | 4.68 | |
| | | Chr05 | 25022241 | 4807522 | 4.76 | |
| | | Chr06 | 24930558 | 4932027 | 4.89 | |
| | | Chr07 | 22499862 | 3971261 | 3.93 | |
| | | Chr08 | 22836500 | 4184907 | 4.15 | |
| | | Chr09 | 22451604 | 3727397 | 3.69 | |
| | | Chr10 | 22136647 | 5465444 | 5.41 | |
| *B. hybridum* Bhyb26 (Bhyb26_9505.1.135632) | *B. distachyon* | Bd1 | 75071545 | 18197278 | 16.89 | **54.13** |
| | | Bd2 | 59130575 | 11770784 | 10.92 | |

| | | | | | | |
|---|---|---|---|---|---|---|
| | | Bd3 | 59640145 | 12474059 | 11.58 | |
| | | Bd4 | 48594894 | 9433671 | 8.76 | |
| | | Bd5 | 28630136 | 6447086 | 5.98 | |
| | *B. stacei* | Chr01 | 31564145 | 6185402 | 5.74 | **45.87** |
| | | Chr02 | 29889700 | 5740492 | 5.33 | |
| | | Chr03 | 27658146 | 5871201 | 5.45 | |
| | | Chr04 | 27050098 | 5074989 | 4.71 | |
| | | Chr05 | 25022241 | 4631210 | 4.30 | |
| | | Chr06 | 24930558 | 4830288 | 4.48 | |
| | | Chr07 | 22499862 | 4419623 | 4.10 | |
| | | Chr08 | 22836500 | 4303209 | 3.99 | |
| | | Chr09 | 22451604 | 4266843 | 3.96 | |
| | | Chr10 | 22136647 | 4095608 | 3.80 | |

**(B)**

| Accession | Reference genome and chromosome | | Chr length (bp) | mapped reads-segments | % per chr | % per (sub)genome |
|---|---|---|---|---|---|---|
| *Br. oleracea* var. Capitata (SRR13355052) | *Br. oleracea* | Bro1 | 43764888 | 36764426 | 8.08 | **83.68** |
| | | Bro2 | 52886895 | 44833070 | 9.85 | |
| | | Bro3 | 64984695 | 55253573 | 12.14 | |
| | | Bro4 | 53719093 | 47061369 | 10.34 | |
| | | Bro5 | 46902585 | 40325888 | 8.86 | |
| | | Bro6 | 39822476 | 32126587 | 7.06 | |
| | | Bro7 | 48366697 | 43906713 | 9.65 | |
| | | Bro8 | 41758685 | 35647618 | 7.83 | |
| | | Bro9 | 54679868 | 44952066 | 9.88 | |
| | *Br. rapa* | Brr01 | 29595527 | 4975212 | 1.09 | 16.32 |
| | | Brr02 | 31442979 | 3892330 | 0.86 | |
| | | Brr03 | 38154160 | 9216553 | 2.02 | |
| | | Brr04 | 21928416 | 4530212 | 1.00 | |
| | | Brr05 | 28493056 | 4386835 | 0.96 | |
| | | Brr06 | 29167992 | 20039719 | 4.40 | |
| | | Brr07 | 28928902 | 4257595 | 0.94 | |
| | | Brr08 | 22981702 | 3552318 | 0.78 | |
| | | Brr09 | 45156810 | 4654146 | 1.02 | |
| | | Brr10 | 20725698 | 14777161 | 3.25 | |
| *Br. oleracea* var. Oleracea (SRR1212991; SRR1212997) | *Br. oleracea* | Bro1 | 43764888 | 17482438 | 7.16 | **78.08** |
| | | Bro2 | 52886895 | 21197362 | 8.68 | |
| | | Bro3 | 64984695 | 27665047 | 11.33 | |
| | | Bro4 | 53719093 | 23048871 | 9.44 | |
| | | Bro5 | 46902585 | 22384538 | 9.17 | |
| | | Bro6 | 39822476 | 14841083 | 6.08 | |
| | | Bro7 | 48366697 | 23417201 | 9.59 | |

| | | | | | | |
|---|---|---|---|---|---|---|
| | | Bro8 | 41758685 | 17178658 | 7.04 | |
| | | Bro9 | 54679868 | 23405487 | 9.59 | |
| | *Br. rapa* | Brr01 | 29595527 | 1759505 | 0.72 | 21.92 |
| | | Brr02 | 31442979 | 970600 | 0.40 | |
| | | Brr03 | 38154160 | 3473138 | 1.42 | |
| | | Brr04 | 21928416 | 1550528 | 0.64 | |
| | | Brr05 | 28493056 | 1380030 | 0.57 | |
| | | Brr06 | 29167992 | 33459745 | 13.71 | |
| | | Brr07 | 28928902 | 1591356 | 0.65 | |
| | | Brr08 | 22981702 | 1181189 | 0.48 | |
| | | Brr09 | 45156810 | 1428999 | 0.59 | |
| | | Brr10 | 20725698 | 6721318 | 2.75 | |
| *Br. rapa* var. Chinensis (SRR16307040) | *Br. oleracea* | Bro1 | 43764888 | 2233905 | 0.91 | 11.53 |
| | | Bro2 | 52886895 | 2801085 | 1.15 | |
| | | Bro3 | 64984695 | 4790831 | 1.96 | |
| | | Bro4 | 53719093 | 2246221 | 0.92 | |
| | | Bro5 | 46902585 | 4229343 | 1.73 | |
| | | Bro6 | 39822476 | 1393627 | 0.57 | |
| | | Bro7 | 48366697 | 5203820 | 2.13 | |
| | | Bro8 | 41758685 | 1983608 | 0.81 | |
| | | Bro9 | 54679868 | 3309244 | 1.35 | |
| | *Br. rapa* | Brr01 | 29595527 | 18237168 | 7.46 | **88.47** |
| | | Brr02 | 31442979 | 14098042 | 5.76 | |
| | | Brr03 | 38154160 | 36347659 | 14.86 | |
| | | Brr04 | 21928416 | 15652674 | 6.40 | |
| | | Brr05 | 28493056 | 18515644 | 7.57 | |
| | | Brr06 | 29167992 | 42591191 | 17.42 | |
| | | Brr07 | 28928902 | 16816376 | 6.88 | |
| | | Brr08 | 22981702 | 12903682 | 5.28 | |
| | | Brr09 | 45156810 | 23055071 | 9.43 | |
| | | Brr10 | 20725698 | 18139337 | 7.42 | |
| *Br. rapa* var. Trilocularis (SRR12738970; SRR12738972) | *Br. oleracea* | Bro1 | 43764888 | 797888 | 1.14 | 12.57 |
| | | Bro2 | 52886895 | 1101149 | 1.57 | |
| | | Bro3 | 64984695 | 1501214 | 2.14 | |
| | | Bro4 | 53719093 | 841788 | 1.20 | |
| | | Bro5 | 46902585 | 976813 | 1.39 | |
| | | Bro6 | 39822476 | 538492 | 0.77 | |
| | | Bro7 | 48366697 | 1362477 | 1.94 | |
| | | Bro8 | 41758685 | 767732 | 1.09 | |
| | | Bro9 | 54679868 | 939270 | 1.34 | |
| | *Br. rapa* | Brr01 | 29595527 | 6073722 | 8.65 | **87.43** |
| | | Brr02 | 31442979 | 4914490 | 7.00 | |
| | | Brr03 | 38154160 | 10974793 | 15.63 | |
| | | Brr04 | 21928416 | 4621749 | 6.58 | |
| | | Brr05 | 28493056 | 5124176 | 7.30 | |

| | | | | | | |
|---|---|---|---|---|---|---|
| | | Brr06 | 29167992 | 7324860 | 10.43 | |
| | | Brr07 | 28928902 | 4991416 | 7.11 | |
| | | Brr08 | 22981702 | 4574843 | 6.51 | |
| | | Brr09 | 45156810 | 6851779 | 9.76 | |
| | | Brr10 | 20725698 | 5944428 | 8.47 | |
| *Br. napus* R16GE06 (SRR15524577) | *Br. oleracea* | Bro1 | 43764888 | 2983638 | 4.76 | **49.89** |
| | | Bro2 | 52886895 | 3755177 | 5.99 | |
| | | Bro3 | 64984695 | 4653662 | 7.42 | |
| | | Bro4 | 53719093 | 3894831 | 6.21 | |
| | | Bro5 | 46902585 | 3485870 | 5.56 | |
| | | Bro6 | 39822476 | 2176956 | 3.47 | |
| | | Bro7 | 48366697 | 3774330 | 6.02 | |
| | | Bro8 | 41758685 | 2895356 | 4.62 | |
| | | Bro9 | 54679868 | 3668246 | 5.85 | |
| | *Br. rapa* | Brr01 | 29595527 | 2507599 | 4.00 | **50.11** |
| | | Brr02 | 31442979 | 2101629 | 3.35 | |
| | | Brr03 | 38154160 | 4678356 | 7.46 | |
| | | Brr04 | 21928416 | 1905009 | 3.04 | |
| | | Brr05 | 28493056 | 2371768 | 3.78 | |
| | | Brr06 | 29167992 | 6593085 | 10.51 | |
| | | Brr07 | 28928902 | 2597935 | 4.14 | |
| | | Brr08 | 22981702 | 1907851 | 3.04 | |
| | | Brr09 | 45156810 | 3083610 | 4.92 | |
| | | Brr10 | 20725698 | 3676043 | 5.86 | |
| *Br. napus* R16GE38 (SRR15524586) | *Br. oleracea* | Bro1 | 43764888 | 2928754 | 4.53 | **47.39** |
| | | Bro2 | 52886895 | 3371279 | 5.21 | |
| | | Bro3 | 64984695 | 4500515 | 6.96 | |
| | | Bro4 | 53719093 | 3690770 | 5.71 | |
| | | Bro5 | 46902585 | 3574664 | 5.53 | |
| | | Bro6 | 39822476 | 2418725 | 3.74 | |
| | | Bro7 | 48366697 | 3830431 | 5.92 | |
| | | Bro8 | 41758685 | 2681951 | 4.15 | |
| | | Bro9 | 54679868 | 3658369 | 5.66 | |
| | *Br. rapa* | Brr01 | 29595527 | 2225069 | 3.44 | **52.61** |
| | | Brr02 | 31442979 | 2264360 | 3.50 | |
| | | Brr03 | 38154160 | 5468175 | 8.45 | |
| | | Brr04 | 21928416 | 1904049 | 2.94 | |
| | | Brr05 | 28493056 | 2366024 | 3.66 | |
| | | Brr06 | 29167992 | 9469299 | 14.64 | |
| | | Brr07 | 28928902 | 2162653 | 3.34 | |
| | | Brr08 | 22981702 | 1816154 | 2.81 | |
| | | Brr09 | 45156810 | 2971728 | 4.59 | |
| | | Brr10 | 20725698 | 3385855 | 5.23 | |
| *Br. napus* R16GE39 (SRR15524585) | *Br. oleracea* | Bro1 | 43764888 | 3106657 | 4.41 | **49.22** |
| | | Bro2 | 52886895 | 3665752 | 5.21 | |

| | | | | | | |
|---|---|---|---|---|---|---|
| | | Bro3 | 64984695 | 5058945 | 7.19 | |
| | | Bro4 | 53719093 | 4162502 | 5.91 | |
| | | Bro5 | 46902585 | 4141814 | 5.88 | |
| | | Bro6 | 39822476 | 2804247 | 3.98 | |
| | | Bro7 | 48366697 | 4260509 | 6.05 | |
| | | Bro8 | 41758685 | 3185445 | 4.52 | |
| | | Bro9 | 54679868 | 4265672 | 6.06 | |
| | *Br. rapa* | Brr01 | 29595527 | 2855331 | 4.06 | **50.78** |
| | | Brr02 | 31442979 | 2747888 | 3.90 | |
| | | Brr03 | 38154160 | 4343285 | 6.17 | |
| | | Brr04 | 21928416 | 2241261 | 3.18 | |
| | | Brr05 | 28493056 | 2266484 | 3.22 | |
| | | Brr06 | 29167992 | 9323269 | 13.24 | |
| | | Brr07 | 28928902 | 2383438 | 3.39 | |
| | | Brr08 | 22981702 | 2134651 | 3.03 | |
| | | Brr09 | 45156810 | 3311544 | 4.70 | |
| | | Brr10 | 20725698 | 4148887 | 5.89 | |
| *Br. napus* R16GE44 (SRR15524584) | *Br. oleracea* | Bro1 | 43764888 | 2777855 | 4.68 | **49.13** |
| | | Bro2 | 52886895 | 3400424 | 5.73 | |
| | | Bro3 | 64984695 | 4152555 | 7.00 | |
| | | Bro4 | 53719093 | 3509944 | 5.92 | |
| | | Bro5 | 46902585 | 3365465 | 5.67 | |
| | | Bro6 | 39822476 | 2339276 | 3.94 | |
| | | Bro7 | 48366697 | 3508092 | 5.91 | |
| | | Bro8 | 41758685 | 2658770 | 4.48 | |
| | | Bro9 | 54679868 | 3439200 | 5.80 | |
| | *Br. rapa* | Brr01 | 29595527 | 2325698 | 3.92 | **50.87** |
| | | Brr02 | 31442979 | 2142751 | 3.61 | |
| | | Brr03 | 38154160 | 4798594 | 8.09 | |
| | | Brr04 | 21928416 | 1701225 | 2.87 | |
| | | Brr05 | 28493056 | 2176387 | 3.67 | |
| | | Brr06 | 29167992 | 6954957 | 11.72 | |
| | | Brr07 | 28928902 | 2165620 | 3.65 | |
| | | Brr08 | 22981702 | 1895669 | 3.19 | |
| | | Brr09 | 45156810 | 2816137 | 4.75 | |
| | | Brr10 | 20725698 | 3204306 | 5.40 | |
| *Br. napus* R16GE45 (SRR15524583) | *Br. oleracea* | Bro1 | 43764888 | 3426505 | 4.79 | **49.07** |
| | | Bro2 | 52886895 | 3695005 | 5.16 | |
| | | Bro3 | 64984695 | 5301393 | 7.40 | |
| | | Bro4 | 53719093 | 4136355 | 5.78 | |
| | | Bro5 | 46902585 | 3978372 | 5.56 | |
| | | Bro6 | 39822476 | 3010448 | 4.20 | |
| | | Bro7 | 48366697 | 4187152 | 5.85 | |
| | | Bro8 | 41758685 | 3323711 | 4.64 | |
| | | Bro9 | 54679868 | 4079041 | 5.70 | |

| | | | | | |
|---|---|---|---|---|---|
| | Br. rapa | Brr01 | 29595527 | 2874850 | 4.01 | **50.93** |
| | | Brr02 | 31442979 | 3049291 | 4.26 | |
| | | Brr03 | 38154160 | 6218883 | 8.68 | |
| | | Brr04 | 21928416 | 2418078 | 3.38 | |
| | | Brr05 | 28493056 | 2819102 | 3.94 | |
| | | Brr06 | 29167992 | 6746156 | 9.42 | |
| | | Brr07 | 28928902 | 2684540 | 3.75 | |
| | | Brr08 | 22981702 | 2227251 | 3.11 | |
| | | Brr09 | 45156810 | 3748986 | 5.24 | |
| | | Brr10 | 20725698 | 3682364 | 5.14 | |

**(C)**

| Accession | Reference genome and chromosome | | Chr length (bp) | mapped reads-segments | % per chr | % per (sub)genome |
|---|---|---|---|---|---|---|
| *Ae. speltoides* (SRR17843190) | *T. urartu* | Tru1 | 584211917 | 6863295 | 0.76 | 5.60 |
| | | Tru2 | 753719114 | 8211601 | 0.91 | |
| | | Tru3 | 747046674 | 7643136 | 0.85 | |
| | | Tru4 | 619584425 | 6150445 | 0.68 | |
| | | Tru5 | 661480603 | 7247326 | 0.81 | |
| | | Tru6 | 575865107 | 5467017 | 0.61 | |
| | | Tru7 | 719689837 | 8690855 | 0.97 | |
| | *Ae. tauschii* | Aet1 | 501967303 | 9645870 | 1.07 | 7.95 |
| | | Aet2 | 650458083 | 12153297 | 1.35 | |
| | | Aet3 | 627456150 | 11240367 | 1.25 | |
| | | Aet4 | 525206139 | 8551441 | 0.95 | |
| | | Aet5 | 576238907 | 10876622 | 1.21 | |
| | | Aet6 | 495363004 | 7854518 | 0.87 | |
| | | Aet7 | 644841383 | 11136648 | 1.24 | |
| | *Ae. speltoides* | Aes1 | 470101627 | 99488032 | 11.07 | **86.45** |
| | | Aes2 | 583316874 | 116275550 | 12.94 | |
| | | Aes3 | 578812325 | 110903521 | 12.34 | |
| | | Aes4 | 498717206 | 98077932 | 10.92 | |
| | | Aes5 | 530918953 | 111836171 | 12.45 | |
| | | Aes6 | 505192780 | 94204678 | 10.49 | |
| | | Aes7 | 597334302 | 145909013 | 16.24 | |
| *Ae. tauschii* (SRR14561445) | *T. urartu* | Tru1 | 584211917 | 5733793 | 0.31 | 2.23 |
| | | Tru2 | 753719114 | 7061227 | 0.38 | |
| | | Tru3 | 747046674 | 6354836 | 0.34 | |
| | | Tru4 | 619584425 | 4854345 | 0.26 | |
| | | Tru5 | 661480603 | 6201754 | 0.33 | |
| | | Tru6 | 575865107 | 4207822 | 0.22 | |
| | | Tru7 | 719689837 | 7353420 | 0.39 | |
| | *Ae. tauschii* | Aet1 | 501967303 | 221470523 | 11.81 | **96.71** |

| | | | | | | |
|---|---|---|---|---|---|---|
| | | Aet2 | 650458083 | 294254762 | 15.69 | |
| | | Aet3 | 627456150 | 284216228 | 15.15 | |
| | | Aet4 | 525206139 | 241651901 | 12.88 | |
| | | Aet5 | 576238907 | 259476044 | 13.83 | |
| | | Aet6 | 495363004 | 225956775 | 12.05 | |
| | | Aet7 | 644841383 | 287019369 | 15.30 | |
| | *Ae. speltoides* | Aes1 | 470101627 | 3787421 | 0.20 | 1.06 |
| | | Aes2 | 583316874 | 2749697 | 0.15 | |
| | | Aes3 | 578812325 | 2576222 | 0.14 | |
| | | Aes4 | 498717206 | 2226572 | 0.12 | |
| | | Aes5 | 530918953 | 2384694 | 0.13 | |
| | | Aes6 | 505192780 | 1986444 | 0.11 | |
| | | Aes7 | 597334302 | 4201149 | 0.22 | |
| *T. urartu* (SRR4010671; SRR4010672) | *T. urartu* | Tru1 | 584211917 | 4617310 | 11.44 | **93.29** |
| | | Tru2 | 753719114 | 6069550 | 15.04 | |
| | | Tru3 | 747046674 | 6042307 | 14.97 | |
| | | Tru4 | 619584425 | 4914750 | 12.18 | |
| | | Tru5 | 661480603 | 5456639 | 13.52 | |
| | | Tru6 | 575865107 | 4427956 | 10.97 | |
| | | Tru7 | 719689837 | 6121148 | 15.17 | |
| | *Ae. tauschii* | Aet1 | 501967303 | 413141 | 1.02 | 4.73 |
| | | Aet2 | 650458083 | 317936 | 0.79 | |
| | | Aet3 | 627456150 | 237847 | 0.59 | |
| | | Aet4 | 525206139 | 192950 | 0.48 | |
| | | Aet5 | 576238907 | 308275 | 0.76 | |
| | | Aet6 | 495363004 | 162883 | 0.40 | |
| | | Aet7 | 644841383 | 274890 | 0.68 | |
| | *Ae. speltoides* | Aes1 | 470101627 | 61869 | 0.15 | 1.99 |
| | | Aes2 | 583316874 | 59103 | 0.15 | |
| | | Aes3 | 578812325 | 50859 | 0.13 | |
| | | Aes4 | 498717206 | 40661 | 0.10 | |
| | | Aes5 | 530918953 | 105908 | 0.26 | |
| | | Aes6 | 505192780 | 42773 | 0.11 | |
| | | Aes7 | 597334302 | 440122 | 1.09 | |
| *T. turgidum* (SRR15681642) | *T. urartu* | Tru1 | 584211917 | 50664844 | 5.80 | **46.34** |
| | | Tru2 | 753719114 | 64896271 | 7.43 | |
| | | Tru3 | 747046674 | 62767172 | 7.19 | |
| | | Tru4 | 619584425 | 54235557 | 6.21 | |
| | | Tru5 | 661480603 | 59291586 | 6.79 | |
| | | Tru6 | 575865107 | 49041974 | 5.62 | |
| | | Tru7 | 719689837 | 63694234 | 7.30 | |
| | *Ae. tauschii* | Aet1 | 501967303 | 17036675 | 1.95 | 13.87 |
| | | Aet2 | 650458083 | 21043743 | 2.41 | |
| | | Aet3 | 627456150 | 18258213 | 2.09 | |
| | | Aet4 | 525206139 | 14714642 | 1.69 | |

| | | | | | | |
|---|---|---|---|---|---|---|
| | | Aet5 | 576238907 | 18383729 | 2.11 | |
| | | Aet6 | 495363004 | 13434856 | 1.54 | |
| | | Aet7 | 644841383 | 18189835 | 2.08 | |
| | *Ae. speltoides* | Aes1 | 470101627 | 44549099 | 5.10 | **39.79** |
| | | Aes2 | 583316874 | 51657425 | 5.92 | |
| | | Aes3 | 578812325 | 52018705 | 5.96 | |
| | | Aes4 | 498717206 | 45559207 | 5.22 | |
| | | Aes5 | 530918953 | 50296821 | 5.76 | |
| | | Aes6 | 505192780 | 47022688 | 5.39 | |
| | | Aes7 | 597334302 | 56296249 | 6.45 | |
| *T. aestivum* (SRR5817288) | *T. urartu* | Tru1 | 584211917 | 33425386 | 4.31 | **34.16** |
| | | Tru2 | 753719114 | 42216326 | 5.45 | |
| | | Tru3 | 747046674 | 40924897 | 5.28 | |
| | | Tru4 | 619584425 | 35629776 | 4.60 | |
| | | Tru5 | 661480603 | 38717108 | 5.00 | |
| | | Tru6 | 575865107 | 32092560 | 4.14 | |
| | | Tru7 | 719689837 | 41687288 | 5.38 | |
| | *Ae. tauschii* | Aet1 | 501967303 | 36071283 | 4.65 | **36.64** |
| | | Aet2 | 650458083 | 47224042 | 6.09 | |
| | | Aet3 | 627456150 | 42985172 | 5.55 | |
| | | Aet4 | 525206139 | 37181689 | 4.80 | |
| | | Aet5 | 576238907 | 41520585 | 5.36 | |
| | | Aet6 | 495363004 | 34213593 | 4.42 | |
| | | Aet7 | 644841383 | 44752786 | 5.78 | |
| | *Ae. speltoides* | Aes1 | 470101627 | 29149315 | 3.76 | **29.20** |
| | | Aes2 | 583316874 | 33450551 | 4.32 | |
| | | Aes3 | 578812325 | 33965072 | 4.38 | |
| | | Aes4 | 498717206 | 29370227 | 3.79 | |
| | | Aes5 | 530918953 | 32603396 | 4.21 | |
| | | Aes6 | 505192780 | 30218265 | 3.90 | |
| | | Aes7 | 597334302 | 37502092 | 4.84 | |

**(D)**

| **Sample** | **Reference genome and chromosome** | | **Chr length (bp)** | **mapped reads-segments** | **% per chr** | **% per (sub)genome** |
|---|---|---|---|---|---|---|
| *S. kudriavzevii* CLIB 1664 (ERR9706976) | *S. cerevisiae* | Sce01 | 230218 | 91 | 0.00 | 0.94 |
| | | Sce02 | 813184 | 190 | 0.00 | |
| | | Sce03 | 316620 | 223 | 0.00 | |
| | | Sce04 | 1531933 | 206 | 0.00 | |
| | | Sce05 | 576874 | 884 | 0.02 | |
| | | Sce06 | 270161 | 48 | 0.00 | |
| | | Sce07 | 1090940 | 1005 | 0.02 | |
| | | Sce08 | 562643 | 289 | 0.01 | |

| | | | | | |
|---|---|---|---|---|---|
| | Sce09 | 439888 | 1173 | 0.02 | |
| | Sce10 | 745751 | 43 | 0.00 | |
| | Sce11 | 666816 | 442 | 0.01 | |
| | Sce12 | 1078177 | 41752 | 0.83 | |
| | Sce13 | 924431 | 114 | 0.00 | |
| | Sce14 | 784333 | 729 | 0.01 | |
| | Sce15 | 1091291 | 45 | 0.00 | |
| | Sce16 | 948066 | 222 | 0.00 | |
| *S. kudriavzevii* | Sku01 | 216969 | 95130 | 1.88 | **97.70** |
| | Sku02 | 806230 | 318528 | 6.30 | |
| | Sku03 | 355935 | 151745 | 3.00 | |
| | Sku04 | 1470946 | 564469 | 11.17 | |
| | Sku05 | 556891 | 229203 | 4.54 | |
| | Sku06 | 288445 | 122293 | 2.42 | |
| | Sku07 | 1119750 | 424466 | 8.40 | |
| | Sku08 | 523467 | 212046 | 4.20 | |
| | Sku09 | 456381 | 185780 | 3.68 | |
| | Sku10 | 719910 | 289465 | 5.73 | |
| | Sku11 | 686475 | 265733 | 5.26 | |
| | Sku12 | 1040685 | 637703 | 12.62 | |
| | Sku13 | 921530 | 362420 | 7.17 | |
| | Sku14 | 797769 | 320984 | 6.35 | |
| | Sku15 | 1050702 | 408480 | 8.09 | |
| | Sku16 | 901691 | 347730 | 6.88 | |
| *S. mikatae* | Smi01 | 248052 | 3008 | 0.06 | 0.62 |
| | Smi02 | 811203 | 153 | 0.00 | |
| | Smi03 | 322525 | 305 | 0.01 | |
| | Smi04 | 1495645 | 156 | 0.00 | |
| | Smi05 | 599206 | 60 | 0.00 | |
| | Smi06 | 848074 | 190 | 0.00 | |
| | Smi07 | 663440 | 1606 | 0.03 | |
| | Smi08 | 542523 | 126 | 0.00 | |
| | Smi09 | 462568 | 155 | 0.00 | |
| | Smi10 | 734496 | 91 | 0.00 | |
| | Smi11 | 687171 | 83 | 0.00 | |
| | Smi12 | 1104771 | 24342 | 0.48 | |
| | Smi13 | 932466 | 266 | 0.01 | |
| | Smi14 | 753128 | 108 | 0.00 | |
| | Smi15 | 1095438 | 371 | 0.01 | |
| | Smi16 | 804165 | 160 | 0.00 | |
| *S. paradoxus* | Spa01 | 235665 | 128 | 0.00 | 0.09 |
| | Spa02 | 835812 | 249 | 0.00 | |
| | Spa03 | 357685 | 115 | 0.00 | |

| | | | | | | |
|---|---|---|---|---|---|---|
| | | Spa04 | 1477988 | 179 | 0.00 | |
| | | Spa05 | 565208 | 103 | 0.00 | |
| | | Spa06 | 296034 | 165 | 0.00 | |
| | | Spa07 | 1119193 | 785 | 0.02 | |
| | | Spa08 | 531267 | 286 | 0.01 | |
| | | Spa09 | 443128 | 263 | 0.01 | |
| | | Spa10 | 743843 | 272 | 0.01 | |
| | | Spa11 | 690553 | 281 | 0.01 | |
| | | Spa12 | 1023116 | 300 | 0.01 | |
| | | Spa13 | 941477 | 340 | 0.01 | |
| | | Spa14 | 778926 | 57 | 0.00 | |
| | | Spa15 | 1078405 | 806 | 0.02 | |
| | | Spa16 | 903028 | 67 | 0.00 | |
| | *S. uvarum* | Suv01 | 204763 | 23 | 0.00 | 0.65 |
| | | Suv02 | 1292445 | 163 | 0.00 | |
| | | Suv03 | 301324 | 166 | 0.00 | |
| | | Suv04 | 991278 | 70 | 0.00 | |
| | | Suv05 | 564469 | 22 | 0.00 | |
| | | Suv06 | 258855 | 19 | 0.00 | |
| | | Suv07 | 1040537 | 194 | 0.00 | |
| | | Suv08 | 834793 | 29 | 0.00 | |
| | | Suv09 | 409318 | 36 | 0.00 | |
| | | Suv10 | 731806 | 108 | 0.00 | |
| | | Suv11 | 643462 | 34 | 0.00 | |
| | | Suv12 | 1035508 | 31472 | 0.62 | |
| | | Suv13 | 967206 | 69 | 0.00 | |
| | | Suv14 | 773076 | 123 | 0.00 | |
| | | Suv15 | 745522 | 198 | 0.00 | |
| | | Suv16 | 920918 | 215 | 0.00 | |
| *S. cerevisiae* BY4743 (SRR13512344) | *S. cerevisiae* | Sce01 | 230218 | 10612 | 1.77 | **97.01** |
| | | Sce02 | 813184 | 36834 | 6.15 | |
| | | Sce03 | 316620 | 13913 | 2.32 | |
| | | Sce04 | 1531933 | 69659 | 11.62 | |
| | | Sce05 | 576874 | 26207 | 4.37 | |
| | | Sce06 | 270161 | 11976 | 2.00 | |
| | | Sce07 | 1090940 | 49441 | 8.25 | |
| | | Sce08 | 562643 | 26442 | 4.41 | |
| | | Sce09 | 439888 | 20089 | 3.35 | |
| | | Sce10 | 745751 | 34148 | 5.70 | |
| | | Sce11 | 666816 | 30230 | 5.04 | |
| | | Sce12 | 1078177 | 82302 | 13.73 | |
| | | Sce13 | 924431 | 42076 | 7.02 | |
| | | Sce14 | 784333 | 35105 | 5.86 | |

| | | | | | |
|---|---|---|---|---|---|
| | Sce15 | 1091291 | 49640 | 8.28 | |
| | Sce16 | 948066 | 42666 | 7.12 | |
| *S. kudriavzevii* | Sku01 | 216969 | 0 | 0.00 | 1.23 |
| | Sku02 | 806230 | 9 | 0.00 | |
| | Sku03 | 355935 | 12 | 0.00 | |
| | Sku04 | 1470946 | 158 | 0.03 | |
| | Sku05 | 556891 | 0 | 0.00 | |
| | Sku06 | 288445 | 54 | 0.01 | |
| | Sku07 | 1119750 | 102 | 0.02 | |
| | Sku08 | 523467 | 55 | 0.01 | |
| | Sku09 | 456381 | 2 | 0.00 | |
| | Sku10 | 719910 | 29 | 0.00 | |
| | Sku11 | 686475 | 103 | 0.02 | |
| | Sku12 | 1040685 | 6830 | 1.14 | |
| | Sku13 | 921530 | 1 | 0.00 | |
| | Sku14 | 797769 | 0 | 0.00 | |
| | Sku15 | 1050702 | 6 | 0.00 | |
| | Sku16 | 901691 | 3 | 0.00 | |
| *S. mikatae* | Smi01 | 248052 | 468 | 0.08 | 0.83 |
| | Smi02 | 811203 | 8 | 0.00 | |
| | Smi03 | 322525 | 5 | 0.00 | |
| | Smi04 | 1495645 | 13 | 0.00 | |
| | Smi05 | 599206 | 67 | 0.01 | |
| | Smi06 | 848074 | 2 | 0.00 | |
| | Smi07 | 663440 | 32 | 0.01 | |
| | Smi08 | 542523 | 208 | 0.03 | |
| | Smi09 | 462568 | 0 | 0.00 | |
| | Smi10 | 734496 | 2 | 0.00 | |
| | Smi11 | 687171 | 95 | 0.02 | |
| | Smi12 | 1104771 | 3526 | 0.59 | |
| | Smi13 | 932466 | 40 | 0.01 | |
| | Smi14 | 753128 | 0 | 0.00 | |
| | Smi15 | 1095438 | 499 | 0.08 | |
| | Smi16 | 804165 | 11 | 0.00 | |
| *S. paradoxus* | Spa01 | 235665 | 0 | 0.00 | 0.22 |
| | Spa02 | 835812 | 106 | 0.02 | |
| | Spa03 | 357685 | 60 | 0.01 | |
| | Spa04 | 1477988 | 180 | 0.03 | |
| | Spa05 | 565208 | 7 | 0.00 | |
| | Spa06 | 296034 | 38 | 0.01 | |
| | Spa07 | 1119193 | 241 | 0.04 | |
| | Spa08 | 531267 | 98 | 0.02 | |
| | Spa09 | 443128 | 41 | 0.01 | |

| | | | | | | |
|---|---|---|---|---|---|---|
| | | Spa10 | 743843 | 51 | 0.01 | |
| | | Spa11 | 690553 | 2 | 0.00 | |
| | | Spa12 | 1023116 | 21 | 0.00 | |
| | | Spa13 | 941477 | 9 | 0.00 | |
| | | Spa14 | 778926 | 174 | 0.03 | |
| | | Spa15 | 1078405 | 305 | 0.05 | |
| | | Spa16 | 903028 | 8 | 0.00 | |
| | *S. uvarum* | Suv01 | 204763 | 1 | 0.00 | 0.71 |
| | | Suv02 | 1292445 | 3 | 0.00 | |
| | | Suv03 | 301324 | 10 | 0.00 | |
| | | Suv04 | 991278 | 49 | 0.01 | |
| | | Suv05 | 564469 | 1 | 0.00 | |
| | | Suv06 | 258855 | 43 | 0.01 | |
| | | Suv07 | 1040537 | 37 | 0.01 | |
| | | Suv08 | 834793 | 0 | 0.00 | |
| | | Suv09 | 409318 | 6 | 0.00 | |
| | | Suv10 | 731806 | 2 | 0.00 | |
| | | Suv11 | 643462 | 23 | 0.00 | |
| | | Suv12 | 1035508 | 3866 | 0.65 | |
| | | Suv13 | 967206 | 99 | 0.02 | |
| | | Suv14 | 773076 | 0 | 0.00 | |
| | | Suv15 | 745522 | 104 | 0.02 | |
| | | Suv16 | 920918 | 12 | 0.00 | |
| *S. paradoxus* natural isolate - strain 95-7-1D (SRR1868557) | *S. cerevisiae* | Sce01 | 230218 | 3617 | 0.21 | 4.97 |
| | | Sce02 | 813184 | 1336 | 0.08 | |
| | | Sce03 | 316620 | 601 | 0.04 | |
| | | Sce04 | 1531933 | 4890 | 0.29 | |
| | | Sce05 | 576874 | 1413 | 0.08 | |
| | | Sce06 | 270161 | 365 | 0.02 | |
| | | Sce07 | 1090940 | 2119 | 0.12 | |
| | | Sce08 | 562643 | 844 | 0.05 | |
| | | Sce09 | 439888 | 809 | 0.05 | |
| | | Sce10 | 745751 | 1174 | 0.07 | |
| | | Sce11 | 666816 | 1824 | 0.11 | |
| | | Sce12 | 1078177 | 60103 | 3.51 | |
| | | Sce13 | 924431 | 1566 | 0.09 | |
| | | Sce14 | 784333 | 1551 | 0.09 | |
| | | Sce15 | 1091291 | 1567 | 0.09 | |
| | | Sce16 | 948066 | 1382 | 0.08 | |
| | *S. kudriavzevii* | Sku01 | 216969 | 179 | 0.01 | 2.37 |
| | | Sku02 | 806230 | 79 | 0.00 | |
| | | Sku03 | 355935 | 217 | 0.01 | |
| | | Sku04 | 1470946 | 342 | 0.02 | |

| | | | | | |
|---|---|---|---|---|---|
| | Sku05 | 556891 | 93 | 0.01 | |
| | Sku06 | 288445 | 87 | 0.01 | |
| | Sku07 | 1119750 | 420 | 0.02 | |
| | Sku08 | 523467 | 212 | 0.01 | |
| | Sku09 | 456381 | 165 | 0.01 | |
| | Sku10 | 719910 | 242 | 0.01 | |
| | Sku11 | 686475 | 340 | 0.02 | |
| | Sku12 | 1040685 | 36757 | 2.14 | |
| | Sku13 | 921530 | 680 | 0.04 | |
| | Sku14 | 797769 | 373 | 0.02 | |
| | Sku15 | 1050702 | 259 | 0.02 | |
| | Sku16 | 901691 | 221 | 0.01 | |
| *S. mikatae* | Smi01 | 248052 | 1967 | 0.11 | 2.11 |
| | Smi02 | 811203 | 306 | 0.02 | |
| | Smi03 | 322525 | 161 | 0.01 | |
| | Smi04 | 1495645 | 375 | 0.02 | |
| | Smi05 | 599206 | 306 | 0.02 | |
| | Smi06 | 848074 | 253 | 0.01 | |
| | Smi07 | 663440 | 315 | 0.02 | |
| | Smi08 | 542523 | 233 | 0.01 | |
| | Smi09 | 462568 | 233 | 0.01 | |
| | Smi10 | 734496 | 820 | 0.05 | |
| | Smi11 | 687171 | 189 | 0.01 | |
| | Smi12 | 1104771 | 28185 | 1.64 | |
| | Smi13 | 932466 | 250 | 0.01 | |
| | Smi14 | 753128 | 341 | 0.02 | |
| | Smi15 | 1095438 | 2038 | 0.12 | |
| | Smi16 | 804165 | 216 | 0.01 | |
| *S. paradoxus* | Spa01 | 235665 | 25592 | 1.49 | **89.43** |
| | Spa02 | 835812 | 104585 | 6.10 | |
| | Spa03 | 357685 | 37670 | 2.20 | |
| | Spa04 | 1477988 | 198320 | 11.57 | |
| | Spa05 | 565208 | 69915 | 4.08 | |
| | Spa06 | 296034 | 32127 | 1.87 | |
| | Spa07 | 1119193 | 140561 | 8.20 | |
| | Spa08 | 531267 | 67229 | 3.92 | |
| | Spa09 | 443128 | 53899 | 3.14 | |
| | Spa10 | 743843 | 93840 | 5.47 | |
| | Spa11 | 690553 | 87835 | 5.12 | |
| | Spa12 | 1023116 | 134977 | 7.87 | |
| | Spa13 | 941477 | 122337 | 7.14 | |
| | Spa14 | 778926 | 100580 | 5.87 | |
| | Spa15 | 1078405 | 141274 | 8.24 | |

| | | | | | | |
|---|---|---|---|---|---|---|
| | | Spa16 | 903028 | 122043 | 7.12 | |
| | *S. uvarum* | Suv01 | 204763 | 15 | 0.00 | 1.12 |
| | | Suv02 | 1292445 | 140 | 0.01 | |
| | | Suv03 | 301324 | 63 | 0.00 | |
| | | Suv04 | 991278 | 68 | 0.00 | |
| | | Suv05 | 564469 | 153 | 0.01 | |
| | | Suv06 | 258855 | 31 | 0.00 | |
| | | Suv07 | 1040537 | 158 | 0.01 | |
| | | Suv08 | 834793 | 173 | 0.01 | |
| | | Suv09 | 409318 | 41 | 0.00 | |
| | | Suv10 | 731806 | 49 | 0.00 | |
| | | Suv11 | 643462 | 58 | 0.00 | |
| | | Suv12 | 1035508 | 16498 | 0.96 | |
| | | Suv13 | 967206 | 373 | 0.02 | |
| | | Suv14 | 773076 | 83 | 0.00 | |
| | | Suv15 | 745522 | 123 | 0.01 | |
| | | Suv16 | 920918 | 1202 | 0.07 | |
| *S. uvarum* yHAB60 (SRR7370090) | *S. cerevisiae* | Sce01 | 230218 | 1 | 0.00 | 0.20 |
| | | Sce02 | 813184 | 27 | 0.01 | |
| | | Sce03 | 316620 | 6 | 0.00 | |
| | | Sce04 | 1531933 | 1 | 0.00 | |
| | | Sce05 | 576874 | 12 | 0.00 | |
| | | Sce06 | 270161 | 0 | 0.00 | |
| | | Sce07 | 1090940 | 100 | 0.04 | |
| | | Sce08 | 562643 | 3 | 0.00 | |
| | | Sce09 | 439888 | 4 | 0.00 | |
| | | Sce10 | 745751 | 8 | 0.00 | |
| | | Sce11 | 666816 | 9 | 0.00 | |
| | | Sce12 | 1078177 | 320 | 0.12 | |
| | | Sce13 | 924431 | 1 | 0.00 | |
| | | Sce14 | 784333 | 7 | 0.00 | |
| | | Sce15 | 1091291 | 6 | 0.00 | |
| | | Sce16 | 948066 | 9 | 0.00 | |
| | *S. kudriavzevii* | Sku01 | 216969 | 68 | 0.03 | 0.39 |
| | | Sku02 | 806230 | 14 | 0.01 | |
| | | Sku03 | 355935 | 8 | 0.00 | |
| | | Sku04 | 1470946 | 46 | 0.02 | |
| | | Sku05 | 556891 | 20 | 0.01 | |
| | | Sku06 | 288445 | 16 | 0.01 | |
| | | Sku07 | 1119750 | 73 | 0.03 | |
| | | Sku08 | 523467 | 21 | 0.01 | |
| | | Sku09 | 456381 | 56 | 0.02 | |
| | | Sku10 | 719910 | 16 | 0.01 | |

| | | | | | |
|---|---|---|---|---|---|
| | Sku11 | 686475 | 24 | 0.01 | |
| | Sku12 | 1040685 | 396 | 0.15 | |
| | Sku13 | 921530 | 18 | 0.01 | |
| | Sku14 | 797769 | 149 | 0.06 | |
| | Sku15 | 1050702 | 47 | 0.02 | |
| | Sku16 | 901691 | 49 | 0.02 | |
| *S. mikatae* | Smi01 | 248052 | 260 | 0.10 | 0.32 |
| | Smi02 | 811203 | 6 | 0.00 | |
| | Smi03 | 322525 | 28 | 0.01 | |
| | Smi04 | 1495645 | 68 | 0.03 | |
| | Smi05 | 599206 | 29 | 0.01 | |
| | Smi06 | 848074 | 40 | 0.02 | |
| | Smi07 | 663440 | 20 | 0.01 | |
| | Smi08 | 542523 | 13 | 0.00 | |
| | Smi09 | 462568 | 4 | 0.00 | |
| | Smi10 | 734496 | 11 | 0.00 | |
| | Smi11 | 687171 | 15 | 0.01 | |
| | Smi12 | 1104771 | 271 | 0.10 | |
| | Smi13 | 932466 | 9 | 0.00 | |
| | Smi14 | 753128 | 10 | 0.00 | |
| | Smi15 | 1095438 | 54 | 0.02 | |
| | Smi16 | 804165 | 16 | 0.01 | |
| *S. paradoxus* | Spa01 | 235665 | 30 | 0.01 | 0.20 |
| | Spa02 | 835812 | 84 | 0.03 | |
| | Spa03 | 357685 | 18 | 0.01 | |
| | Spa04 | 1477988 | 22 | 0.01 | |
| | Spa05 | 565208 | 76 | 0.03 | |
| | Spa06 | 296034 | 2 | 0.00 | |
| | Spa07 | 1119193 | 34 | 0.01 | |
| | Spa08 | 531267 | 21 | 0.01 | |
| | Spa09 | 443128 | 12 | 0.00 | |
| | Spa10 | 743843 | 32 | 0.01 | |
| | Spa11 | 690553 | 41 | 0.02 | |
| | Spa12 | 1023116 | 36 | 0.01 | |
| | Spa13 | 941477 | 59 | 0.02 | |
| | Spa14 | 778926 | 7 | 0.00 | |
| | Spa15 | 1078405 | 55 | 0.02 | |
| | Spa16 | 903028 | 10 | 0.00 | |
| *S. uvarum* | Suv01 | 204763 | 3703 | 1.41 | **98.89** |
| | Suv02 | 1292445 | 28384 | 10.77 | |
| | Suv03 | 301324 | 7188 | 2.73 | |
| | Suv04 | 991278 | 21214 | 8.05 | |
| | Suv05 | 564469 | 11339 | 4.30 | |

| | | | | | | |
|---|---|---|---|---|---|---|
| | | Suv06 | 258855 | 5319 | 2.02 | |
| | | Suv07 | 1040537 | 22969 | 8.72 | |
| | | Suv08 | 834793 | 17116 | 6.50 | |
| | | Suv09 | 409318 | 9009 | 3.42 | |
| | | Suv10 | 731806 | 17385 | 6.60 | |
| | | Suv11 | 643462 | 15594 | 5.92 | |
| | | Suv12 | 1035508 | 27811 | 10.56 | |
| | | Suv13 | 967206 | 20114 | 7.63 | |
| | | Suv14 | 773076 | 16426 | 6.23 | |
| | | Suv15 | 745522 | 15986 | 6.07 | |
| | | Suv16 | 920918 | 20979 | 7.96 | |
| *S. mikatae* yHAB328 (SRR7370096) | *S. cerevisiae* | Sce01 | 230218 | 128 | 0.01 | 1.13 |
| | | Sce02 | 813184 | 29 | 0.00 | |
| | | Sce03 | 316620 | 1149 | 0.05 | |
| | | Sce04 | 1531933 | 217 | 0.01 | |
| | | Sce05 | 576874 | 90 | 0.00 | |
| | | Sce06 | 270161 | 7 | 0.00 | |
| | | Sce07 | 1090940 | 1403 | 0.06 | |
| | | Sce08 | 562643 | 28 | 0.00 | |
| | | Sce09 | 439888 | 825 | 0.03 | |
| | | Sce10 | 745751 | 24 | 0.00 | |
| | | Sce11 | 666816 | 337 | 0.01 | |
| | | Sce12 | 1078177 | 23836 | 0.95 | |
| | | Sce13 | 924431 | 24 | 0.00 | |
| | | Sce14 | 784333 | 31 | 0.00 | |
| | | Sce15 | 1091291 | 20 | 0.00 | |
| | | Sce16 | 948066 | 26 | 0.00 | |
| | *S. kudriavzevii* | Sku01 | 216969 | 20 | 0.00 | 1.23 |
| | | Sku02 | 806230 | 461 | 0.02 | |
| | | Sku03 | 355935 | 100 | 0.00 | |
| | | Sku04 | 1470946 | 37 | 0.00 | |
| | | Sku05 | 556891 | 31 | 0.00 | |
| | | Sku06 | 288445 | 4 | 0.00 | |
| | | Sku07 | 1119750 | 71 | 0.00 | |
| | | Sku08 | 523467 | 19 | 0.00 | |
| | | Sku09 | 456381 | 28 | 0.00 | |
| | | Sku10 | 719910 | 26 | 0.00 | |
| | | Sku11 | 686475 | 244 | 0.01 | |
| | | Sku12 | 1040685 | 28034 | 1.12 | |
| | | Sku13 | 921530 | 37 | 0.00 | |
| | | Sku14 | 797769 | 1412 | 0.06 | |
| | | Sku15 | 1050702 | 107 | 0.00 | |
| | | Sku16 | 901691 | 16 | 0.00 | |

| | | | | | |
|---|---|---|---|---|---|
| *S. mikatae* | Smi01 | 248052 | 60060 | 2.40 | **96.96** |
| | Smi02 | 811203 | 157558 | 6.30 | |
| | Smi03 | 322525 | 69559 | 2.78 | |
| | Smi04 | 1495645 | 270228 | 10.81 | |
| | Smi05 | 599206 | 119185 | 4.77 | |
| | Smi06 | 848074 | 158225 | 6.33 | |
| | Smi07 | 663440 | 130384 | 5.22 | |
| | Smi08 | 542523 | 107559 | 4.30 | |
| | Smi09 | 462568 | 95542 | 3.82 | |
| | Smi10 | 734496 | 144141 | 5.77 | |
| | Smi11 | 687171 | 132482 | 5.30 | |
| | Smi12 | 1104771 | 308084 | 12.32 | |
| | Smi13 | 932466 | 176278 | 7.05 | |
| | Smi14 | 753128 | 138891 | 5.56 | |
| | Smi15 | 1095438 | 201064 | 8.04 | |
| | Smi16 | 804165 | 154502 | 6.18 | |
| *S. paradoxus* | Spa01 | 235665 | 34 | 0.00 | 0.08 |
| | Spa02 | 835812 | 37 | 0.00 | |
| | Spa03 | 357685 | 87 | 0.00 | |
| | Spa04 | 1477988 | 71 | 0.00 | |
| | Spa05 | 565208 | 33 | 0.00 | |
| | Spa06 | 296034 | 8 | 0.00 | |
| | Spa07 | 1119193 | 659 | 0.03 | |
| | Spa08 | 531267 | 17 | 0.00 | |
| | Spa09 | 443128 | 39 | 0.00 | |
| | Spa10 | 743843 | 173 | 0.01 | |
| | Spa11 | 690553 | 122 | 0.00 | |
| | Spa12 | 1023116 | 28 | 0.00 | |
| | Spa13 | 941477 | 32 | 0.00 | |
| | Spa14 | 778926 | 16 | 0.00 | |
| | Spa15 | 1078405 | 665 | 0.03 | |
| | Spa16 | 903028 | 29 | 0.00 | |
| *S. uvarum* | Suv01 | 204763 | 27 | 0.00 | 0.60 |
| | Suv02 | 1292445 | 51 | 0.00 | |
| | Suv03 | 301324 | 49 | 0.00 | |
| | Suv04 | 991278 | 314 | 0.01 | |
| | Suv05 | 564469 | 29 | 0.00 | |
| | Suv06 | 258855 | 7 | 0.00 | |
| | Suv07 | 1040537 | 28 | 0.00 | |
| | Suv08 | 834793 | 38 | 0.00 | |
| | Suv09 | 409318 | 35 | 0.00 | |
| | Suv10 | 731806 | 97 | 0.00 | |
| | Suv11 | 643462 | 9 | 0.00 | |

| | | | | | | |
|---|---|---|---|---|---|---|
| | | Suv12 | 1035508 | 14166 | 0.57 | |
| | | Suv13 | 967206 | 198 | 0.01 | |
| | | Suv14 | 773076 | 23 | 0.00 | |
| | | Suv15 | 745522 | 32 | 0.00 | |
| | | Suv16 | 920918 | 18 | 0.00 | |
| Synthetic hybrid *S. cerevisiae* yHRW134 x *S. kudriavzevii* ZP591 x *S. mikatae* IFO1815 x *S. uvarum* CBS7001 (SRR7769310) | *S. cerevisiae* | Sce01 | 230218 | 16816 | 0.77 | **11.84** |
| | | Sce02 | 813184 | 959 | 0.04 | |
| | | Sce03 | 316620 | 4805 | 0.22 | |
| | | Sce04 | 1531933 | 1362 | 0.06 | |
| | | Sce05 | 576874 | 780 | 0.04 | |
| | | Sce06 | 270161 | 11346 | 0.52 | |
| | | Sce07 | 1090940 | 44413 | 2.03 | |
| | | Sce08 | 562643 | 2064 | 0.09 | |
| | | Sce09 | 439888 | 18587 | 0.85 | |
| | | Sce10 | 745751 | 630 | 0.03 | |
| | | Sce11 | 666816 | 28090 | 1.29 | |
| | | Sce12 | 1078177 | 14402 | 0.66 | |
| | | Sce13 | 924431 | 38083 | 1.74 | |
| | | Sce14 | 784333 | 31105 | 1.42 | |
| | | Sce15 | 1091291 | 44130 | 2.02 | |
| | | Sce16 | 948066 | 1073 | 0.05 | |
| | *S. kudriavzevii* | Sku01 | 216969 | 16677 | 0.76 | **37.49** |
| | | Sku02 | 806230 | 34242 | 1.57 | |
| | | Sku03 | 355935 | 23465 | 1.07 | |
| | | Sku04 | 1470946 | 115684 | 5.30 | |
| | | Sku05 | 556891 | 23583 | 1.08 | |
| | | Sku06 | 288445 | 12342 | 0.57 | |
| | | Sku07 | 1119750 | 85222 | 3.90 | |
| | | Sku08 | 523467 | 21662 | 0.99 | |
| | | Sku09 | 456381 | 36479 | 1.67 | |
| | | Sku10 | 719910 | 56999 | 2.61 | |
| | | Sku11 | 686475 | 54042 | 2.47 | |
| | | Sku12 | 1040685 | 121759 | 5.58 | |
| | | Sku13 | 921530 | 72264 | 3.31 | |
| | | Sku14 | 797769 | 32585 | 1.49 | |
| | | Sku15 | 1050702 | 42799 | 1.96 | |
| | | Sku16 | 901691 | 68938 | 3.16 | |
| | *S. mikatae* | Smi01 | 248052 | 12127 | 0.56 | **39.81** |
| | | Smi02 | 811203 | 78577 | 3.60 | |
| | | Smi03 | 322525 | 29306 | 1.34 | |
| | | Smi04 | 1495645 | 65235 | 2.99 | |
| | | Smi05 | 599206 | 51911 | 2.38 | |
| | | Smi06 | 848074 | 68589 | 3.14 | |

| | | | | | |
|---|---|---|---|---|---|
| | Smi07 | 663440 | 56180 | 2.57 | |
| | Smi08 | 542523 | 46744 | 2.14 | |
| | Smi09 | 462568 | 41318 | 1.89 | |
| | Smi10 | 734496 | 62640 | 2.87 | |
| | Smi11 | 687171 | 58686 | 2.69 | |
| | Smi12 | 1104771 | 108629 | 4.97 | |
| | Smi13 | 932466 | 40271 | 1.84 | |
| | Smi14 | 753128 | 32546 | 1.49 | |
| | Smi15 | 1095438 | 48667 | 2.23 | |
| | Smi16 | 804165 | 67813 | 3.11 | |
| | Spa01 | 235665 | 116 | 0.01 | |
| | Spa02 | 835812 | 219 | 0.01 | |
| | Spa03 | 357685 | 265 | 0.01 | |
| | Spa04 | 1477988 | 89 | 0.00 | |
| | Spa05 | 565208 | 115 | 0.01 | |
| | Spa06 | 296034 | 322 | 0.01 | |
| | Spa07 | 1119193 | 195 | 0.01 | |
| *S. paradoxus* | Spa08 | 531267 | 105 | 0.00 | 0.14 |
| | Spa09 | 443128 | 154 | 0.01 | |
| | Spa10 | 743843 | 171 | 0.01 | |
| | Spa11 | 690553 | 145 | 0.01 | |
| | Spa12 | 1023116 | 179 | 0.01 | |
| | Spa13 | 941477 | 113 | 0.01 | |
| | Spa14 | 778926 | 407 | 0.02 | |
| | Spa15 | 1078405 | 458 | 0.02 | |
| | Spa16 | 903028 | 91 | 0.00 | |
| | Suv01 | 204763 | 4896 | 0.22 | |
| | Suv02 | 1292445 | 565 | 0.03 | |
| | Suv03 | 301324 | 5630 | 0.26 | |
| | Suv04 | 991278 | 26059 | 1.19 | |
| | Suv05 | 564469 | 14440 | 0.66 | |
| | Suv06 | 258855 | 5200 | 0.24 | |
| | Suv07 | 1040537 | 27364 | 1.25 | |
| *S. uvarum* | Suv08 | 834793 | 21435 | 0.98 | **10.71** |
| | Suv09 | 409318 | 173 | 0.01 | |
| | Suv10 | 731806 | 9378 | 0.43 | |
| | Suv11 | 643462 | 145 | 0.01 | |
| | Suv12 | 1035508 | 54304 | 2.49 | |
| | Suv13 | 967206 | 437 | 0.02 | |
| | Suv14 | 773076 | 180 | 0.01 | |
| | Suv15 | 745522 | 39502 | 1.81 | |
| | Suv16 | 920918 | 24153 | 1.11 | |
| *S. cerevisiae* | Sce01 | 230218 | 21 | 0.00 | 0.86 |

| | | | | | | |
|---|---|---|---|---|---|---|
| | | Sce02 | 813184 | 126 | 0.01 | |
| | | Sce03 | 316620 | 410 | 0.02 | |
| | | Sce04 | 1531933 | 289 | 0.02 | |
| | | Sce05 | 576874 | 149 | 0.01 | |
| | | Sce06 | 270161 | 24 | 0.00 | |
| | | Sce07 | 1090940 | 723 | 0.04 | |
| | | Sce08 | 562643 | 88 | 0.00 | |
| | | Sce09 | 439888 | 319 | 0.02 | |
| | | Sce10 | 745751 | 51 | 0.00 | |
| | | Sce11 | 666816 | 913 | 0.05 | |
| | | Sce12 | 1078177 | 12188 | 0.65 | |
| | | Sce13 | 924431 | 671 | 0.04 | |
| | | Sce14 | 784333 | 122 | 0.01 | |
| | | Sce15 | 1091291 | 47 | 0.00 | |
| | | Sce16 | 948066 | 77 | 0.00 | |
| Synthetic hybrid *S. mikatae* IFO1815 x *S. kudriavzevii* ZP591 (SRR7769312) | *S. kudriavzevii* | Sku01 | 216969 | 16434 | 0.87 | **47.54** |
| | | Sku02 | 806230 | 57492 | 3.05 | |
| | | Sku03 | 355935 | 27279 | 1.45 | |
| | | Sku04 | 1470946 | 100868 | 5.35 | |
| | | Sku05 | 556891 | 42507 | 2.26 | |
| | | Sku06 | 288445 | 23079 | 1.23 | |
| | | Sku07 | 1119750 | 77284 | 4.10 | |
| | | Sku08 | 523467 | 39936 | 2.12 | |
| | | Sku09 | 456381 | 33993 | 1.80 | |
| | | Sku10 | 719910 | 53037 | 2.82 | |
| | | Sku11 | 686475 | 50185 | 2.66 | |
| | | Sku12 | 1040685 | 114045 | 6.05 | |
| | | Sku13 | 921530 | 64443 | 3.42 | |
| | | Sku14 | 797769 | 57977 | 3.08 | |
| | | Sku15 | 1050702 | 73860 | 3.92 | |
| | | Sku16 | 901691 | 63105 | 3.35 | |
| | *S. mikatae* | Smi01 | 248052 | 23480 | 1.25 | **50.87** |
| | | Smi02 | 811203 | 62837 | 3.34 | |
| | | Smi03 | 322525 | 23270 | 1.24 | |
| | | Smi04 | 1495645 | 107877 | 5.73 | |
| | | Smi05 | 599206 | 46952 | 2.49 | |
| | | Smi06 | 848074 | 63790 | 3.39 | |
| | | Smi07 | 663440 | 52547 | 2.79 | |
| | | Smi08 | 542523 | 43795 | 2.33 | |
| | | Smi09 | 462568 | 38816 | 2.06 | |
| | | Smi10 | 734496 | 57943 | 3.08 | |
| | | Smi11 | 687171 | 53817 | 2.86 | |
| | | Smi12 | 1104771 | 113085 | 6.00 | |

| | | | | | | |
|---|---|---|---|---|---|---|
| | | Smi13 | 932466 | 70177 | 3.73 | |
| | | Smi14 | 753128 | 57338 | 3.04 | |
| | | Smi15 | 1095438 | 81386 | 4.32 | |
| | | Smi16 | 804165 | 61114 | 3.24 | |
| | *S. paradoxus* | Spa01 | 235665 | 42 | 0.00 | 0.18 |
| | | Spa02 | 835812 | 78 | 0.00 | |
| | | Spa03 | 357685 | 48 | 0.00 | |
| | | Spa04 | 1477988 | 46 | 0.00 | |
| | | Spa05 | 565208 | 83 | 0.00 | |
| | | Spa06 | 296034 | 18 | 0.00 | |
| | | Spa07 | 1119193 | 855 | 0.05 | |
| | | Spa08 | 531267 | 447 | 0.02 | |
| | | Spa09 | 443128 | 88 | 0.00 | |
| | | Spa10 | 743843 | 69 | 0.00 | |
| | | Spa11 | 690553 | 233 | 0.01 | |
| | | Spa12 | 1023116 | 142 | 0.01 | |
| | | Spa13 | 941477 | 532 | 0.03 | |
| | | Spa14 | 778926 | 26 | 0.00 | |
| | | Spa15 | 1078405 | 610 | 0.03 | |
| | | Spa16 | 903028 | 34 | 0.00 | |
| | *S. uvarum* | Suv01 | 204763 | 14 | 0.00 | 0.55 |
| | | Suv02 | 1292445 | 77 | 0.00 | |
| | | Suv03 | 301324 | 19 | 0.00 | |
| | | Suv04 | 991278 | 78 | 0.00 | |
| | | Suv05 | 564469 | 16 | 0.00 | |
| | | Suv06 | 258855 | 6 | 0.00 | |
| | | Suv07 | 1040537 | 513 | 0.03 | |
| | | Suv08 | 834793 | 14 | 0.00 | |
| | | Suv09 | 409318 | 35 | 0.00 | |
| | | Suv10 | 731806 | 41 | 0.00 | |
| | | Suv11 | 643462 | 28 | 0.00 | |
| | | Suv12 | 1035508 | 9181 | 0.49 | |
| | | Suv13 | 967206 | 26 | 0.00 | |
| | | Suv14 | 773076 | 35 | 0.00 | |
| | | Suv15 | 745522 | 202 | 0.01 | |
| | | Suv16 | 920918 | 27 | 0.00 | |
| Diploidized *S. cerevisiae* GLBRCY101 (SRR7769313) | *S. cerevisiae* | Sce01 | 230218 | 32990 | 1.83 | **95.73** |
| | | Sce02 | 813184 | 112185 | 6.23 | |
| | | Sce03 | 316620 | 39337 | 2.18 | |
| | | Sce04 | 1531933 | 186658 | 10.36 | |
| | | Sce05 | 576874 | 81686 | 4.53 | |
| | | Sce06 | 270161 | 42203 | 2.34 | |
| | | Sce07 | 1090940 | 145628 | 8.08 | |

| | | | | | |
|---|---|---|---|---|---|
| | | Sce08 | 562643 | 81270 | 4.51 | |
| | | Sce09 | 439888 | 66914 | 3.71 | |
| | | Sce10 | 745751 | 103438 | 5.74 | |
| | | Sce11 | 666816 | 92517 | 5.14 | |
| | | Sce12 | 1078177 | 239858 | 13.32 | |
| | | Sce13 | 924431 | 125505 | 6.97 | |
| | | Sce14 | 784333 | 111592 | 6.19 | |
| | | Sce15 | 1091291 | 136313 | 7.57 | |
| | | Sce16 | 948066 | 126251 | 7.01 | |
| | *S. kudriavzevii* | Sku01 | 216969 | 8 | 0.00 | 1.61 |
| | | Sku02 | 806230 | 63 | 0.00 | |
| | | Sku03 | 355935 | 105 | 0.01 | |
| | | Sku04 | 1470946 | 518 | 0.03 | |
| | | Sku05 | 556891 | 10 | 0.00 | |
| | | Sku06 | 288445 | 266 | 0.01 | |
| | | Sku07 | 1119750 | 234 | 0.01 | |
| | | Sku08 | 523467 | 514 | 0.03 | |
| | | Sku09 | 456381 | 15 | 0.00 | |
| | | Sku10 | 719910 | 20 | 0.00 | |
| | | Sku11 | 686475 | 513 | 0.03 | |
| | | Sku12 | 1040685 | 26545 | 1.47 | |
| | | Sku13 | 921530 | 31 | 0.00 | |
| | | Sku14 | 797769 | 9 | 0.00 | |
| | | Sku15 | 1050702 | 44 | 0.00 | |
| | | Sku16 | 901691 | 30 | 0.00 | |
| | *S. mikatae* | Smi01 | 248052 | 2001 | 0.11 | 1.05 |
| | | Smi02 | 811203 | 42 | 0.00 | |
| | | Smi03 | 322525 | 36 | 0.00 | |
| | | Smi04 | 1495645 | 247 | 0.01 | |
| | | Smi05 | 599206 | 272 | 0.02 | |
| | | Smi06 | 848074 | 50 | 0.00 | |
| | | Smi07 | 663440 | 183 | 0.01 | |
| | | Smi08 | 542523 | 773 | 0.04 | |
| | | Smi09 | 462568 | 27 | 0.00 | |
| | | Smi10 | 734496 | 87 | 0.00 | |
| | | Smi11 | 687171 | 336 | 0.02 | |
| | | Smi12 | 1104771 | 12475 | 0.69 | |
| | | Smi13 | 932466 | 522 | 0.03 | |
| | | Smi14 | 753128 | 18 | 0.00 | |
| | | Smi15 | 1095438 | 1781 | 0.10 | |
| | | Smi16 | 804165 | 68 | 0.00 | |
| | *S. paradoxus* | Spa01 | 235665 | 216 | 0.01 | 0.72 |
| | | Spa02 | 835812 | 858 | 0.05 | |

| | | | | | | |
|---|---|---|---|---|---|---|
| | | Spa03 | 357685 | 882 | 0.05 | |
| | | Spa04 | 1477988 | 949 | 0.05 | |
| | | Spa05 | 565208 | 263 | 0.01 | |
| | | Spa06 | 296034 | 1071 | 0.06 | |
| | | Spa07 | 1119193 | 1914 | 0.11 | |
| | | Spa08 | 531267 | 1190 | 0.07 | |
| | | Spa09 | 443128 | 350 | 0.02 | |
| | | Spa10 | 743843 | 543 | 0.03 | |
| | | Spa11 | 690553 | 364 | 0.02 | |
| | | Spa12 | 1023116 | 536 | 0.03 | |
| | | Spa13 | 941477 | 149 | 0.01 | |
| | | Spa14 | 778926 | 1505 | 0.08 | |
| | | Spa15 | 1078405 | 1946 | 0.11 | |
| | | Spa16 | 903028 | 146 | 0.01 | |
| | *S. uvarum* | Suv01 | 204763 | 12 | 0.00 | 0.90 |
| | | Suv02 | 1292445 | 27 | 0.00 | |
| | | Suv03 | 301324 | 41 | 0.00 | |
| | | Suv04 | 991278 | 26 | 0.00 | |
| | | Suv05 | 564469 | 5 | 0.00 | |
| | | Suv06 | 258855 | 280 | 0.02 | |
| | | Suv07 | 1040537 | 231 | 0.01 | |
| | | Suv08 | 834793 | 97 | 0.01 | |
| | | Suv09 | 409318 | 53 | 0.00 | |
| | | Suv10 | 731806 | 11 | 0.00 | |
| | | Suv11 | 643462 | 221 | 0.01 | |
| | | Suv12 | 1035508 | 14557 | 0.81 | |
| | | Suv13 | 967206 | 303 | 0.02 | |
| | | Suv14 | 773076 | 16 | 0.00 | |
| | | Suv15 | 745522 | 351 | 0.02 | |
| | | Suv16 | 920918 | 43 | 0.00 | |
| Synthetic hybrid *S. mikatae* IFO1815 x *S. uvarum* CBS7001 (SRR7769316) | *S. cerevisiae* | Sce01 | 230218 | 90 | 0.01 | 1.08 |
| | | Sce02 | 813184 | 169 | 0.01 | |
| | | Sce03 | 316620 | 496 | 0.03 | |
| | | Sce04 | 1531933 | 948 | 0.05 | |
| | | Sce05 | 576874 | 113 | 0.01 | |
| | | Sce06 | 270161 | 41 | 0.00 | |
| | | Sce07 | 1090940 | 1094 | 0.06 | |
| | | Sce08 | 562643 | 102 | 0.01 | |
| | | Sce09 | 439888 | 72 | 0.00 | |
| | | Sce10 | 745751 | 108 | 0.01 | |
| | | Sce11 | 666816 | 495 | 0.03 | |
| | | Sce12 | 1078177 | 14204 | 0.82 | |
| | | Sce13 | 924431 | 107 | 0.01 | |

| | | | | | |
|---|---|---|---|---|---|
| | Sce14 | 784333 | 196 | 0.01 | |
| | Sce15 | 1091291 | 202 | 0.01 | |
| | Sce16 | 948066 | 126 | 0.01 | |
| *S. kudriavzevii* | Sku01 | 216969 | 118 | 0.01 | 1.32 |
| | Sku02 | 806230 | 233 | 0.01 | |
| | Sku03 | 355935 | 183 | 0.01 | |
| | Sku04 | 1470946 | 461 | 0.03 | |
| | Sku05 | 556891 | 114 | 0.01 | |
| | Sku06 | 288445 | 54 | 0.00 | |
| | Sku07 | 1119750 | 365 | 0.02 | |
| | Sku08 | 523467 | 101 | 0.01 | |
| | Sku09 | 456381 | 218 | 0.01 | |
| | Sku10 | 719910 | 201 | 0.01 | |
| | Sku11 | 686475 | 517 | 0.03 | |
| | Sku12 | 1040685 | 19094 | 1.11 | |
| | Sku13 | 921530 | 267 | 0.02 | |
| | Sku14 | 797769 | 260 | 0.02 | |
| | Sku15 | 1050702 | 323 | 0.02 | |
| | Sku16 | 901691 | 241 | 0.01 | |
| *S. mikatae* | Smi01 | 248052 | 24758 | 1.43 | **59.57** |
| | Smi02 | 811203 | 66486 | 3.85 | |
| | Smi03 | 322525 | 28431 | 1.65 | |
| | Smi04 | 1495645 | 115890 | 6.71 | |
| | Smi05 | 599206 | 50902 | 2.95 | |
| | Smi06 | 848074 | 68523 | 3.97 | |
| | Smi07 | 663440 | 56263 | 3.26 | |
| | Smi08 | 542523 | 47187 | 2.73 | |
| | Smi09 | 462568 | 41537 | 2.41 | |
| | Smi10 | 734496 | 61998 | 3.59 | |
| | Smi11 | 687171 | 57526 | 3.33 | |
| | Smi12 | 1104771 | 120357 | 6.97 | |
| | Smi13 | 932466 | 75496 | 4.37 | |
| | Smi14 | 753128 | 61180 | 3.54 | |
| | Smi15 | 1095438 | 85994 | 4.98 | |
| | Smi16 | 804165 | 65937 | 3.82 | |
| *S. paradoxus* | Spa01 | 235665 | 666 | 0.04 | 0.29 |
| | Spa02 | 835812 | 150 | 0.01 | |
| | Spa03 | 357685 | 142 | 0.01 | |
| | Spa04 | 1477988 | 439 | 0.03 | |
| | Spa05 | 565208 | 184 | 0.01 | |
| | Spa06 | 296034 | 41 | 0.00 | |
| | Spa07 | 1119193 | 575 | 0.03 | |
| | Spa08 | 531267 | 98 | 0.01 | |

| | | | | | | |
|---|---|---|---|---|---|---|
| | | Spa09 | 443128 | 146 | 0.01 | |
| | | Spa10 | 743843 | 237 | 0.01 | |
| | | Spa11 | 690553 | 147 | 0.01 | |
| | | Spa12 | 1023116 | 175 | 0.01 | |
| | | Spa13 | 941477 | 817 | 0.05 | |
| | | Spa14 | 778926 | 137 | 0.01 | |
| | | Spa15 | 1078405 | 959 | 0.06 | |
| | | Spa16 | 903028 | 153 | 0.01 | |
| | *S. uvarum* | Suv01 | 204763 | 10730 | 0.62 | **37.75** |
| | | Suv02 | 1292445 | 65799 | 3.81 | |
| | | Suv03 | 301324 | 16195 | 0.94 | |
| | | Suv04 | 991278 | 50305 | 2.91 | |
| | | Suv05 | 564469 | 28513 | 1.65 | |
| | | Suv06 | 258855 | 12663 | 0.73 | |
| | | Suv07 | 1040537 | 52349 | 3.03 | |
| | | Suv08 | 834793 | 40396 | 2.34 | |
| | | Suv09 | 409318 | 22519 | 1.30 | |
| | | Suv10 | 731806 | 39500 | 2.29 | |
| | | Suv11 | 643462 | 34692 | 2.01 | |
| | | Suv12 | 1035508 | 100937 | 5.85 | |
| | | Suv13 | 967206 | 48657 | 2.82 | |
| | | Suv14 | 773076 | 40280 | 2.33 | |
| | | Suv15 | 745522 | 40061 | 2.32 | |
| | | Suv16 | 920918 | 48164 | 2.79 | |
| Synthetic hybrid *S. cerevisiae* yHRW134 x *S. uvarum* CBS7001 (SRR7769317) | *S. cerevisiae* | Sce01 | 230218 | 20869 | 1.13 | **58.86** |
| | | Sce02 | 813184 | 68028 | 3.69 | |
| | | Sce03 | 316620 | 25427 | 1.38 | |
| | | Sce04 | 1531933 | 112866 | 6.12 | |
| | | Sce05 | 576874 | 52382 | 2.84 | |
| | | Sce06 | 270161 | 26934 | 1.46 | |
| | | Sce07 | 1090940 | 88315 | 4.79 | |
| | | Sce08 | 562643 | 50202 | 2.72 | |
| | | Sce09 | 439888 | 43197 | 2.34 | |
| | | Sce10 | 745751 | 63364 | 3.43 | |
| | | Sce11 | 666816 | 58192 | 3.15 | |
| | | Sce12 | 1078177 | 168335 | 9.12 | |
| | | Sce13 | 924431 | 78195 | 4.24 | |
| | | Sce14 | 784333 | 66327 | 3.59 | |
| | | Sce15 | 1091291 | 86691 | 4.70 | |
| | | Sce16 | 948066 | 76976 | 4.17 | |
| | *S. kudriavzevii* | Sku01 | 216969 | 161 | 0.01 | 1.67 |
| | | Sku02 | 806230 | 234 | 0.01 | |
| | | Sku03 | 355935 | 196 | 0.01 | |

| | | | | | |
|---|---|---|---|---|---|
| | Sku04 | 1470946 | 571 | 0.03 | |
| | Sku05 | 556891 | 138 | 0.01 | |
| | Sku06 | 288445 | 134 | 0.01 | |
| | Sku07 | 1119750 | 395 | 0.02 | |
| | Sku08 | 523467 | 162 | 0.01 | |
| | Sku09 | 456381 | 255 | 0.01 | |
| | Sku10 | 719910 | 237 | 0.01 | |
| | Sku11 | 686475 | 296 | 0.02 | |
| | Sku12 | 1040685 | 26883 | 1.46 | |
| | Sku13 | 921530 | 261 | 0.01 | |
| | Sku14 | 797769 | 324 | 0.02 | |
| | Sku15 | 1050702 | 344 | 0.02 | |
| | Sku16 | 901691 | 259 | 0.01 | |
| *S. mikatae* | Smi01 | 248052 | 336 | 0.02 | 0.88 |
| | Smi02 | 811203 | 125 | 0.01 | |
| | Smi03 | 322525 | 87 | 0.00 | |
| | Smi04 | 1495645 | 510 | 0.03 | |
| | Smi05 | 599206 | 207 | 0.01 | |
| | Smi06 | 848074 | 307 | 0.02 | |
| | Smi07 | 663440 | 267 | 0.01 | |
| | Smi08 | 542523 | 234 | 0.01 | |
| | Smi09 | 462568 | 142 | 0.01 | |
| | Smi10 | 734496 | 203 | 0.01 | |
| | Smi11 | 687171 | 156 | 0.01 | |
| | Smi12 | 1104771 | 12028 | 0.65 | |
| | Smi13 | 932466 | 130 | 0.01 | |
| | Smi14 | 753128 | 197 | 0.01 | |
| | Smi15 | 1095438 | 1039 | 0.06 | |
| | Smi16 | 804165 | 181 | 0.01 | |
| *S. paradoxus* | Spa01 | 235665 | 242 | 0.01 | 0.40 |
| | Spa02 | 835812 | 592 | 0.03 | |
| | Spa03 | 357685 | 384 | 0.02 | |
| | Spa04 | 1477988 | 473 | 0.03 | |
| | Spa05 | 565208 | 314 | 0.02 | |
| | Spa06 | 296034 | 645 | 0.03 | |
| | Spa07 | 1119193 | 556 | 0.03 | |
| | Spa08 | 531267 | 219 | 0.01 | |
| | Spa09 | 443128 | 303 | 0.02 | |
| | Spa10 | 743843 | 478 | 0.03 | |
| | Spa11 | 690553 | 206 | 0.01 | |
| | Spa12 | 1023116 | 476 | 0.03 | |
| | Spa13 | 941477 | 290 | 0.02 | |
| | Spa14 | 778926 | 949 | 0.05 | |

| | | | | | | |
|---|---|---|---|---|---|---|
| | | Spa15 | 1078405 | 948 | 0.05 | |
| | | Spa16 | 903028 | 263 | 0.01 | |
| | *S. uvarum* | Suv01 | 204763 | 12569 | 0.68 | **38.19** |
| | | Suv02 | 1292445 | 67719 | 3.67 | |
| | | Suv03 | 301324 | 19463 | 1.05 | |
| | | Suv04 | 991278 | 52985 | 2.87 | |
| | | Suv05 | 564469 | 31877 | 1.73 | |
| | | Suv06 | 258855 | 14535 | 0.79 | |
| | | Suv07 | 1040537 | 55147 | 2.99 | |
| | | Suv08 | 834793 | 45451 | 2.46 | |
| | | Suv09 | 409318 | 26556 | 1.44 | |
| | | Suv10 | 731806 | 45449 | 2.46 | |
| | | Suv11 | 643462 | 37848 | 2.05 | |
| | | Suv12 | 1035508 | 108310 | 5.87 | |
| | | Suv13 | 967206 | 50485 | 2.74 | |
| | | Suv14 | 773076 | 43422 | 2.35 | |
| | | Suv15 | 745522 | 42021 | 2.28 | |
| | | Suv16 | 920918 | 51053 | 2.77 | |
| Synthetic hybrid *S. kudriavzevii* ZP591 x *S. paradoxus* CBS432 (SRR7769319) | *S. cerevisiae* | Sce01 | 230218 | 44 | 0.00 | 1.64 |
| | | Sce02 | 813184 | 176 | 0.01 | |
| | | Sce03 | 316620 | 102 | 0.01 | |
| | | Sce04 | 1531933 | 1704 | 0.08 | |
| | | Sce05 | 576874 | 333 | 0.02 | |
| | | Sce06 | 270161 | 78 | 0.00 | |
| | | Sce07 | 1090940 | 956 | 0.05 | |
| | | Sce08 | 562643 | 190 | 0.01 | |
| | | Sce09 | 439888 | 390 | 0.02 | |
| | | Sce10 | 745751 | 131 | 0.01 | |
| | | Sce11 | 666816 | 876 | 0.04 | |
| | | Sce12 | 1078177 | 25779 | 1.28 | |
| | | Sce13 | 924431 | 1202 | 0.06 | |
| | | Sce14 | 784333 | 558 | 0.03 | |
| | | Sce15 | 1091291 | 252 | 0.01 | |
| | | Sce16 | 948066 | 241 | 0.01 | |
| | *S. kudriavzevii* | Sku01 | 216969 | 16576 | 0.82 | **48.32** |
| | | Sku02 | 806230 | 64346 | 3.20 | |
| | | Sku03 | 355935 | 28208 | 1.40 | |
| | | Sku04 | 1470946 | 111364 | 5.54 | |
| | | Sku05 | 556891 | 44365 | 2.21 | |
| | | Sku06 | 288445 | 23382 | 1.16 | |
| | | Sku07 | 1119750 | 84605 | 4.21 | |
| | | Sku08 | 523467 | 42194 | 2.10 | |
| | | Sku09 | 456381 | 35775 | 1.78 | |

| | | | | | |
|---|---|---|---|---|---|
| | Sku10 | 719910 | 55964 | 2.78 | |
| | Sku11 | 686475 | 53413 | 2.66 | |
| | Sku12 | 1040685 | 128974 | 6.41 | |
| | Sku13 | 921530 | 70864 | 3.52 | |
| | Sku14 | 797769 | 61821 | 3.07 | |
| | Sku15 | 1050702 | 80905 | 4.02 | |
| | Sku16 | 901691 | 68853 | 3.42 | |
| *S. mikatae* | Smi01 | 248052 | 3817 | 0.19 | 0.87 |
| | Smi02 | 811203 | 667 | 0.03 | |
| | Smi03 | 322525 | 56 | 0.00 | |
| | Smi04 | 1495645 | 419 | 0.02 | |
| | Smi05 | 599206 | 10 | 0.00 | |
| | Smi06 | 848074 | 48 | 0.00 | |
| | Smi07 | 663440 | 108 | 0.01 | |
| | Smi08 | 542523 | 610 | 0.03 | |
| | Smi09 | 462568 | 132 | 0.01 | |
| | Smi10 | 734496 | 32 | 0.00 | |
| | Smi11 | 687171 | 28 | 0.00 | |
| | Smi12 | 1104771 | 11120 | 0.55 | |
| | Smi13 | 932466 | 136 | 0.01 | |
| | Smi14 | 753128 | 22 | 0.00 | |
| | Smi15 | 1095438 | 176 | 0.01 | |
| | Smi16 | 804165 | 146 | 0.01 | |
| *S. paradoxus* | Spa01 | 235665 | 20472 | 1.02 | **48.50** |
| | Spa02 | 835812 | 67431 | 3.35 | |
| | Spa03 | 357685 | 29022 | 1.44 | |
| | Spa04 | 1477988 | 117133 | 5.83 | |
| | Spa05 | 565208 | 46724 | 2.32 | |
| | Spa06 | 296034 | 25646 | 1.28 | |
| | Spa07 | 1119193 | 88536 | 4.40 | |
| | Spa08 | 531267 | 44882 | 2.23 | |
| | Spa09 | 443128 | 38207 | 1.90 | |
| | Spa10 | 743843 | 60862 | 3.03 | |
| | Spa11 | 690553 | 56832 | 2.83 | |
| | Spa12 | 1023116 | 80391 | 4.00 | |
| | Spa13 | 941477 | 76103 | 3.78 | |
| | Spa14 | 778926 | 62623 | 3.11 | |
| | Spa15 | 1078405 | 88197 | 4.39 | |
| | Spa16 | 903028 | 72090 | 3.59 | |
| *S. uvarum* | Suv01 | 204763 | 129 | 0.01 | 0.67 |
| | Suv02 | 1292445 | 104 | 0.01 | |
| | Suv03 | 301324 | 38 | 0.00 | |
| | Suv04 | 991278 | 64 | 0.00 | |

| | | | | | | |
|---|---|---|---|---|---|---|
| | | Suv05 | 564469 | 7 | 0.00 | |
| | | Suv06 | 258855 | 7 | 0.00 | |
| | | Suv07 | 1040537 | 39 | 0.00 | |
| | | Suv08 | 834793 | 23 | 0.00 | |
| | | Suv09 | 409318 | 367 | 0.02 | |
| | | Suv10 | 731806 | 82 | 0.00 | |
| | | Suv11 | 643462 | 93 | 0.00 | |
| | | Suv12 | 1035508 | 11364 | 0.57 | |
| | | Suv13 | 967206 | 607 | 0.03 | |
| | | Suv14 | 773076 | 76 | 0.00 | |
| | | Suv15 | 745522 | 238 | 0.01 | |
| | | Suv16 | 920918 | 249 | 0.01 | |
| Synthetic hybrid *S. uvarum* CBS7001 x *S. mikatae* IFO1815 x *S. kudriavzevii* ZP591 (SRR7769320) | *S. cerevisiae* | Sce01 | 230218 | 89 | 0.01 | 1.11 |
| | | Sce02 | 813184 | 137 | 0.01 | |
| | | Sce03 | 316620 | 258 | 0.02 | |
| | | Sce04 | 1531933 | 948 | 0.06 | |
| | | Sce05 | 576874 | 134 | 0.01 | |
| | | Sce06 | 270161 | 25 | 0.00 | |
| | | Sce07 | 1090940 | 1087 | 0.06 | |
| | | Sce08 | 562643 | 103 | 0.01 | |
| | | Sce09 | 439888 | 210 | 0.01 | |
| | | Sce10 | 745751 | 84 | 0.00 | |
| | | Sce11 | 666816 | 412 | 0.02 | |
| | | Sce12 | 1078177 | 14898 | 0.88 | |
| | | Sce13 | 924431 | 72 | 0.00 | |
| | | Sce14 | 784333 | 135 | 0.01 | |
| | | Sce15 | 1091291 | 144 | 0.01 | |
| | | Sce16 | 948066 | 115 | 0.01 | |
| | *S. kudriavzevii* | Sku01 | 216969 | 6139 | 0.36 | **37.05** |
| | | Sku02 | 806230 | 40343 | 2.38 | |
| | | Sku03 | 355935 | 16793 | 0.99 | |
| | | Sku04 | 1470946 | 76282 | 4.50 | |
| | | Sku05 | 556891 | 29349 | 1.73 | |
| | | Sku06 | 288445 | 14727 | 0.87 | |
| | | Sku07 | 1119750 | 57236 | 3.38 | |
| | | Sku08 | 523467 | 26069 | 1.54 | |
| | | Sku09 | 456381 | 23393 | 1.38 | |
| | | Sku10 | 719910 | 36919 | 2.18 | |
| | | Sku11 | 686475 | 35639 | 2.10 | |
| | | Sku12 | 1040685 | 95795 | 5.66 | |
| | | Sku13 | 921530 | 46753 | 2.76 | |
| | | Sku14 | 797769 | 21047 | 1.24 | |
| | | Sku15 | 1050702 | 54902 | 3.24 | |

| | | | | | |
|---|---|---|---|---|---|
| | Sku16 | 901691 | 46033 | 2.72 | |
| *S. mikatae* | Smi01 | 248052 | 14861 | 0.88 | **37.77** |
| | Smi02 | 811203 | 45208 | 2.67 | |
| | Smi03 | 322525 | 12286 | 0.73 | |
| | Smi04 | 1495645 | 79733 | 4.71 | |
| | Smi05 | 599206 | 18044 | 1.07 | |
| | Smi06 | 848074 | 46671 | 2.76 | |
| | Smi07 | 663440 | 19449 | 1.15 | |
| | Smi08 | 542523 | 30136 | 1.78 | |
| | Smi09 | 462568 | 25885 | 1.53 | |
| | Smi10 | 734496 | 40884 | 2.41 | |
| | Smi11 | 687171 | 37412 | 2.21 | |
| | Smi12 | 1104771 | 95451 | 5.64 | |
| | Smi13 | 932466 | 49582 | 2.93 | |
| | Smi14 | 753128 | 41462 | 2.45 | |
| | Smi15 | 1095438 | 59648 | 3.52 | |
| | Smi16 | 804165 | 22841 | 1.35 | |
| *S. paradoxus* | Spa01 | 235665 | 681 | 0.04 | 0.27 |
| | Spa02 | 835812 | 118 | 0.01 | |
| | Spa03 | 357685 | 70 | 0.00 | |
| | Spa04 | 1477988 | 435 | 0.03 | |
| | Spa05 | 565208 | 108 | 0.01 | |
| | Spa06 | 296034 | 36 | 0.00 | |
| | Spa07 | 1119193 | 447 | 0.03 | |
| | Spa08 | 531267 | 85 | 0.01 | |
| | Spa09 | 443128 | 100 | 0.01 | |
| | Spa10 | 743843 | 168 | 0.01 | |
| | Spa11 | 690553 | 145 | 0.01 | |
| | Spa12 | 1023116 | 141 | 0.01 | |
| | Spa13 | 941477 | 852 | 0.05 | |
| | Spa14 | 778926 | 66 | 0.00 | |
| | Spa15 | 1078405 | 925 | 0.05 | |
| | Spa16 | 903028 | 113 | 0.01 | |
| *S. uvarum* | Suv01 | 204763 | 6259 | 0.37 | **23.80** |
| | Suv02 | 1292445 | 44956 | 2.65 | |
| | Suv03 | 301324 | 5092 | 0.30 | |
| | Suv04 | 991278 | 33499 | 1.98 | |
| | Suv05 | 564469 | 9572 | 0.57 | |
| | Suv06 | 258855 | 8165 | 0.48 | |
| | Suv07 | 1040537 | 36294 | 2.14 | |
| | Suv08 | 834793 | 26509 | 1.57 | |
| | Suv09 | 409318 | 14254 | 0.84 | |
| | Suv10 | 731806 | 24960 | 1.47 | |

| | | | |
|---|---|---|---|
| Suv11 | 643462 | 22183 | 1.31 |
| Suv12 | 1035508 | 68507 | 4.05 |
| Suv13 | 967206 | 29839 | 1.76 |
| Suv14 | 773076 | 13786 | 0.81 |
| Suv15 | 745522 | 26484 | 1.56 |
| Suv16 | 920918 | 32693 | 1.93 |

**Table S4.** Statistics of syntenic positions between master and secondary reference genomes of *Brachypodium* **(A)**, *Brassica* **(B)**, *Triticum-Aegilops* **(C)** and *Saccharomyces* **(D)** complexes calculated by the WGA algorithm. Columns 'Chr.' and 'Chr. length' correspond to the chromosome code and length of the master reference genome. Columns 'syntenic pos. (bp) per Chr', '% per Chr' and '% per complete genome' correspond to the number of syntenic positions between the master and secondary reference genomes, the percentage of syntenic positions per chromosome of the master genome and the total percentage of syntenic positions with respect to the master reference genome. The values in parentheses indicate the number of syntenic positions detected in coding regions (genes) and its percentage with respect to the total number of syntenic positions.

**(A)**

| **Master reference genome** | | **Secondary reference genome** | | **Outgroup genome** | |
|---|---|---|---|---|---|
| ***B. distachyon*** | | ***B. stacei*** | | ***O. sativa*** | |
| **Chr.** | **Chr. length (bp)** | **syntenic pos. (bp) per Chr** | **% per Chr** | **syntenic pos. (bp) per Chr** | **% per Chr** |
| **Bd1** | 75071545 | 22858427 | 30.4 | 2442206 | 3.3 |
| **Bd2** | 59130575 | 16992888 | 28.7 | 1687472 | 2.9 |
| **Bd3** | 59640145 | 18952267 | 31.8 | 1518257 | 2.5 |
| **Bd4** | 48594894 | 11936307 | 24.6 | 344244 | 0.7 |
| **Bd5** | 28630136 | 7858374 | 27.4 | 743565 | 2.6 |
| **TOTAL** | 271067295 | 78598263 (57777080; 73.5%) | 29.0 | 6735744 (6644302; 98.6%) | 2.5 |

**(B)**

| **Master reference genome** | | **Secondary reference genome** | |
|---|---|---|---|
| ***Br. oleracea*** | | ***Br. rapa*** | |
| **Chr.** | **Chr. length (bp)** | **syntenic pos. (bp) per Chr** | **% per Chr** |
| **Bro1** | 43764888 | 1964593 | 4.5 |
| **Bro2** | 52886895 | 3323914 | 6.3 |
| **Bro3** | 64984695 | 2893434 | 4.5 |
| **Bro4** | 53719093 | 2577970 | 4.8 |
| **Bro5** | 46902585 | 2482047 | 5.3 |
| **Bro6** | 39822476 | 1876709 | 4.7 |
| **Bro7** | 48366697 | 2219808 | 4.6 |
| **Bro8** | 41758685 | 2709966 | 6.5 |
| **Bro9** | 54679868 | 3592487 | 6.6 |
| **TOTAL** | 446885882 | 23640928 (11349305; 48.0%) | 5.3 |

**(C)**

| **Master reference genome** | | **Secondary reference genomes** | | | |
|---|---|---|---|---|---|
| ***T. urartu*** | | ***Ae. speltoides*** | | ***Ae. tauschii*** | |
| **Chr.** | **Chr. length (bp)** | **syntenic pos. (bp) per Chr** | **% per Chr** | **syntenic pos. (bp) per Chr** | **% per Chr** |
| **Tru1** | 584211917 | 16982754 | 2.9 | 19504893 | 3.3 |
| **Tru2** | 753719114 | 20827520 | 2.8 | 23848486 | 3.2 |
| **Tru3** | 747046674 | 19424293 | 2.6 | 22210636 | 3.0 |
| **Tru4** | 619584425 | 14274024 | 2.3 | 15642417 | 2.5 |
| **Tru5** | 661480603 | 19778118 | 3.0 | 22457558 | 3.4 |
| **Tru6** | 575865107 | 14943388 | 2.6 | 16865312 | 2.9 |
| **Tru7** | 719689837 | 19584304 | 2.7 | 21842890 | 3.0 |
| **TOTAL** | 4661597677 | 125814401 (59701973; 47.5%) | 2.7 | 142372192 (62256127; 43.7%) | 3.1 |

(D)

| Master reference genome | | Secondary reference genomes | | | | | | | |
|---|---|---|---|---|---|---|---|---|---|
| *S. cerevisiae* | | *S. kudriavzevii* | | *S. mikatae* | | *S. paradoxus* | | *S. uvarum* | |
| **Chr.** | **Chr. length (bp)** | **syntenic pos. (bp) per Chr** | **% per Chr** | **syntenic pos. (bp) per Chr** | **% per Chr** | **syntenic pos. (bp) per Chr** | **% per Chr** | **syntenic pos. (bp) per Chr** | **% per Chr** |
| **Sce01** | 230218 | 71644 | 31.1 | 81369 | 35.3 | 102705 | 44.6 | 62140 | 27.0 |
| **Sce02** | 813184 | 360282 | 44.3 | 377088 | 46.4 | 467023 | 57.4 | 274518 | 33.8 |
| **Sce03** | 316620 | 114834 | 36.3 | 127536 | 40.3 | 159477 | 50.4 | 97468 | 30.8 |
| **Sce04** | 1531933 | 622217 | 40.6 | 666933 | 43.5 | 820789 | 53.6 | 477619 | 31.2 |
| **Sce05** | 576874 | 226548 | 39.3 | 239494 | 41.5 | 310593 | 53.8 | 175940 | 30.5 |
| **Sce06** | 270161 | 100480 | 37.2 | 105801 | 39.2 | 136502 | 50.5 | 68051 | 25.2 |
| **Sce07** | 1090940 | 457207 | 41.9 | 476776 | 43.7 | 584428 | 53.6 | 345241 | 31.6 |
| **Sce08** | 562643 | 228948 | 40.7 | 242671 | 43.1 | 300977 | 53.5 | 184370 | 32.8 |
| **Sce09** | 439888 | 181637 | 41.3 | 194570 | 44.2 | 236723 | 53.8 | 138768 | 31.5 |
| **Sce10** | 745751 | 322518 | 43.2 | 333419 | 44.7 | 416125 | 55.8 | 248886 | 33.4 |
| **Sce11** | 666816 | 294312 | 44.1 | 302199 | 45.3 | 382009 | 57.3 | 222630 | 33.4 |
| **Sce12** | 1078177 | 452208 | 41.9 | 477258 | 44.3 | 586168 | 54.4 | 353484 | 32.8 |
| **Sce13** | 924431 | 409570 | 44.3 | 432568 | 46.8 | 528400 | 57.2 | 320229 | 34.6 |
| **Sce14** | 784333 | 351117 | 44.8 | 341222 | 43.5 | 452973 | 57.8 | 267743 | 34.1 |
| **Sce15** | 1091291 | 460123 | 42.2 | 476397 | 43.7 | 607368 | 55.7 | 344500 | 31.6 |
| **Sce16** | 948066 | 405712 | 42.8 | 418780 | 44.2 | 519596 | 54.8 | 301006 | 31.7 |
| **TOTAL** | 12071326 | 5059357 (4772846; 94.3%) | 41.9 | 5294081 (4917829; 92.9%) | 43.9 | 6611856 (5740596; 86.8%) | 54.8 | 3882593 (3736241; 96.2%) | 32.2 |

**Table S5.** Statistics of SHPs detected by VCF2SYNTENY according to the information of valid loci and syntenic positions provided by VCF2ALIGMENT and WGA algorithms. For each sample of the studied complexes *Brachypodium* **(A)**, *Brassica* **(B)**, *Triticum-Aegilops* **(C)** and *Saccharomyces* **(D)**, the number and percentage of syntenic valid nucleotides per subgenome are shown. The following abbreviations of the subgenomes are used: Bdis (*Brachypodium distachyon*), Bsta (*B. stacei*), Bro (*Brassica oleracea*), Brr (*Br. rapa*), Aes (*Aegilops speltoides*), Aet (*Ae. tauschii*), Tru (*Triticum urartu*), Sce (*Saccharomyces cerevisiae*), Sku (*S. kudriavzevii*), Smi (*S. mikatae*), Spa (S. *paradoxus*), and Suv (*S. uvarum*). The numbers of valid syntenic loci (no. of valid loci), syntenic polymorphic loci (no. of polymorphic loci) and the total alignment length in base pairs are given at the end of the table. Bold numbers indicate the predominant syntenic valid nucleotides respect to its subgenomes.

**(A)**

| **Sample** | **Subgenome** | **Single Homeologous Polymorphisms (SHPs)** | |
|---|---|---|---|
| | | **no.** | **%** |
| *B. distachyon* ABR2 | Bdis (D-subgenome) | 68972216 | **99.1** |
| | Bsta (S-subgenome) | 621850 | 0.9 |
| *B. distachyon* Bd21Control | Bdis (D-subgenome) | 69077363 | **99.4** |
| | Bsta (S-subgenome) | 444496 | 0.6 |
| *B. hybridum* Bhyb26 | Bdis (D-subgenome) | 67654320 | **49.6** |
| | Bsta (S-subgenome) | 68780216 | **50.4** |
| *B. hybridum* ABR113 | Bdis (D-subgenome) | 61116630 | **49.3** |
| | Bsta (S-subgenome) | 62844287 | **50.7** |
| *B. stacei* ABR114 | Bdis (D-subgenome) | 527613 | 0.8 |
| | Bsta (S-subgenome) | 67064626 | **99.2** |
| *B. stacei* TE4.3 | Bdis (D-subgenome) | 131526 | 0.2 |
| | Bsta (S-subgenome) | 69504597 | **99.8** |
| **no. of valid loci = 139535109**<br>**no. of polymorphic loci = 1781189**<br>**Total alignment length (bp)= 70409602** | | | |

**(B)**

| **Sample** | **Subgenome** | **Single Homeologous Polymorphisms (SHPs)** | |
|---|---|---|---|
| | | **no.** | **%** |
| *Br. oleracea* var. capitata | Bro (Co-subgenome) | 17468904 | **85.2** |
| | Brr (Ar-subgenome) | 3043231 | 14.8 |
| *Br. oleracea* var. oleracea | Bro (Co-subgenome) | 17836663 | **97.4** |
| | Brr (Ar-subgenome) | 477481 | 2.6 |
| *Br. rapa* var. chinensis | Bro (Co-subgenome) | 2349654 | 11.1 |
| | Brr (Ar-subgenome) | 18858516 | **88.9** |
| *Br. rapa* var. trilocularis | Bro (Co-subgenome) | 1677264 | 8.4 |
| | Brr (Ar-subgenome) | 18206830 | **91.6** |
| *Br. napus* R16GE06 | Bro (Co-subgenome) | 14720762 | **49.2** |
| | Brr (Ar-subgenome) | 15179472 | **50.8** |
| *Br. napus* R16GE38 | Bro (Co-subgenome) | 13443536 | **48.6** |
| | Brr (Ar-subgenome) | 14195202 | **51.4** |
| *Br. napus* R16GE39 | Bro (Co-subgenome) | 15187983 | **48.5** |
| | Brr (Ar-subgenome) | 16142241 | **51.5** |
| *Br. napus* R16GE44 | Bro (Co-subgenome) | 13354240 | **48.8** |
| | Brr (Ar-subgenome) | 13988525 | **51.2** |
| *Br. napus* R16GE45 | Bro (Co-subgenome) | 15405557 | **47.5** |
| | Brr (Ar-subgenome) | 17059950 | **52.5** |

**no. of valid loci = 37473519**
**no. of polymorphic loci = 574039**
**Total alignment length (bp) = 19628642**

**(C)**

| **Sample** | **Subgenome** | **Single Homeologous Polymorphisms (SHPs)** | |
|---|---|---|---|
| | | **no.** | **%** |
| *Ae. speltoides* | Aes (B-subgenome) | 73901947 | **65.4** |
| | Aet (D-subgenome) | 24463626 | 21.6 |
| | Tru (A-subgenome) | 14684228 | 13.0 |
| *Ae. tauschii* | Aes (B-subgenome) | 2482820 | 1.9 |
| | Aet (D-subgenome) | 122536578 | **95.6** |
| | Tru (A-subgenome) | 3111307 | 2.4 |
| *T. aestivum* | Aes (B-subgenome) | 55583639 | **19.7** |
| | Aet (D-subgenome) | 120095972 | **42.5** |
| | Tru (A-subgenome) | 106828499 | **37.8** |
| *T. turgidum* | Aes (B-subgenome) | 70165455 | **34.2** |
| | Aet (D-subgenome) | 25620160 | 12.5 |
| | Tru (A-subgenome) | 109092641 | **53.2** |
| *T. urartu* | Aes (B-subgenome) | 28938 | 0.3 |
| | Aet (D-subgenome) | 74493 | 0.8 |
| | Tru (A-subgenome) | 9621853 | **98.9** |

**no. of valid loci = 311156693**
**no. of polymorphic loci = 4043913**
**Total alignment length (bp) = 155473837**

**(D)**

| Sample | Subgenome | Single Homeologous Polymorphisms (SHPs) | |
|---|---|---|---|
| | | no. | % |
| *S. kudriavzevii* CLIB 1664 | Sce | 442 | 0.01 |
| | Sku | 4764398 | **99.96** |
| | Smi | 698 | 0.01 |
| | Spa | 347 | 0.01 |
| | Suv | 313 | 0.01 |
| *S. cerevisiae* BY4743 | Sce | 4126171 | **99.98** |
| | Sku | 0 | 0.00 |
| | Smi | 386 | 0.01 |
| | Spa | 310 | 0.01 |
| | Suv | 0 | 0.00 |
| *S. paradoxus* natural isolate - strain 95-7-1D: Sample 95-7-1D- 2 | Sce | 15301 | 0.36 |
| | Sku | 1100 | 0.03 |
| | Smi | 1944 | 0.05 |
| | Spa | 4189512 | **99.56** |
| | Suv | 0 | 0.00 |
| *S. uvarum* yHAB60 | Sce | 131 | 0.01 |
| | Sku | 0 | 0.00 |
| | Smi | 165 | 0.02 |
| | Spa | 308 | 0.03 |
| | Suv | 1000844 | **99.94** |
| *S. mikatae* yHAB328 | Sce | 1303 | 0.03 |
| | Sku | 0 | 0.00 |
| | Smi | 5042262 | **99.97** |
| | Spa | 0 | 0.00 |
| | Suv | 0 | 0.00 |
| Synthetic hybrid *S. cerevisiae* yHRW134 x *S. kudriavzevii* ZP591 x *S. mikatae* IFO1815 x *S. uvarum* CBS7001 | Sce | 651723 | **10.76** |
| | Sku | 2471474 | **40.80** |
| | Smi | 2746960 | **45.35** |
| | Spa | 638 | 0.01 |
| | Suv | 186392 | **3.08** |
| Synthetic hybrid *S. mikatae* IFO1815 x *S.* kudriavzevii ZP591 | Sce | 638 | 0.01 |
| | Sku | 3010461 | **47.69** |
| | Smi | 3301269 | **52.30** |
| | Spa | 0 | 0.00 |
| | Suv | 0 | 0.00 |
| Diploidized *S. cerevisiae* GLBRCY101 | Sce | 5978168 | **99.89** |
| | Sku | 0 | 0.00 |
| | Smi | 726 | 0.01 |
| | Spa | 5633 | 0.09 |
| | Suv | 0 | 0.00 |
| | Sce | 905 | 0.02 |

| | | | |
|---|---|---|---|
| Synthetic hybrid *S. mikatae* IFO1815 x *S. uvarum* CBS7001 | Sku | 0 | 0.00 |
| | Smi | 3550227 | **77.76** |
| | Spa | 19 | 0.00 |
| | Suv | 1014642 | **22.22** |
| Synthetic hybrid *S. cerevisiae* yHRW134 x *S. uvarum* CBS7001 | Sce | 4942381 | **81.35** |
| | Sku | 65 | 0.00 |
| | Smi | 457 | 0.01 |
| | Spa | 3282 | 0.05 |
| | Suv | 1129103 | **18.59** |
| Synthetic hybrid *S. kudriavzevii* ZP591 x *S. paradoxus* CBS432 | Sce | 1240 | 0.02 |
| | Sku | 3312625 | **44.14** |
| | Smi | 440 | 0.01 |
| | Spa | 4191241 | **55.84** |
| | Suv | 0 | 0.00 |
| Synthetic hybrid *S. uvarum* CBS7001 x *S. mikatae* IFO1815 x *S. kudriavzevii* ZP591 | Sce | 1251 | 0.04 |
| | Sku | 1512559 | **43.30** |
| | Smi | 1603479 | **45.91** |
| | Spa | 0 | 0.00 |
| | Suv | 375633 | **10.75** |
| **no. of valid loci = 21652756**<br>**no. of polymorphic loci = 163940**<br>**Total alignment length (bp) = 7104681** | | | |

**Table S6.** Site information of multiple sequence alignment of SHPs. Total and parsimony-informative sites of the complete alignment after filtering by *snp-sites* tool, and total sites, variable and parsimony-informative sites of the alignment, including only the subgenome sequences and the positions without missing data (Ns). All sites of the *Saccharomyces* subgenome sequences of the alignment contain at least one missing data and no data (nd) is shown for the statistics without missing data.

| | **Total alignment (with missing data-Ns)** | | **Allopolyploid subgenomes alignment (without missing data-Ns)** | | |
|---|---|---|---|---|---|
| | **Total sites** | **Parsimony-informative sites** | **Total sites** | **Variable sites** | **Parsimony-informative sites** |
| **BRACHYPODIUM** | 5958612 | 4378851 | 4617818 | 3975284 | 2889528 |
| **BRASSICA** | 980493 | 787595 | 98487 | 83299 | 77242 |
| **TRITICUM-AEGILOPS** | 8556827 | 6377583 | 1541991 | 1403364 | 937201 |
| **SACCHAROMYCES** | 1540088 | 1441544 | nd | nd | nd |

**Table S7.** Genome-wide average nucleotide identity (gwANI) for the MSA of SHPs used to infer the phylogeny for the *Brachypodium* **(A)**, *Brassica* **(B)**, *Triticum-Aegilops* **(C)**, and *Saccharomyces* **(D)** complexes. Same colors indicate identities between the same subgenomes of different allopolyploids/hybrids. Light colors indicate identities between subgenomes and their diploid parents.

**(A)**

| | *B. distachyon* ABR2 | *B. distachyon* Bd21Control | *B. hybridum* Bhyb26_Bdis | *B. hybridum* Bhyb26_Bsta | *B. hybridum* ABR113_Bdis | *B. hybridum* ABR113_Bsta | *B. stacei* ABR114 | *B. stacei* TE4.3 | *O. sativa* |
|---|---|---|---|---|---|---|---|---|---|
| *B. distachyon* ABR2 | 100.0 | 96.9 | 88.1 | 23.8 | 94.6 | 25.7 | 25.6 | 25.4 | 12.9 |
| *B. distachyon* Bd21Control | - | 100.0 | 88.3 | 24.0 | 95.4 | 25.9 | 25.8 | 25.6 | 13.0 |
| *B. hybridum* Bhyb26_Bdis | - | - | 100.0 | 23.0 | 88.4 | 24.8 | 24.7 | 24.5 | 12.4 |
| *B. hybridum* Bhyb26_Bsta | - | - | - | 100.0 | 24.2 | 87.3 | 87.1 | 87.0 | 10.6 |
| *B. hybridum* ABR113_Bdis | - | - | - | - | 100.0 | 26.0 | 25.9 | 25.7 | 12.7 |
| *B. hybridum* ABR113_Bsta | - | - | - | - | - | 100.0 | 97.6 | 97.2 | 11.0 |
| *B. stacei* ABR114 | - | - | - | - | - | - | 100.0 | 97.2 | 11.1 |
| *B. stacei* TE4.3 | - | - | - | - | - | - | - | 100.0 | 11.1 |
| *O. sativa* | - | - | - | - | - | - | - | - | 100.0 |

**(B)**

| | *Br. oleracea* capitata | *Br. oleracea* oleracea | *Br. rapa* chinensis | *Br. rapa* trilocularis | *Br. napus* R16GE06 Bro | *Br. napus* R16GE06 Brr | *Br. napus* R16GE38 Bro | *Br. napus* R16GE38 Brr | *Br. napus* R16GE39 Bro | *Br. napus* R16GE39 Brr | *Br. napus* R16GE44 Bro | *Br. napus* R16GE44 Brr | *Br. napus* R16GE45 Bro | *Br. napus* R16GE45 Brr |
|---|---|---|---|---|---|---|---|---|---|---|---|---|---|---|
| *Br. oleracea* var. capitata | 100.0 | 89.0 | 25.7 | 25.4 | 90.2 | 25.6 | 90.0 | 25.7 | 90.7 | 25.6 | 90.0 | 25.6 | 90.7 | 25.2 |
| *Br. oleracea* var. oleracea | - | 100.0 | 25.7 | 25.4 | 90.7 | 25.3 | 90.7 | 25.4 | 90.7 | 25.4 | 91.1 | 25.4 | 90.8 | 25.0 |
| *Br. rapa* subsp. chinesis | - | - | 100.0 | 82.1 | 25.8 | 84.3 | 25.7 | 85.0 | 25.7 | 83.8 | 25.9 | 83.9 | 25.6 | 86.0 |
| *Br. rapa* subsp. trilocularis | - | - | - | 100.0 | 25.5 | 79.8 | 25.4 | 80.1 | 25.4 | 79.3 | 25.7 | 79.4 | 25.3 | 80.7 |
| *Br. napus* R16GE06_Bro | - | - | - | - | 100.0 | 25.6 | 98.0 | 25.5 | 97.4 | 25.6 | 97.1 | 25.6 | 97.7 | 25.2 |
| *Br. napus* R16GE06_Brr | - | - | - | - | - | 100.0 | 25.3 | 90.5 | 25.5 | 93.3 | 25.6 | 92.3 | 25.4 | 93.7 |
| *Br. napus* R16GE38_Bro | - | - | - | - | - | - | 100.0 | 25.5 | 97.1 | 25.4 | 97.0 | 25.4 | 97.5 | 25.1 |
| *Br. napus* R16GE38_Brr | - | - | - | - | - | - | - | 100.0 | 25.6 | 91.4 | 25.7 | 92.1 | 25.5 | 94.1 |
| *Br. napus* R16GE39_Bro | - | - | - | - | - | - | - | - | 100.0 | 25.7 | 96.8 | 25.6 | 98.0 | 25.2 |
| *Br. napus* R16GE39_Brr | - | - | - | - | - | - | - | - | - | 100.0 | 25.8 | 92.3 | 25.5 | 94.0 |
| *Br. napus* R16GE44_Bro | - | - | - | - | - | - | - | - | - | - | 100.0 | 25.9 | 96.6 | 25.4 |
| *Br. napus* R16GE44_Brr | - | - | - | - | - | - | - | - | - | - | - | 100.0 | 25.5 | 94.8 |
| *Br. napus* R16GE45_Bro | - | - | - | - | - | - | - | - | - | - | - | - | 100.0 | 25.3 |
| *Br. napus* R16GE45_Brr | - | - | - | - | - | - | - | - | - | - | - | - | - | 100.0 |

**(C)**

| | *Ae. speltoides* | *Ae .tauschii* | *T. aestivum* Aes (B-subgenome) | *T. aestivum* Aet (D-subgenome) | *T. aestivum* Tru (A-subgenome) | *T. turgidum* Aes (B-subgenome) | *T .turgidum* Tru (A-subgenome) | *T. urartu* |
|---|---|---|---|---|---|---|---|---|
| *Ae. speltoides* | 100.0 | 38.9 | 69.5 | 38.9 | 32.1 | 68.0 | 32.1 | 34.4 |
| *Ae. tauschii* | - | 100.0 | 35.4 | 96.6 | 21.8 | 35.6 | 21.8 | 26.6 |
| *T. aestivum*_Aes (B-subgenome) | - | - | 100.0 | 35.7 | 29.2 | 95.2 | 29.1 | 31.4 |
| *T. aestivum*_Aet (D-subgenome) | - | - | - | 100.0 | 22.4 | 36.0 | 22.4 | 27.4 |
| *T. aestivum*_Tru (A-subgenome) | - | - | - | - | 100.0 | 29.3 | 96.1 | 90.9 |
| *T. turgidum*_Aes (B-subgenome) | - | - | - | - | - | 100.0 | 29.3 | 31.5 |
| *T. turgidum*_Tru (A-subgenome) | - | - | - | - | - | - | 100.0 | 90.8 |
| *T. urartu* | - | - | - | - | - | - | - | 100.0 |

**(D)**

| | ERR9706976 Sku | SRR13512344 Sce | SRR1868557 Spa | SRR7370090 Suv | SRR7370096 Smi | SRR7769310 Sce | SRR7769310 Sku | SRR7769310 Smi | SRR7769310 Suv | SRR7769312 Sku | SRR7769312 Smi | SRR7769313 Sce | SRR7769316 Smi | SRR7769316 Suv | SRR7769317 Sce | SRR7769317 Suv | SRR7769319 Sku | SRR7769319 Spa | SRR7769320 Sku | SRR7769320 Smi | SRR7769320 Suv |
|---|---|---|---|---|---|---|---|---|---|---|---|---|---|---|---|---|---|---|---|---|---|
| ERR9706976_Sku | 100.0 | 32.9 | 38.3 | 43.6 | 35.0 | 33.0 | 99.4 | 35.3 | 44.3 | 99.4 | 35.1 | 32.6 | 35.1 | 43.8 | 32.7 | 43.9 | 99.4 | 39.3 | 99.4 | 35.2 | 44.0 |
| SRR13512344_Sce | - | 100.0 | 55.1 | 36.6 | 38.2 | 98.4 | 32.9 | 38.5 | 37.4 | 33.0 | 38.4 | 98.3 | 38.4 | 37.0 | 98.3 | 36.9 | 33.0 | 57.3 | 33.1 | 38.2 | 37.0 |
| SRR1868557_Spa | - | - | 100.0 | 40.3 | 44.7 | 55.1 | 38.2 | 45.0 | 40.4 | 38.5 | 44.8 | 54.9 | 44.8 | 40.3 | 55.0 | 40.2 | 38.4 | 86.6 | 38.4 | 44.8 | 40.5 |
| SRR7370090_Suv | - | - | - | 100.0 | 38.0 | 36.6 | 43.7 | 38.0 | 99.1 | 43.6 | 38.1 | 36.6 | 38.0 | 99.3 | 36.6 | 99.3 | 43.7 | 40.9 | 43.7 | 38.1 | 99.3 |
| SRR7370096_Smi | - | - | - | - | 100.0 | 38.4 | 35.0 | 99.1 | 37.4 | 35.1 | 99.1 | 38.0 | 99.1 | 37.5 | 38.1 | 37.6 | 35.1 | 45.9 | 35.2 | 99.1 | 37.8 |
| SRR7769310_Sce | - | - | - | - | - | 100.0 | 33.2 | 38.9 | 36.1 | 32.8 | 38.7 | 100.0 | 38.6 | 36.5 | 100.0 | 36.7 | 33.1 | 57.2 | 33.0 | 38.1 | 37.9 |
| SRR7769310_Sku | - | - | - | - | - | - | 100.0 | 35.3 | 44.7 | 100.0 | 35.1 | 32.7 | 35.2 | 43.8 | 32.7 | 44.0 | 100.0 | 39.2 | 100.0 | 35.3 | 44.2 |
| SRR7769310_Smi | - | - | - | - | - | - | - | 100.0 | 37.4 | 35.5 | 100.0 | 38.3 | 100.0 | 37.6 | 38.3 | 37.7 | 35.4 | 46.3 | 35.6 | 100.0 | 37.8 |
| SRR7769310_Suv | - | - | - | - | - | - | - | - | 100.0 | 44.5 | 37.8 | 37.2 | 37.5 | 100.0 | 37.2 | 100.0 | 44.0 | 41.1 | 44.6 | 37.6 | 100.0 |
| SRR7769312_Sku | - | - | - | - | - | - | - | - | - | 100.0 | 35.3 | 32.7 | 35.3 | 43.8 | 32.7 | 43.9 | 100.0 | 39.4 | 100.0 | 35.3 | 44.0 |
| SRR7769312_Smi | - | - | - | - | - | - | - | - | - | - | 100.0 | 38.1 | 100.0 | 37.6 | 38.2 | 37.6 | 35.2 | 46.0 | 35.2 | 100.0 | 37.8 |
| SRR7769313_Sce | - | - | - | - | - | - | - | - | - | - | - | 100.0 | 38.2 | 36.9 | 100.0 | 36.9 | 32.8 | 57.0 | 32.8 | 38.0 | 37.0 |
| SRR7769316_Smi | - | - | - | - | - | - | - | - | - | - | - | - | 100.0 | 37.6 | 38.3 | 37.6 | 35.2 | 46.0 | 35.3 | 100.0 | 37.8 |
| SRR7769316_Suv | - | - | - | - | - | - | - | - | - | - | - | - | - | 100.0 | 36.8 | 100.0 | 43.8 | 41.0 | 44.0 | 38.0 | 100.0 |
| SRR7769317_Sce | - | - | - | - | - | - | - | - | - | - | - | - | - | - | 100.0 | 36.8 | 32.8 | 57.1 | 32.8 | 38.1 | 37.1 |
| SRR7769317_Suv | - | - | - | - | - | - | - | - | - | - | - | - | - | - | - | 100.0 | 43.9 | 40.9 | 44.1 | 37.7 | 100.0 |
| SRR7769319_Sku | - | - | - | - | - | - | - | - | - | - | - | - | - | - | - | - | 100.0 | 39.4 | 100.0 | 35.4 | 43.9 |
| SRR7769319_Spa | - | - | - | - | - | - | - | - | - | - | - | - | - | - | - | - | - | 100.0 | 39.3 | 46.0 | 41.3 |
| SRR7769320_Sku | - | - | - | - | - | - | - | - | - | - | - | - | - | - | - | - | - | - | 100.0 | 35.7 | 44.0 |
| SRR7769320_Smi | - | - | - | - | - | - | - | - | - | - | - | - | - | - | - | - | - | - | - | 100.0 | 38.4 |
| SRR7769320_Suv | - | - | - | - | - | - | - | - | - | - | - | - | - | - | - | - | - | - | - | - | 100.0 |

**Table S8.** Computational requirements for each of the datasets analyzed. **(A)** Computing time in seconds and required RAM in megabytes for each of the three executed algorithms (WGA, VCF2ALIGNMENT and VCF2SYNTENY) and for each of the analyzed databases (ID Analysis) and their corresponding input files. **(B)** The size in megabytes of each of the output files generated by each of the three algorithms executed (WGA, VCF2ALIGNMENT, and VCF2SYNTENY) and for each of the databases analyzed (ID Analysis). An asterisk in Table 8A indicates the input files corresponding to the reference genomes in FASTA format, from which the scaffold sequences were previously removed, and the chromosome headers were renamed. Two asterisks in Table 8B indicate that the VCF2SYNTENY output file sizes correspond to the alignment before removal of artifactual subgenomes.

**(A)**

| ID analysis | Input files | input file size (Mb) | WGA | | VCF2ALIGMENT | | VCF2SYNTENY | |
|---|---|---|---|---|---|---|---|---|
| | | | time (s) | RAM (Mb) | time (s) | RAM (Mb) | time (s) | RAM (Mb) |
| BRACHYPODIUM | Reference genome *Brachypodium distachyon* (FASTA)<br>Reference genome *Brachypodium stacei* (FASTA)<br>Outgroup genome *Oryza sativa* (FASTA)<br>VCF *Brachypodium* (bgzip) | 274<br>259<br>356<br>6454 | Bdis_Bsta: 1301<br>Bdis_Osat: 1170 | Bdis_Bsta: 251<br>Bdis_Osat: 967.3 | 7039 | 7720.7 | 16678 | 49474.7 |
| BRASSICA | Reference genome *Brassica oleracea** (FASTA)<br>Reference genome *Brassica rapa** (FASTA)<br>VCF *Brassica* (bgzip) | 499<br>357<br>16193 | 1931 | 716,4 | 6357 | 14675.3 | 10571 | 14089.3 |
| TRITICUM | Reference genome *Triticum urartu** (FASTA)<br>Reference genome *Aegilops tauschii** (FASTA)<br>Reference genome *Aegilops speltoides** (FASTA)<br>VCF *Triticum* (bgzip) | 4661<br>4021<br>3764<br>139028 | Tur_Aet: 8359<br>Tur_Aes: 8522 | Tur_Aet: 8115.2<br>Tur_Aes: 7642.1 | 93163 | 95795.7 | 148075 | 101193.2 |
| SACCHAROMYCES | Reference *Saccharomyces cerevisiae** (FASTA)<br>Reference *Saccharomyces kudriavzevii** (FASTA)<br>Reference *Saccharomyces mikatae** (FASTA)<br>Reference *Saccharomyces paradoxus** (FASTA)<br>Reference *Saccharomyces uvarum** (FASTA)<br>VCF *Saccharomyces* (bgzip) | 12<br>12<br>12<br>12<br>12<br>8823 | Sce_Sku: 193<br>Sce_Smi: 190<br>Sce_Spa: 210<br>Sce_Suv: 169 | Sce_Sku: 86.6<br>Sce_Smi: 88.7<br>Sce_Spa: 86.6<br>Sce_Suv: 89.7 | 2540 | 1669.9 | 785156 | 5944.9 |

**(B)**

| ID ANALYSIS | OUTPUT FILES (size in Mb) | | | | |
|---|---|---|---|---|---|
| | WGA | | VCF2ALIGNMENT | VCF2SYNTENY | |
| | FOLDER | BED | LOG | FASTA** | nohup_out |
| BRACHYPODIUM | Bdis_Bsta: 6606<br>Bdis_Osat: 2943 | Bdis_Bsta: 4366<br>Bdis_Osat: 369 | 13717<br>(compress .gz: 1304) | 915 | 3549 |
| BRASSICA | Bro_Brr: 4319 | Bro_Brr: 1335 | 18170<br>(compress .gz: 1739) | 353 | 1008 |
| TRITICUM | Tur_Aet: 23847<br>Tur_Aes: 22859 | Tur_Aet: 8758<br>Tur_Aes: 7734 | 223337<br>(compress .gz: 20279) | 2332 | 8850 |
| SACCHAROMYCES | Sce_Sku: 876<br>Sce_Smi: 886<br>Sce_Spa: 950<br>Sce_Suv: 809 | Sce_Sku: 253<br>Sce_Smi: 265<br>Sce_Spa: 330<br>Sce_Suv: 194 | 1386<br>(compress .gz: 139) | 426 | 456 |

**Table S9.** Statistics of the mappings of the *Brachypodium hybridum* Bhyb26 and ABR113 ecotypes against the concatenated diploid reference genomes composed of their diploid ancestors *B. distachyon* and *B. stacei*, plus the diploid perennial *B. sylvaticum*. Accession (species and ecotype), Chr length (reference chromosome length in base pairs), mapped read segments (number of reads mapped against the reference genome), % per chr (percentage of mappings per reference chromosome), % per (sub)genome (total percentage of mappings per reference genome). Bold numbers indicate predominant mappings.

| Accession | Reference genome and chromosome | | Chr length (bp) | mapped reads-segments | % per chr | % per (sub)genome |
|---|---|---|---|---|---|---|
| *Brachypodium hybridum* Bhyb26 | *B. distachyon* | Bd1 | 75071545 | 17433906 | 16.0 | |
| | | Bd2 | 59130575 | 11304488 | 10.4 | |
| | | Bd3 | 59640145 | 11946922 | 11.0 | **51.3** |
| | | Bd4 | 48594894 | 8990536 | 8.3 | |
| | | Bd5 | 28630136 | 6142869 | 5.6 | |
| | *B. stacei* | Chr01 | 31564145 | 6074353 | 5.6 | |
| | | Chr02 | 29889700 | 5628120 | 5.2 | |
| | | Chr03 | 27658146 | 5730048 | 5.3 | |
| | | Chr04 | 27050098 | 4965602 | 4.6 | |
| | | Chr05 | 25022241 | 4518145 | 4.2 | |
| | | Chr06 | 24930558 | 4695311 | 4.3 | **44.3** |
| | | Chr07 | 22499862 | 4314412 | 4.0 | |
| | | Chr08 | 22836500 | 4189091 | 3.8 | |
| | | Chr09 | 22451604 | 4184291 | 3.8 | |
| | | Chr10 | 22136647 | 3980102 | 3.7 | |
| | *B. sylvaticum* | chr1 | 52668086 | 774721 | 0.7 | |
| | | chr2 | 42843931 | 551673 | 0.5 | |
| | | chr3 | 30155131 | 438419 | 0.4 | |
| | | chr4 | 38764466 | 531075 | 0.5 | |
| | | chr5 | 48036586 | 752409 | 0.7 | 4.4 |
| | | chr6 | 23360316 | 309317 | 0.3 | |
| | | chr7 | 22377421 | 320709 | 0.3 | |
| | | chr8 | 28316078 | 624180 | 0.6 | |
| | | chr9 | 31930771 | 464901 | 0.4 | |
| *Brachypodium hybridum* ABR113 | *B. distachyon* | Bd1 | 75071545 | 15653973 | 15.4 | |
| | | Bd2 | 59130575 | 10123143 | 10.0 | |
| | | Bd3 | 59640145 | 10545769 | 10.4 | **48.8** |
| | | Bd4 | 48594894 | 7962488 | 7.9 | |
| | | Bd5 | 28630136 | 5145933 | 5.1 | |
| | *B. stacei* | Chr01 | 31564145 | 5873836 | 5.8 | |
| | | Chr02 | 29889700 | 5441000 | 5.4 | |
| | | Chr03 | 27658146 | 6879575 | 6.8 | **48.9** |
| | | Chr04 | 27050098 | 4647262 | 4.6 | |
| | | Chr05 | 25022241 | 4750431 | 4.7 | |

| | | | | | | |
|---|---|---|---|---|---|---|
| | | Chr06 | 24930558 | 4858254 | 4.8 | |
| | | Chr07 | 22499862 | 3940615 | 3.9 | |
| | | Chr08 | 22836500 | 4147458 | 4.1 | |
| | | Chr09 | 22451604 | 3694168 | 3.6 | |
| | | Chr10 | 22136647 | 5399378 | 5.3 | |
| | *B. sylvaticum* | chr1 | 52668086 | 299796 | 0.3 | 2.3 |
| | | chr2 | 42843931 | 244241 | 0.2 | |
| | | chr3 | 30155131 | 382532 | 0.4 | |
| | | chr4 | 38764466 | 206792 | 0.2 | |
| | | chr5 | 48036586 | 286211 | 0.3 | |
| | | chr6 | 23360316 | 117195 | 0.1 | |
| | | chr7 | 22377421 | 125449 | 0.1 | |
| | | chr8 | 28316078 | 470502 | 0.5 | |
| | | chr9 | 31930771 | 199178 | 0.2 | |

**Table S10.** Statistics of the mappings of *B. distachyon* ecotypes (ABR2, Bd21 and BdTR8i from Spanish, Turkish and EDF clade/groups) against the concatenated diploid reference genomes composed of **(A)** *B. distachyon* ecotypes Tek2 and ABR3 from EDF clade and Spanish group and **(B)** the previous reference *B. distachyon* ecotypes plus BdTR12c from Turkish group. Accession (species, ecotype, and phylogenetic clade/group), Chr length (reference chromosome length in base pairs), mapped read-segments (number of reads mapped against the reference genome), % per chr (percentage of mappings per reference chromosome), % per (sub)genome (total percentage of mappings per reference genome). Bold numbers indicate dominant mappings.

**(A)**

| **Accession** | **Reference genome and chromosome** | | **Chr length (bp)** | **mapped reads-segments** | **% per chr** | **% per (sub)genome** |
|---|---|---|---|---|---|---|
| *Brachypodium distachyon* ABR2 (Spanish group) | *B. distachyon* Tek2 (EDF clade) pangenome ref | Tek2_pseudomolecule_1 | 52350941 | 8088047 | 4.2 | 19.4 |
| | | Tek2_pseudomolecule_2 | 39888752 | 7109507 | 3.7 | |
| | | Tek2_pseudomolecule_3 | 29447999 | 5827829 | 3.0 | |
| | | Tek2_pseudomolecule_4 | 38739049 | 6347099 | 3.3 | |
| | | Tek2_pseudomolecule_5 | 15982489 | 2719068 | 1.4 | |
| | | Tek2_pseudomolecule_6 | 7303 | 1271 | 0.0 | |
| | | Tek2_pseudomolecule_7 | 64063095 | 7522468 | 3.9 | |
| | *B. distachyon* ABR3 (+S group) pangenome ref | ABR3_pseudomolecule_1 | 63574859 | 36731179 | 19.0 | **80.6** |
| | | ABR3_pseudomolecule_2 | 51228775 | 26311775 | 13.6 | |
| | | ABR3_pseudomolecule_3 | 47197441 | 25014291 | 12.9 | |
| | | ABR3_pseudomolecule_4 | 39262724 | 19995236 | 10.3 | |
| | | ABR3_pseudomolecule_5 | 20286418 | 10657071 | 5.5 | |
| | | ABR3_pseudomolecule_6 | 18019 | 9904 | 0.0 | |
| | | ABR3_pseudomolecule_7 | 8363 | 1265 | 0.0 | |
| | | ABR3_pseudomolecule_8 | 50187461 | 37235237 | 19.2 | |
| | | Tek2_pseudomolecule_1 | 52350941 | 13308994 | 7.6 | 35.6 |

| | | | | | | |
|---|---|---|---|---|---|---|
| *Brachypodium distachyon* Bd21 (Turkish group) | *B. distachyon* Tek2 (EDF clade) pangenome ref | Tek2_pseudomolecule_2 | 39888752 | 10346369 | 5.9 | |
| | | Tek2_pseudomolecule_3 | 29447999 | 7865915 | 4.5 | |
| | | Tek2_pseudomolecule_4 | 38739049 | 10585720 | 6.0 | |
| | | Tek2_pseudomolecule_5 | 15982489 | 4641638 | 2.7 | |
| | | Tek2_pseudomolecule_6 | 7303 | 2713 | 0.0 | |
| | | Tek2_pseudomolecule_7 | 64063095 | 15500855 | 8.9 | |
| | *B. distachyon* ABR3 (+S group) pangenome ref | ABR3_pseudomolecule_1 | 63574859 | 26431302 | 15.1 | **64.4** |
| | | ABR3_pseudomolecule_2 | 51228775 | 18546470 | 10.6 | |
| | | ABR3_pseudomolecule_3 | 47197441 | 16730527 | 9.6 | |
| | | ABR3_pseudomolecule_4 | 39262724 | 14729485 | 8.4 | |
| | | ABR3_pseudomolecule_5 | 20286418 | 6848324 | 3.9 | |
| | | ABR3_pseudomolecule_6 | 18019 | 6628 | 0.0 | |
| | | ABR3_pseudomolecule_7 | 8363 | 3841 | 0.0 | |
| | | ABR3_pseudomolecule_8 | 50187461 | 29423170 | 16.8 | |
| *Brachypodium distachyon* BdTR8i (EDF clade) | *B. distachyon* Tek2 (EDF clade) pangenome ref | Tek2_pseudomolecule_1 | 52350941 | 9576785 | 13.9 | **62.4** |
| | | Tek2_pseudomolecule_2 | 39888752 | 7215271 | 10.5 | |
| | | Tek2_pseudomolecule_3 | 29447999 | 5433178 | 7.9 | |
| | | Tek2_pseudomolecule_4 | 38739049 | 7158456 | 10.4 | |
| | | Tek2_pseudomolecule_5 | 15982489 | 2864869 | 4.2 | |
| | | Tek2_pseudomolecule_6 | 7303 | 1200 | 0.0 | |
| | | Tek2_pseudomolecule_7 | 64063095 | 10793549 | 15.7 | |
| | *B. distachyon* ABR3 (+S group) pangenome ref | ABR3_pseudomolecule_1 | 63574859 | 5138361 | 7.5 | 37.6 |
| | | ABR3_pseudomolecule_2 | 51228775 | 3004437 | 4.4 | |
| | | ABR3_pseudomolecule_3 | 47197441 | 2581547 | 3.7 | |
| | | ABR3_pseudomolecule_4 | 39262724 | 2646662 | 3.8 | |
| | | ABR3_pseudomolecule_5 | 20286418 | 1363447 | 2.0 | |
| | | ABR3_pseudomolecule_6 | 18019 | 3603 | 0.0 | |
| | | ABR3_pseudomolecule_7 | 8363 | 869 | 0.0 | |
| | | ABR3_pseudomolecule_8 | 50187461 | 11154086 | 16.2 | |

**(B)**

| Accession | Reference genome and chromosome | | Chr length (bp) | mapped reads-segments | % per chr | % per (sub)genome |
|---|---|---|---|---|---|---|
| *Brachypodium distachyon* ABR2 (Spanish group) | *B. distachyon* Tek2 (EDF clade) pangenome ref | Tek2_pseudomolecule_1 | 52350941 | 5198268 | 2.7 | 12.7 |
| | | Tek2_pseudomolecule_2 | 39888752 | 4658563 | 2.4 | |
| | | Tek2_pseudomolecule_3 | 29447999 | 3824351 | 2.0 | |
| | | Tek2_pseudomolecule_4 | 38739049 | 4291277 | 2.2 | |
| | | Tek2_pseudomolecule_5 | 15982489 | 1751569 | 0.9 | |
| | | Tek2_pseudomolecule_6 | 7303 | 887 | 0.0 | |
| | | Tek2_pseudomolecule_7 | 64063095 | 4966052 | 2.6 | |
| | *B. distachyon* ABR3 (+S | ABR3_pseudomolecule_1 | 63574859 | 24877624 | 12.830 | **58.3** |
| | | ABR3_pseudomolecule_2 | 51228775 | 18191128 | 9.4 | |
| | | ABR3_pseudomolecule_3 | 47197441 | 18145583 | 9.4 | |

| | | | | | | |
|---|---|---|---|---|---|---|
| | group) pangenome ref | ABR3_pseudomolecule_4 | 39262724 | 14599197 | 7.5 | |
| | | ABR3_pseudomolecule_5 | 20286418 | 7702942 | 4.0 | |
| | | ABR3_pseudomolecule_6 | 18019 | 9833 | 0.0 | |
| | | ABR3_pseudomolecule_7 | 8363 | 560 | 0.0 | |
| | | ABR3_pseudomolecule_8 | 50187461 | 29438836 | 15.2 | |
| | *B. distachyon* BdTR12c (+T group) pangenome ref | BdTR12c_pseudomolecule_1 | 51572474 | 12072632 | 6.2 | 29.0 |
| | | BdTR12c_pseudomolecule_2 | 39284944 | 9137261 | 4.7 | |
| | | BdTR12c_pseudomolecule_3 | 29274147 | 6383187 | 3.3 | |
| | | BdTR12c_pseudomolecule_4 | 36082448 | 7843029 | 4.0 | |
| | | BdTR12c_pseudomolecule_5 | 15810652 | 3373739 | 1.7 | |
| | | BdTR12c_pseudomolecule_6 | 55150 | 13570 | 0.0 | |
| | | BdTR12c_pseudomolecule_7 | 9663 | 3451 | 0.0 | |
| | | BdTR12c_pseudomolecule_8 | 81201328 | 17415846 | 9.0 | |
| *Brachypodium distachyon* Bd21 (Turkish group) | *B. distachyon* Tek2 (EDF clade) pangenome ref | Tek2_pseudomolecule_1 | 52350941 | 9130623 | 5.2 | 24.1 |
| | | Tek2_pseudomolecule_2 | 39888752 | 6788646 | 3.9 | |
| | | Tek2_pseudomolecule_3 | 29447999 | 5001011 | 2.9 | |
| | | Tek2_pseudomolecule_4 | 38739049 | 6935297 | 4.0 | |
| | | Tek2_pseudomolecule_5 | 15982489 | 3154891 | 1.8 | |
| | | Tek2_pseudomolecule_6 | 7303 | 1460 | 0.0 | |
| | | Tek2_pseudomolecule_7 | 64063095 | 11306269 | 6.4 | |
| | *B. distachyon* ABR3 (+S group) pangenome ref | ABR3_pseudomolecule_1 | 63574859 | 15595777 | 8.9 | **40.6** |
| | | ABR3_pseudomolecule_2 | 51228775 | 11191925 | 6.4 | |
| | | ABR3_pseudomolecule_3 | 47197441 | 10090967 | 5.8 | |
| | | ABR3_pseudomolecule_4 | 39262724 | 9195250 | 5.2 | |
| | | ABR3_pseudomolecule_5 | 20286418 | 4262595 | 2.4 | |
| | | ABR3_pseudomolecule_6 | 18019 | 3076 | 0.0 | |
| | | ABR3_pseudomolecule_7 | 8363 | 3385 | 0.0 | |
| | | ABR3_pseudomolecule_8 | 50187461 | 20870583 | 11.9 | |
| | *B. distachyon* BdTR12c (+T group) pangenome ref | BdTR12c_pseudomolecule_1 | 51572474 | 12147149 | 6.9 | 35.3 |
| | | BdTR12c_pseudomolecule_2 | 39284944 | 9387815 | 5.4 | |
| | | BdTR12c_pseudomolecule_3 | 29274147 | 7156892 | 4.1 | |
| | | BdTR12c_pseudomolecule_4 | 36082448 | 8675083 | 4.9 | |
| | | BdTR12c_pseudomolecule_5 | 15810652 | 3540419 | 2.0 | |
| | | BdTR12c_pseudomolecule_6 | 55150 | 15313 | 0.0 | |
| | | BdTR12c_pseudomolecule_7 | 9663 | 1232 | 0.0 | |
| | | BdTR12c_pseudomolecule_8 | 81201328 | 20984955 | 12.0 | |
| *Brachypodium distachyon* BdTR8i (EDF clade) | *B. distachyon* Tek2 (EDF clade) pangenome ref | Tek2_pseudomolecule_1 | 52350941 | 8440468 | 12.2 | **55.0** |
| | | Tek2_pseudomolecule_2 | 39888752 | 6336165 | 9.2 | |
| | | Tek2_pseudomolecule_3 | 29447999 | 4721257 | 6.8 | |
| | | Tek2_pseudomolecule_4 | 38739049 | 6188881 | 9.0 | |
| | | Tek2_pseudomolecule_5 | 15982489 | 2486636 | 3.6 | |
| | | Tek2_pseudomolecule_6 | 7303 | 917 | 0.0 | |
| | | Tek2_pseudomolecule_7 | 64063095 | 9719436 | 14.1 | |
| | *B. distachyon* | ABR3_pseudomolecule_1 | 63574859 | 3438591 | 5.0 | 27.5 |
| | | ABR3_pseudomolecule_2 | 51228775 | 2268894 | 3.3 | |

| | | | | | |
|---|---|---|---|---|---|
| ABR3 (+S group) pangenome ref | ABR3_pseudomolecule_3 | 47197441 | 1879991 | 2.7 | |
| | ABR3_pseudomolecule_4 | 39262724 | 2032610 | 3.0 | |
| | ABR3_pseudomolecule_5 | 20286418 | 1031009 | 1.5 | |
| | ABR3_pseudomolecule_6 | 18019 | 3521 | 0.0 | |
| | ABR3_pseudomolecule_7 | 8363 | 753 | 0.0 | |
| | ABR3_pseudomolecule_8 | 50187461 | 8299183 | 12.0 | |
| *B. distachyon* BdTR12c (+T group) pangenome ref | BdTR12c_pseudomolecule_1 | 51572474 | 1776369 | 2.6 | 17.5 |
| | BdTR12c_pseudomolecule_2 | 39284944 | 1323417 | 1.9 | |
| | BdTR12c_pseudomolecule_3 | 29274147 | 1026610 | 1.5 | |
| | BdTR12c_pseudomolecule_4 | 36082448 | 1362162 | 2.0 | |
| | BdTR12c_pseudomolecule_5 | 15810652 | 542105 | 0.8 | |
| | BdTR12c_pseudomolecule_6 | 55150 | 1451 | 0.0 | |
| | BdTR12c_pseudomolecule_7 | 9663 | 418 | 0.0 | |
| | BdTR12c_pseudomolecule_8 | 81201328 | 6039763 | 8.8 | |
